## Supplementary Material for "HaploCart: Human mtDNA Haplogroup Classification Using a Pangenomic Reference Graph"

Nicola Vogel[1]

Peter Wad Sackett[1]

Shyam Gopalakrishnan[2]

Gabriel Renaud[1]

[1]Department of Health Technology, Section for Bioinformatics, Technical University of Denmark, Kongens Lyngby, Denmark
[2] Center for Evolutionary Hologenomics, GLOBE Institute, Faculty of Health and Medical Sciences, Copenhagen, Denmark

October 25, 2022

### 1 Methods

#### 1.1 Mappability Scores from GenMap

To obtain site-specific mappability scores, chromosomes 1 to 22 plus the X chromosome from human genome build CHM13v1.1 were concatenated with the Y chromosome and the chrY_KI270740v1_random scaffold from hg38 and the rCRS to form a new reference genome. This reference was indexed using GenMap v1.3.0 with default parameters and subsequently mapped with the map command from GenMap with parameters -K 30 -E 2 -t -w -bg -fl[14]. The resultant per-base mappability scores are then used to calibrate the quantity $P(\neg M)$, i.e. the probability that a read is mismapped or originates from a NuMT, as described in the Inference section of the main text.

#### 1.2 Incorporation of Artificial NuMT Reads

Our database of NuMT reads was taken from the UCSC Genome Browser, the data being "provided by Francesco Maria Calabrese, Domenico Simone and Marcella Attimonelli from the Department of Biochemistry and Molecular Biology "Ernesto Quagliariello (University of Bari, Italy)". These sequences were "[o]btained by running Blast2seq (program: BlastN) between each chromosome of of the Human Genome hg18 build and the human mitochondrial reference sequence (rCRS, AC: NC_012920), fixing the e-value threshold to 1e-03." Mapping coordinates were lifted over from hg18 to hg19 by using the Lift-Over part of the Galaxy software suite. Assembly of the HSPs was performed "with spreadsheet interpolation and manual inspection. BED format is used for the first three annotation tracks, while for the last one the SAM/BAM format is preferred"[8, 16, 5].

Each included NuMT read was selected uniformly from this compiled database of NuMT sequences of variable length using a custom script (`add_numt.sh` in the data repository - see the "Availability of Data and Materials" section of the main text).

#### 1.3 PCR Duplicate Removal

`HaploCart` removes duplicate PCR reads from a GAM file before performing inference. This step is necessary because the inference algorithm assumes independence among reads, and if this assumption does not hold we may over-penalize errors in the read sequences.

We have implemented a subcommand within the `vgan` package for duplicate removal called `vgan duprm`. `HaploCart` calls this subcommand internally but it is also available for use as a standalone program. The subcommand requires that the GAM file has already been sorted with respect to coordinates using `vg gamsort`. Here we describe the procedure used. First, a few definitions from `vg` parlance are required.

Each node in a `vg` graph is identified by a unique node identifier called a *node ID*. Each node is associated with a unique node sequence which can be read in one of two *orientations* (forward or reverse). If the orientation is reverse the sequence must be read as a reverse complement. The index of a given base in a node sequence from the standpoint of a given orientation is termed the *offset* of that base. When `vg giraffe` maps a read to the graph, it is possible (and very likely) for the read to be mapped to multiple nodes. Each alignment of a read subsequence to a node sequence in a given orientation is termed a *mapping*, and the totality of all mappings of a read to connected nodes is called a *path*[21].

Our PCR duplicate removal procedure runs as follows. First we record the starting coordinates (node id and offset) of the first mapping to a node of the first read in the sorted GAM file. Call this read the *candidate*. Then for each subsequent read (call it the *putative duplicate*) we check whether the starting coordinates between the candidate and putative duplicate are the same. For paired-end reads we also require that the ending coordinates (i.e. the node ID and offset of the final mapping) are the same between the candidate and putative duplicate.

After this is done, the second read in the sorted GAM file becomes the new candidate, the third read becomes the putative duplicate, and we repeat the loop described above. This procedure continues until every pair of reads in the GAM file has been checked. Reads which have been marked as duplicates are then removed.

#### 1.4 Test Data and Nucleotide Frequencies

The table of the FASTA IDs is found in Table S1, information about the 1000 Genomes samples used is found in Table S2 and nucleotide background frequency is found in Table S3.

#### `HaploGrouper` Results on Empirical Consensus FASTA Sequences

`HaploGrouper` takes as input VCF files. To obtain `HaploGrouper` predictions on consensus FASTA sequences we first perform a pairwise alignment with the rCRS using `minimap2`[**li2018minimap2**]. As with the paired-end FASTQ dataset we then call variants with `bcftools` under a haploid model (options: –ploidy 1 -c -v -Oz).

#### Software Versions

For all experiments we used `HaploCheck` version 1.3.2 and `HaploGrep2` version 2.4.0. All experiments were performed using `VG` version `Obolo`. All `samtools` commands used version 1.13. Our Snakemake version was 5.10.0. Our `ART` version was 2.5.8. Our `ODGI` version was v0.6.3-52-g0d7f950 ("Pulizia"). We downloaded `HaploGrouper` from the Gitlab repository `https://gitlab.com/bio_anth_decode/haploGrouper` with commit SHA e4c0a0e0. Our `bcftools` version was 1.10.2 using `htslib` version 1.10.2-3. Our `minimap2` version was 2.24-r1122.

Supplementary Table 1: Ground truth haplogroups of samples using in the downsampling and masking experiments. Haplogroups were determined by running `HaploGrep2 classify` with the `--phylotree 17` flag. Sample JX154035 was excluded from downsampling experiments as described but is still shown in the posterior plots.

| Haplogroup | NCBI Accession |
|---|---|
| HV4b | KP340180 |
| L0a1a1 | MK295855 |
| L1c4b | MN894773 |
| B2b3a | MW057682 |
| H2a2a1 | JX154035 [1] |
| J2a1a1a1 | MZ190830 |
| L2a1a3c | KR135866 |
| T2e1a1b1 | JN828512 |
| L2a1j | KR135846 |
| L1c2b | MN894780 [2] |
| Q1 | MN849793 |
| C1b | MN894713 |
| Z1a | MG660559 |
| D1 | KP172430 |
| A2 | MZ387838 |
| Y1b | GU123044 |
| P | MN849673 |
| L3b1 | KT819256 |
| U2e1b1 | KT698031 |
| F1a1 | MH553920 |
| I2b | MN516596 |
| S1a | DQ404440 |
| V3 | MN516629 |
| X3a | JQ245804 |
| E1a1b | EF061151 |

[1] the sample was removed due to being the same haplogroup as the rCRS, see main methods.
[2] the sample was removed due to unstable mutations, see main methods.

Supplementary Table 2: Ground truth haplogroups in empirical paired-end FASTQ experiments. Samples were chosen such that each sample comes from a distinct sequencing center and so that the dataset reflects a high degree of geographic diversity. Africa is intentionally over-represented owing to the fact that African populations harbor the majority of the mitodiversity of humans.

| Haplogroup | NCBI Accession | Population |
|---|---|---|
| D6a1a | HG00473 | Southern Han Chinese (CHS) |
| K1a4a1h | HG01051 | Puerto Rican in Puerto Rico (PUR) |
| L3e4a | HG02666 | Gambian in Western Division, The Gambia - Mandinka (GWD) |
| L1b1a3 | HG03112 | Esan in Nigeria (ESN) |
| L0a1a3 | NA18510 | Yoruba in Ibadan, Nigeria (YRI) |
| L3b1a1a | NA19036 | Luhya in Webuye, Kenya (LWK) |
| D1h1 | NA19661 | Mexican Ancestry in Los Angeles, California (MXL) |
| J2a1a1e | NA20518 | Toscani in Italy (TSI) |

Supplementary Table 3: Background frequencies of nucleobases in the graph. Values were computed with the `ODGI stats -S` command. Note that these statistics are on the graph itself, not the embedded haplogroup, meaning that each base is counted once even if it is traversed by multiple embedded paths.

| Base | Background frequency |
|---|---|
| A | 0.27532 |
| C | 0.30044 |
| T | 0.25780 |
| G | 0.16644 |

### 2 Results

We present additional results for empirical data in FASTA format on page 9 and for simulated paired-end NGS read data in FASTQ format on page 9.

#### 2.1 Empirical Data (Consensus FASTA)

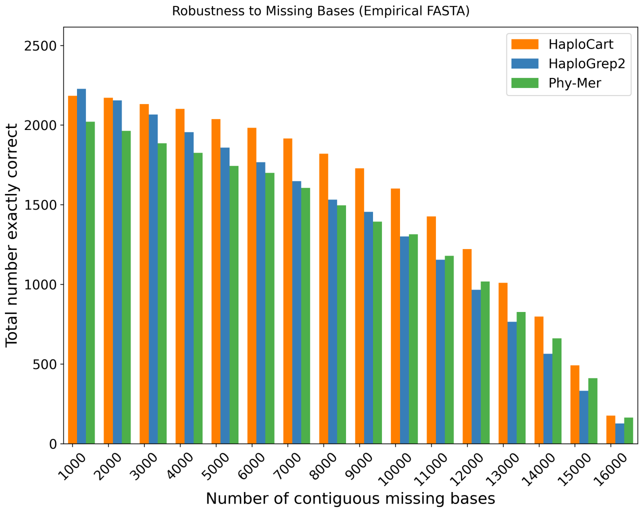

Supplementary Figure 1: **Total Number of Exactly Correct Predictions as a Function of the Number of Contiguous Masked Bases on Consensus FASTA Input.** Counts are provided for `HaploCart`, `Phy-Mer`, and `HaploGrep2`. `HaploCart` outperforms the other two programs from 2Kb up to 16Kb.

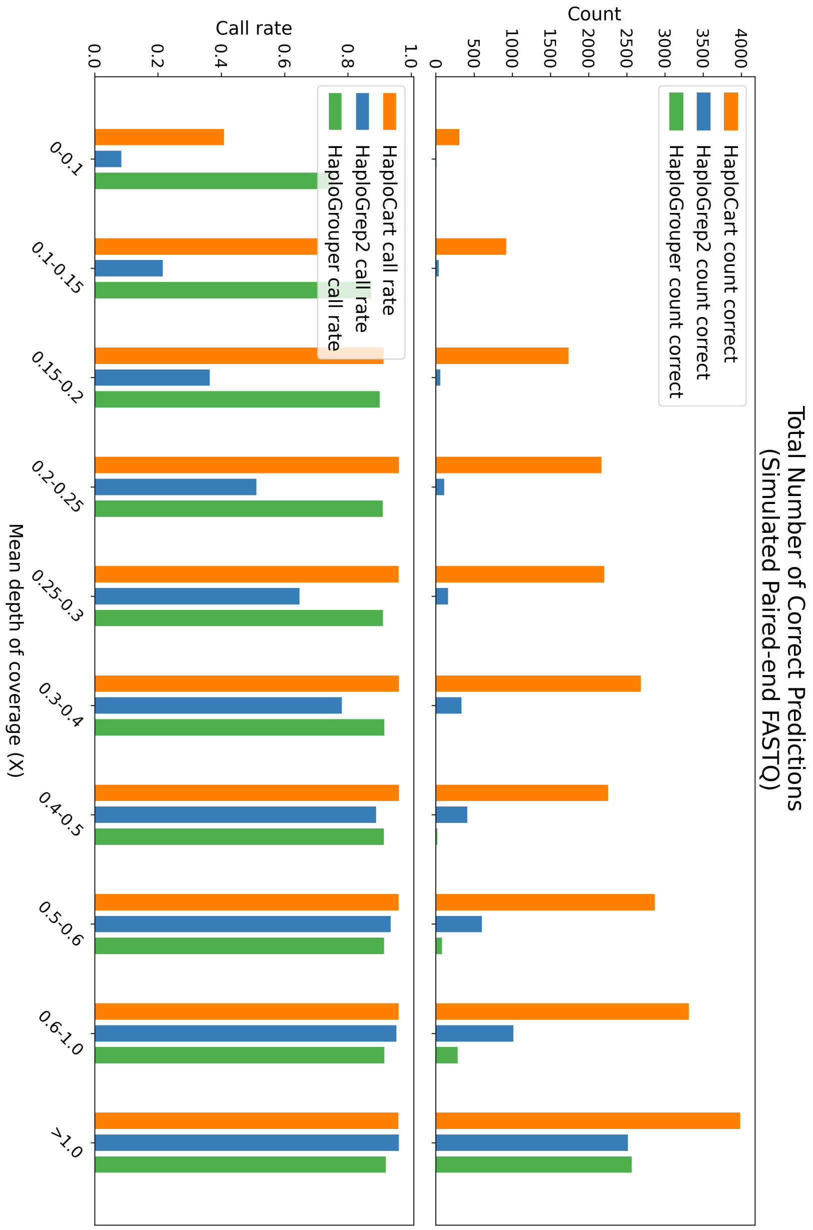

Supplementary Figure 2: **Total number of predictions and call rates on simulated paired-end FASTQ data.** $[TOP]$ Total number of predictions on the simulated replicates which exactly match the underlying haplogroup, as determined by running `HaploCheck` at full coverage. $[BOTTOM]$ Call rates (i.e. proportion of samples for which a haplogroup assignment is provided by the program). For each window, `HaploCart` outperforms `HaploGrep2` and `HaploGrouper` by providing more reliable haplogroup assignments at a higher call rate.

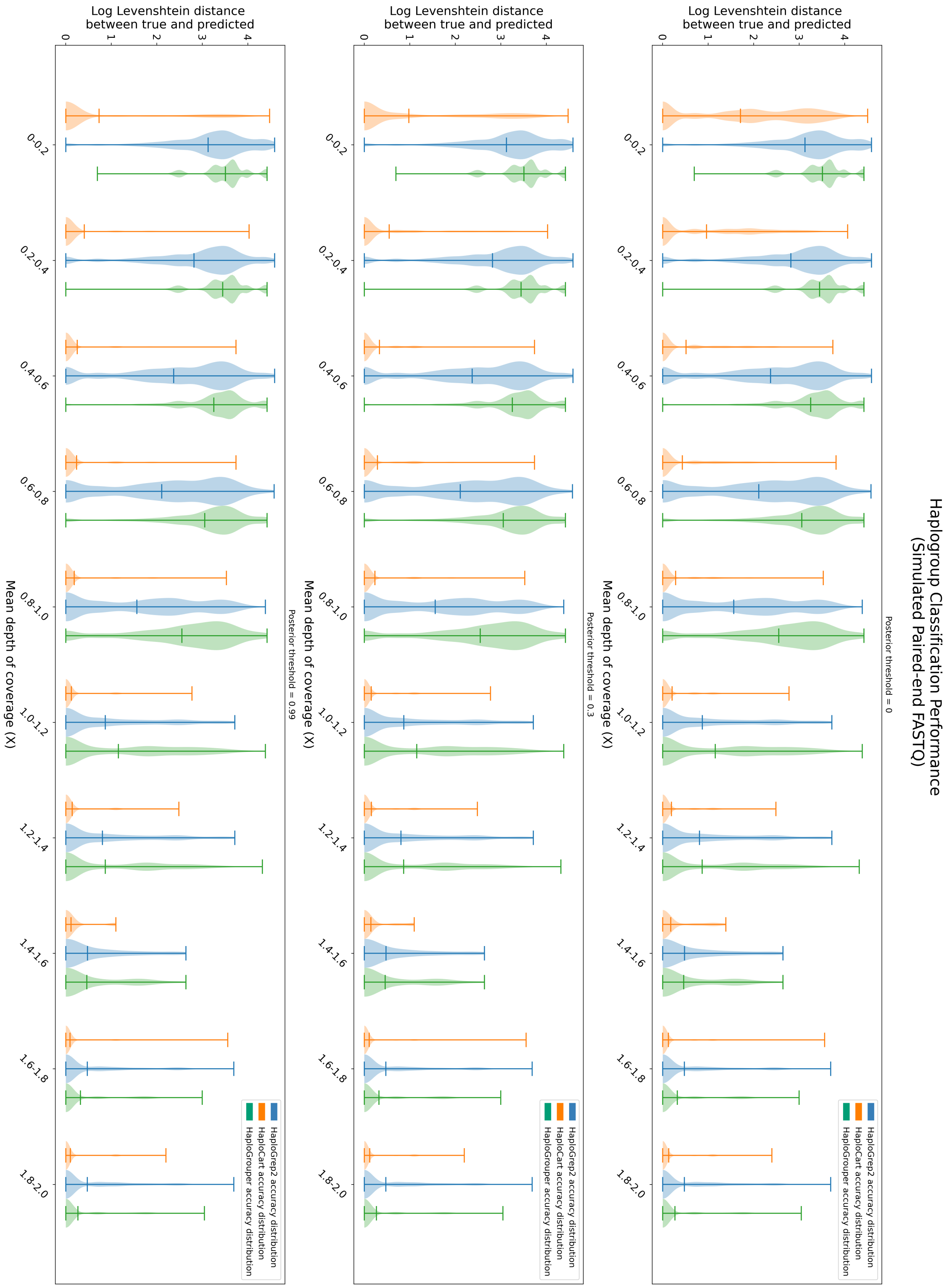

Supplementary Figure 3: **Simulated Paired-end Downsampled FASTQ Samples at Varying Posterior Thresholds.** Distribution of log edit (Levenshtein) distances on the simulated paired-end FASTQ dataset at three different lower thresholds (0, 0.3, 0.99) on the `HaploCart` posterior probability of the haplogroup assignment. No threshold is applied to `HaploGrep2` or `HaploGrouper`. We observe a clear improvement in the edit distances of the most anomalous predictions as the threshold increases, which demonstrates the utility of `HaploCart` posterior probabilities for use in quality control.

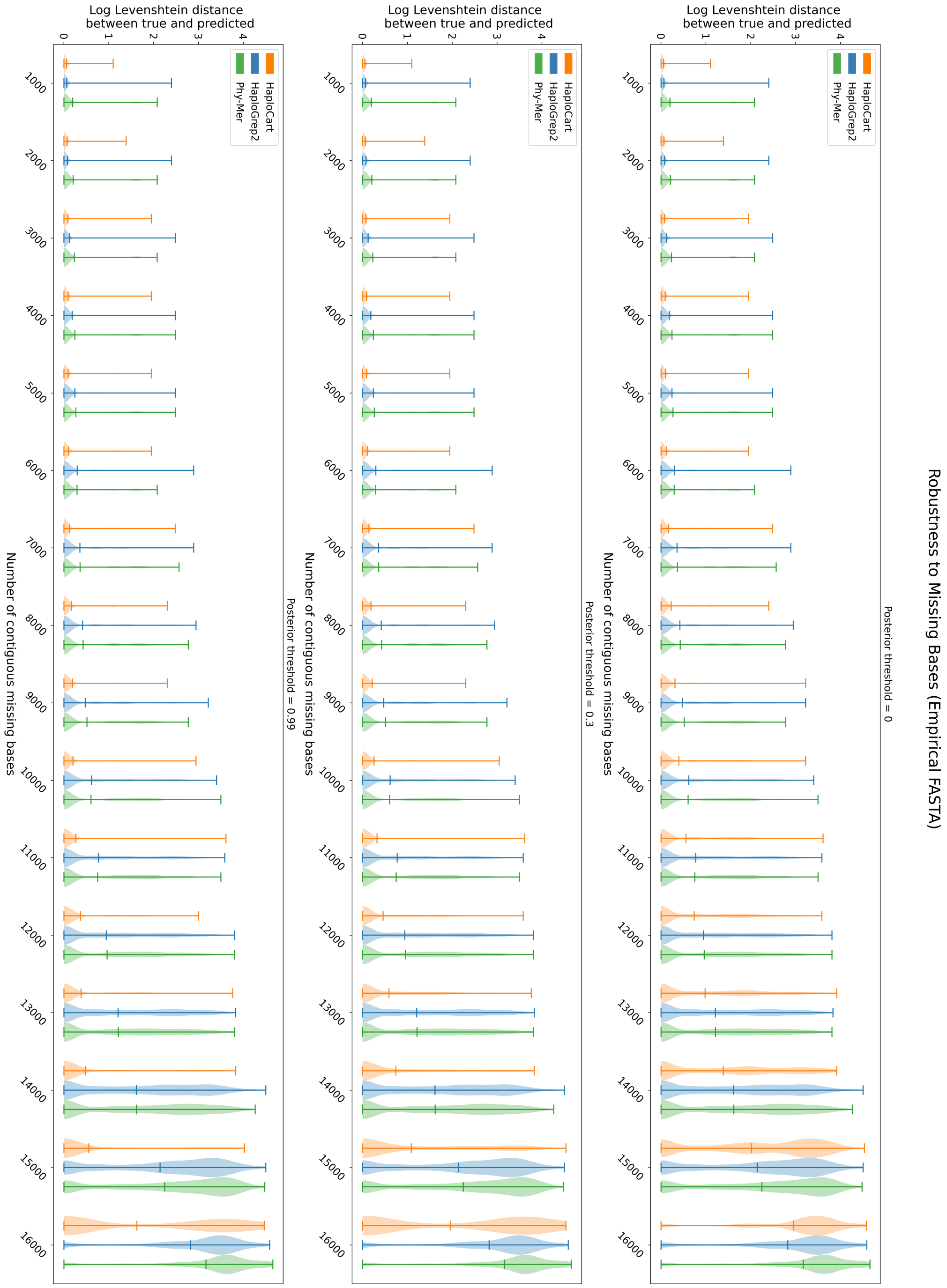

Supplementary Figure 4: **Masking Experiment on Consensus FASTA Input at Varying Posterior Thresholds.** Distribution of edit (Levenshtein) distances on the masked consensus FASTA dataset at three different lower thresholds (0, 0.3, 0.99) on the `HaploCart` posterior probability of the haplogroup assignment. No threshold is applied to `HaploGrep2` or `Phy-Mer`. We observe a clear improvement in the edit distance distribution for `HaploCart` as the threshold increases, demonstrating the utility of `HaploCart` posterior probabilities for use in quality control.

##### 2.1.1 HaploGrouper Predictions on Empirical Consensus FASTA Sequences

The vast majority of predictions concord with both HaploCart and HaploGrep2. For some samples, such as AY950293 and AY950293, HaploGrouper (prediction: M31a1) does not concord with a prediction shared by the other two programs (prediction: M31a1a). In another case (sample AF382002) we observe all three programs providing different predictions (H1a, H100, and HV_A73G), indicating that the underlying sample must be far from any sequence in the tree. We also see a case (sample AF346978) where HaploGrouper (prediction: HV0_T195C) agrees with HaploGrep2 (prediction: HV0+195) but not HaploCart (prediction: HV0d). Finally we see one case (sample AF381997) where HaploGrouper agrees with HaploCart on haplogroup R, but not with HaploGrep2 which calls haplogroup HV+73.

Since these data are empirical and may well constitute haplogroups outside the known tree, we do not have ground truth labels. Nonetheless, the fact that HaploGrouper does not seem to preferentially agree with either tool suggests that our program performs at least as well as HaploGrep2 at calling haplogroups on consensus sequences.

#### 2.2 Simulated Data in FASTQ Format

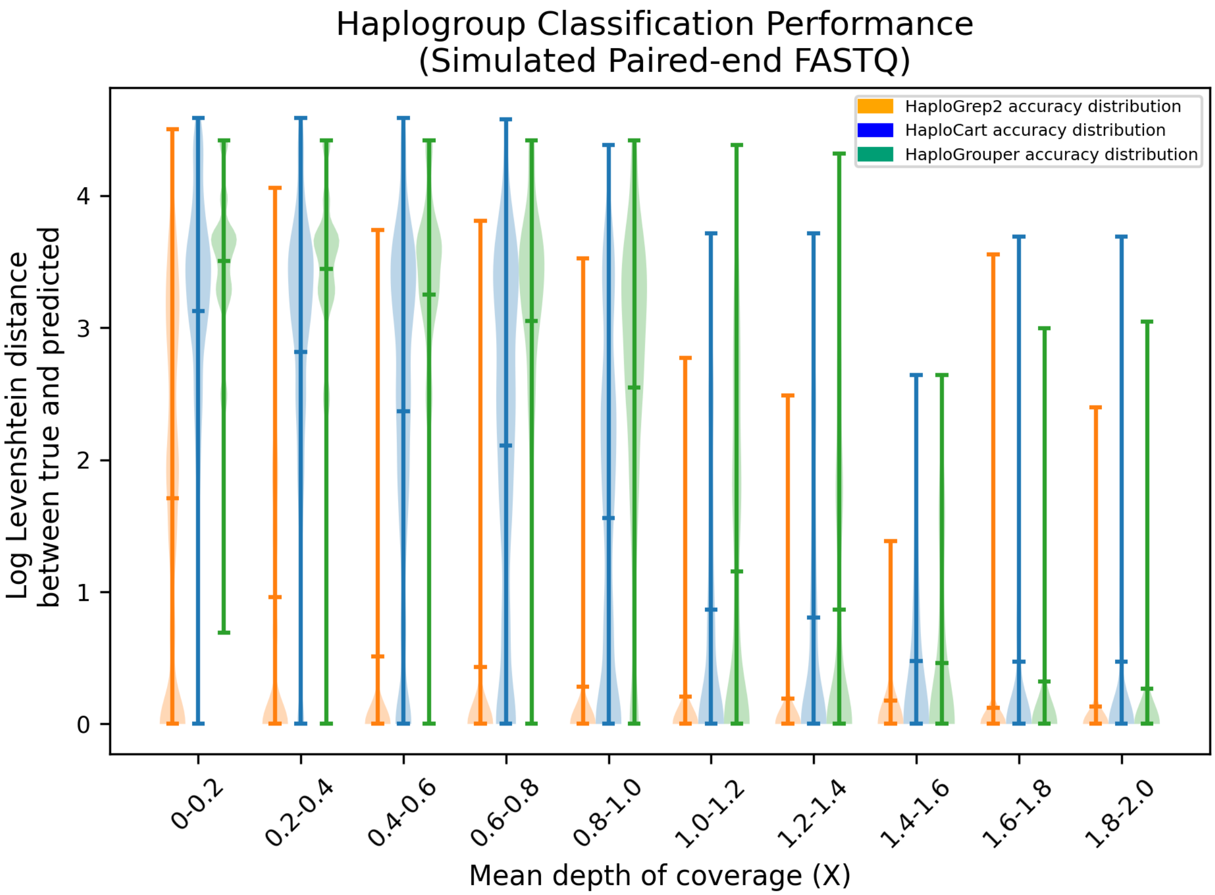

Supplementary Figure 5: **Distribution of Log Edit Distances between Ground Truth and Predicted Haplogroups on Simulated FASTQ Data** Distribution of edit (Levenshtein) distances between assigned and underlying haplogroup of replicates from the simulated dataset. Central line represent the arithmetic mean of the distribution. For each window, `HaploCart` outperforms `HaploGrep2` and `HaploGrouper` at all coverage windows as evidenced by the mean of the distributions. Note that unlike `HaploGrep2` and `HaploGrouper`, `HaploCart` makes a prediction if even a single read maps to the graph.

#### 2.3 Simulated Paired-end FASTQ Posterior Plots

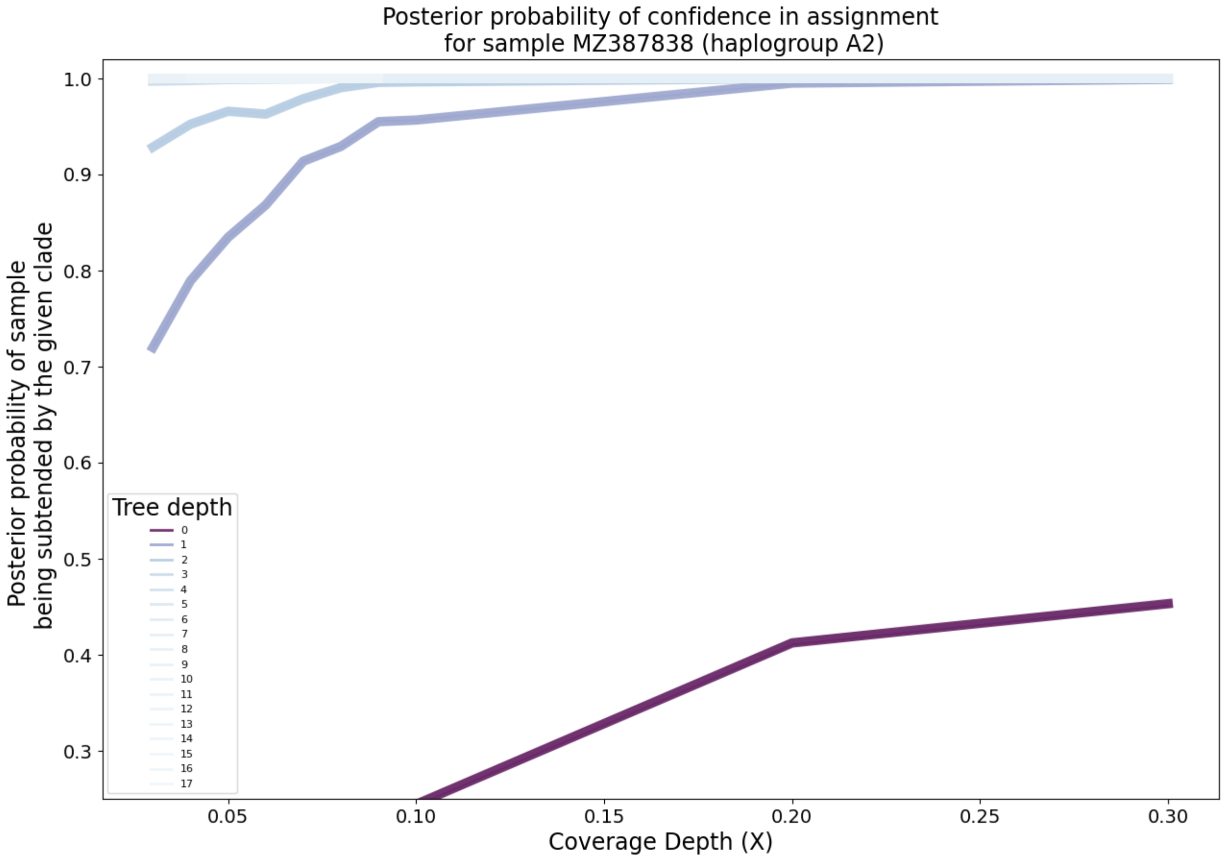

Supplementary Figure 6: **Clade-level posterior probabilities of haplogroup assignment on simulated paired-end FASTQ data.** Each lineplot represents the mean over replicates at a fixed depth on the mitochondrial tree. The darker the line, the more basal the haplogroups.

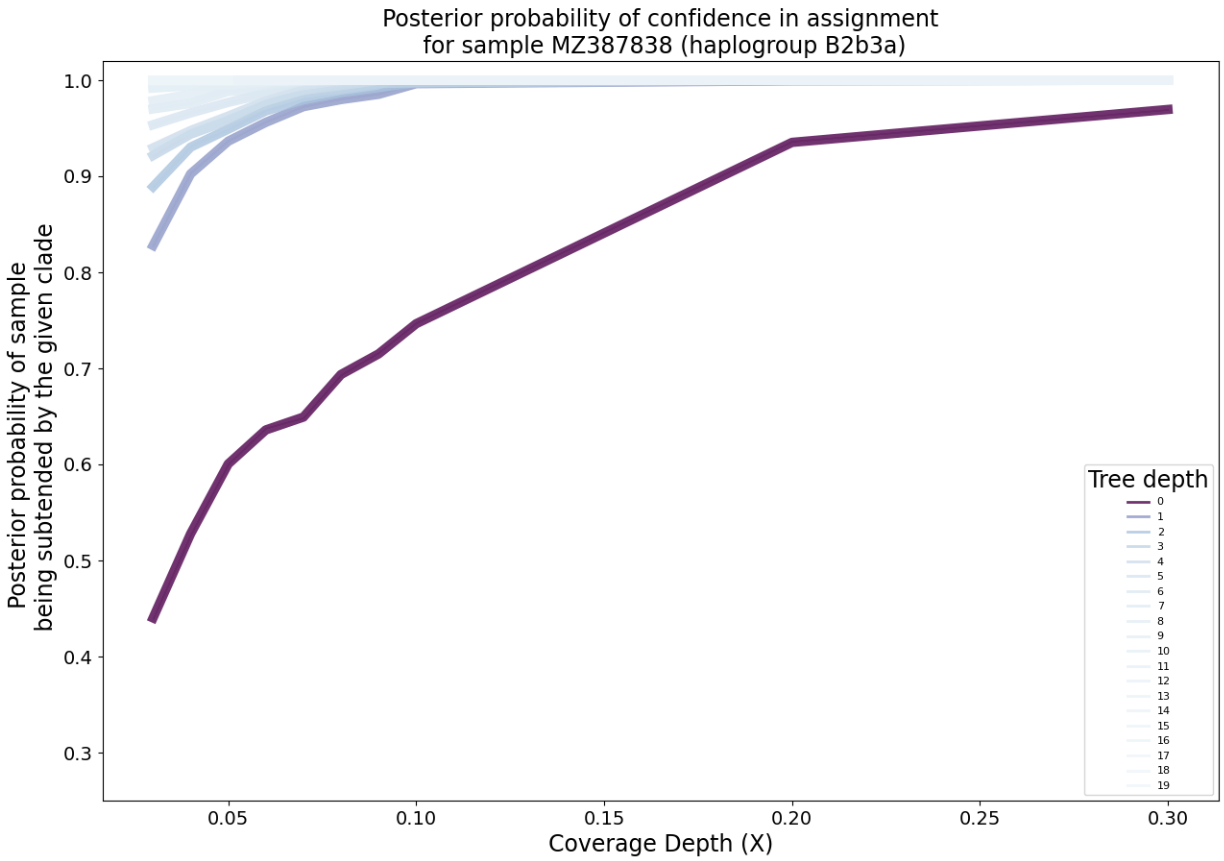

Supplementary Figure 7: **Clade-level posterior probabilities of haplogroup assignment on simulated paired-end FASTQ data.** Each lineplot represents the mean over replicates at a fixed depth on the mitochondrial tree. The darker the line, the more basal the haplogroups.

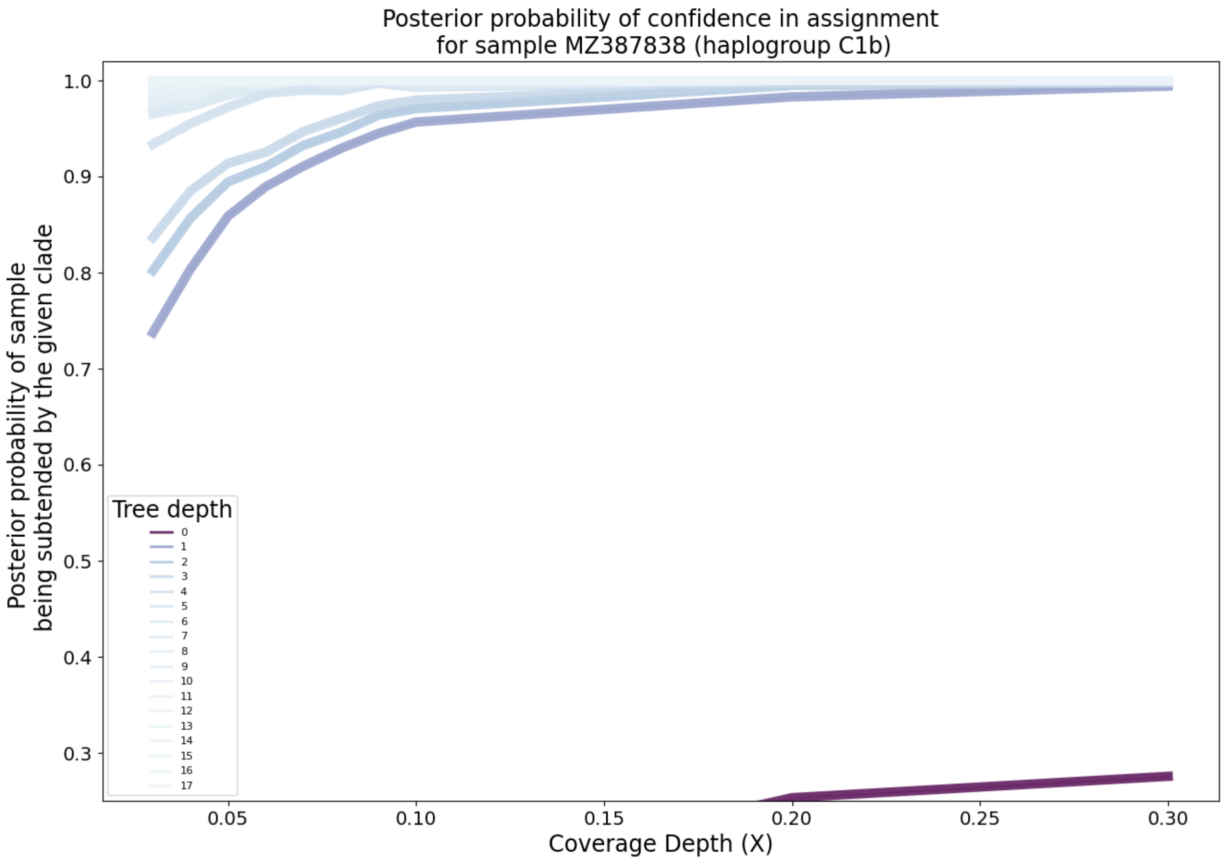

Supplementary Figure 8: **Clade-level posterior probabilities of haplogroup assignment on simulated paired-end FASTQ data.** Each lineplot represents the mean over replicates at a fixed depth on the mitochondrial tree. The darker the line, the more basal the haplogroups.

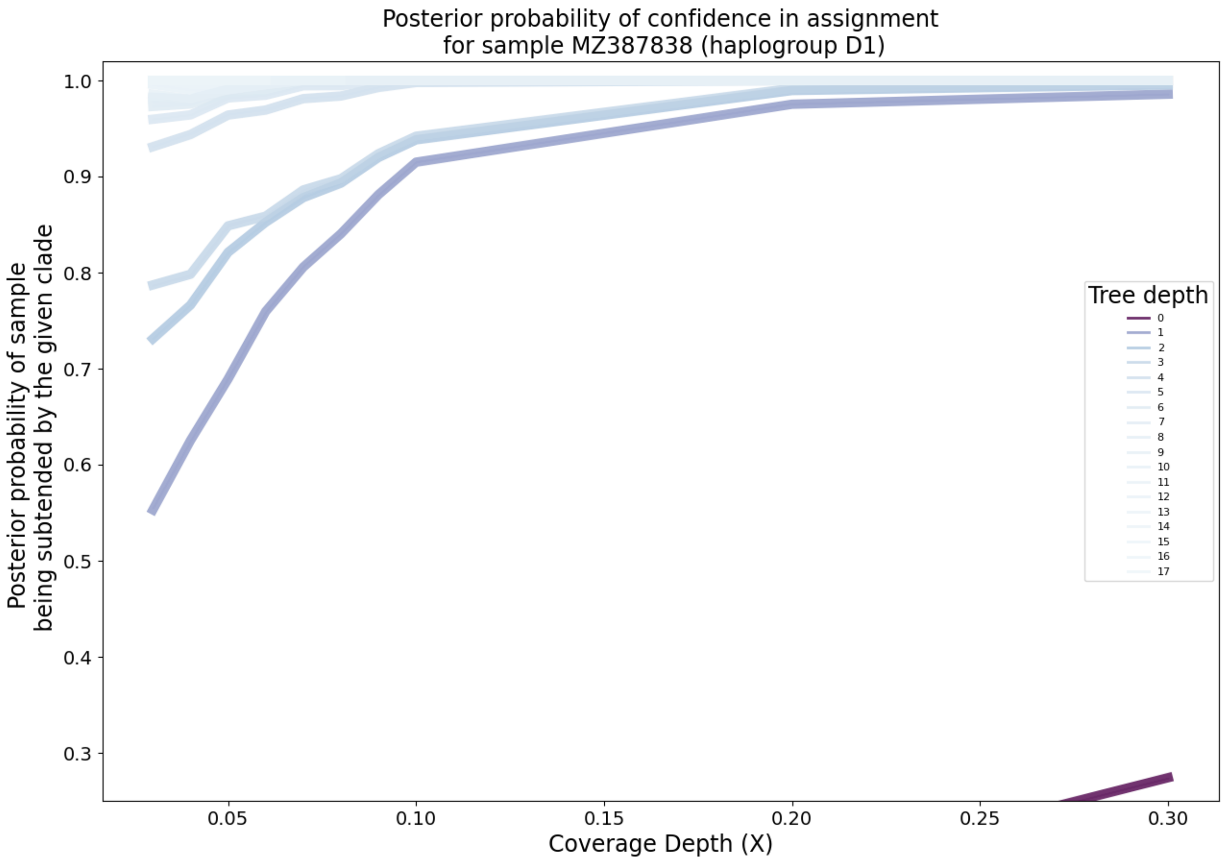

Supplementary Figure 9: **Clade-level posterior probabilities of haplogroup assignment on simulated paired-end FASTQ data.** Each lineplot represents the mean over replicates at a fixed depth on the mitochondrial tree. The darker the line, the more basal the haplogroups.

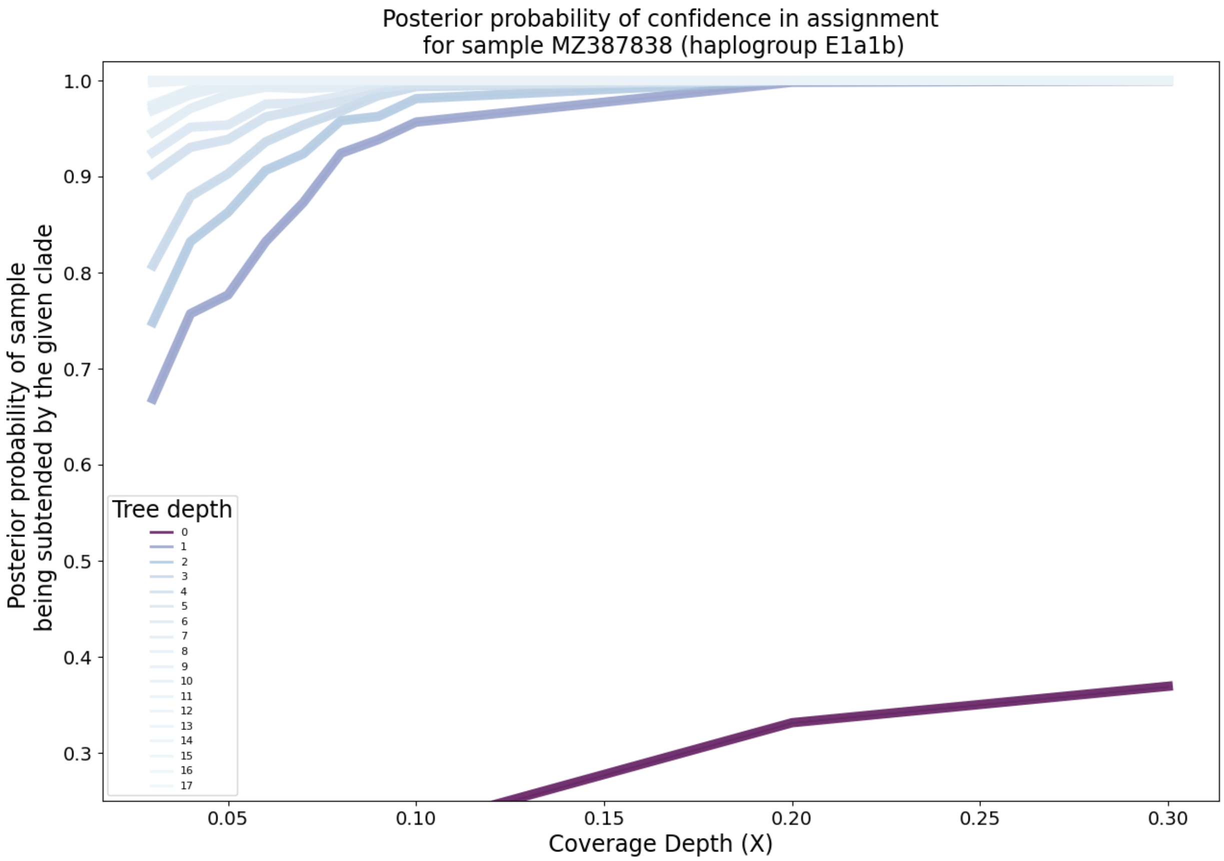

Supplementary Figure 10: **Clade-level posterior probabilities of haplogroup assignment on simulated paired-end FASTQ data.** Each lineplot represents the mean over replicates at a fixed depth on the mitochondrial tree. The darker the line, the more basal the haplogroups.

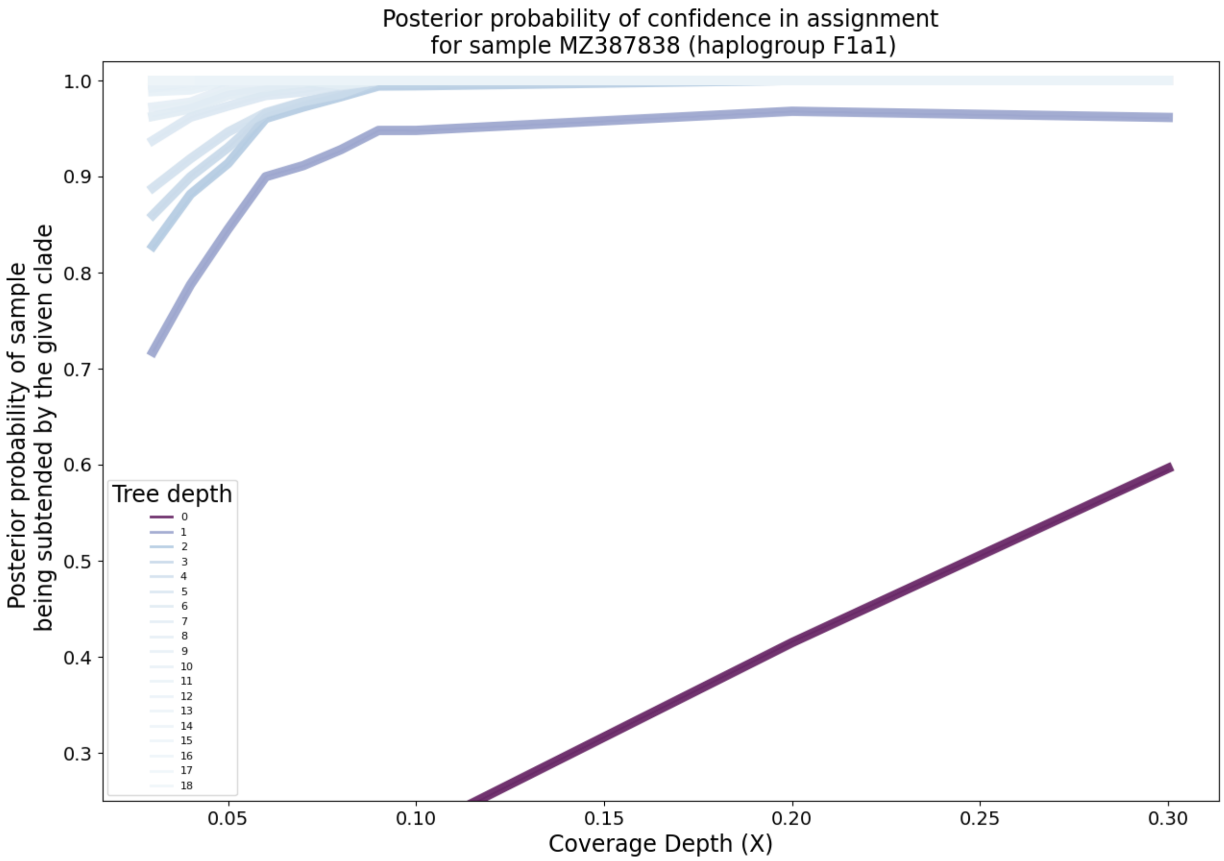

Supplementary Figure 11: **Clade-level posterior probabilities of haplogroup assignment on simulated paired-end FASTQ data.** Each lineplot represents the mean over replicates at a fixed depth on the mitochondrial tree. The darker the line, the more basal the haplogroups.

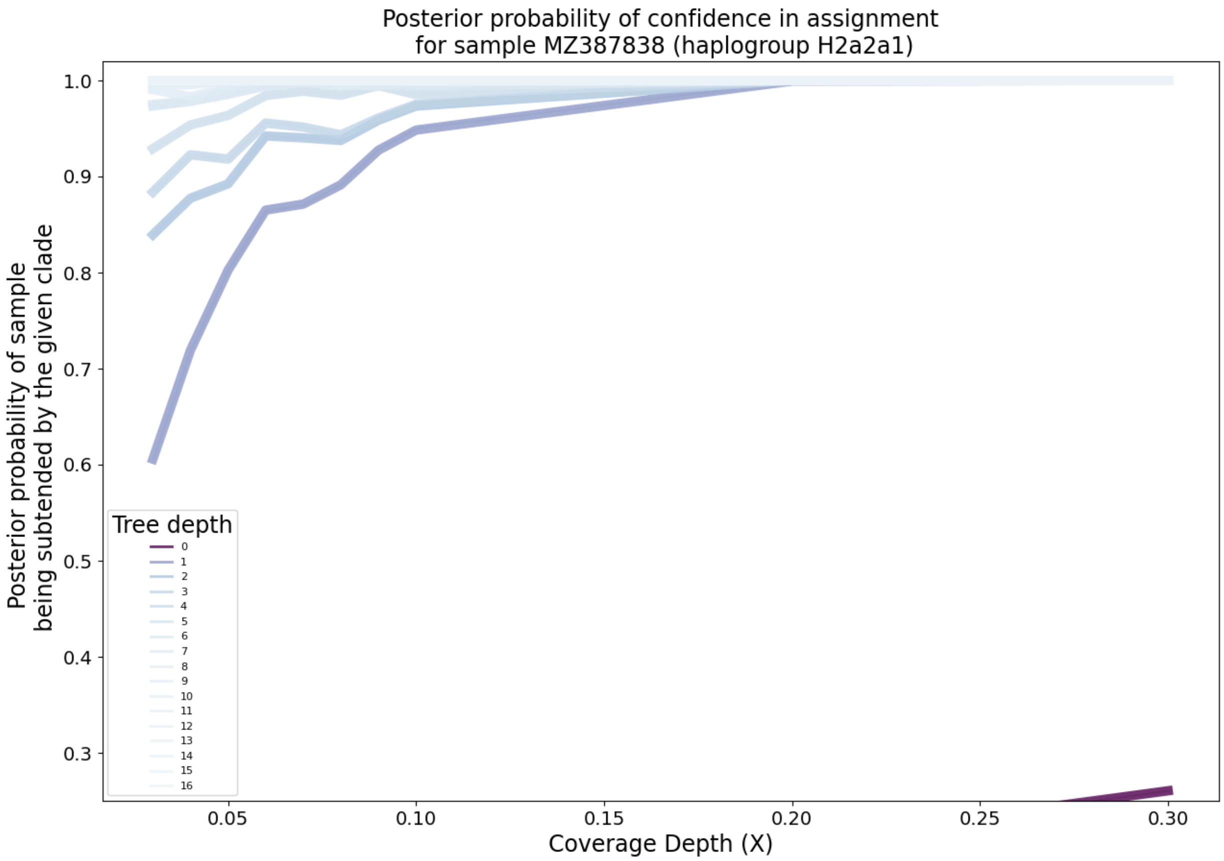

Supplementary Figure 12: **Clade-level posterior probabilities of haplogroup assignment on simulated paired-end FASTQ data.** Each lineplot represents the mean over replicates at a fixed depth on the mitochondrial tree. The darker the line, the more basal the haplogroups.

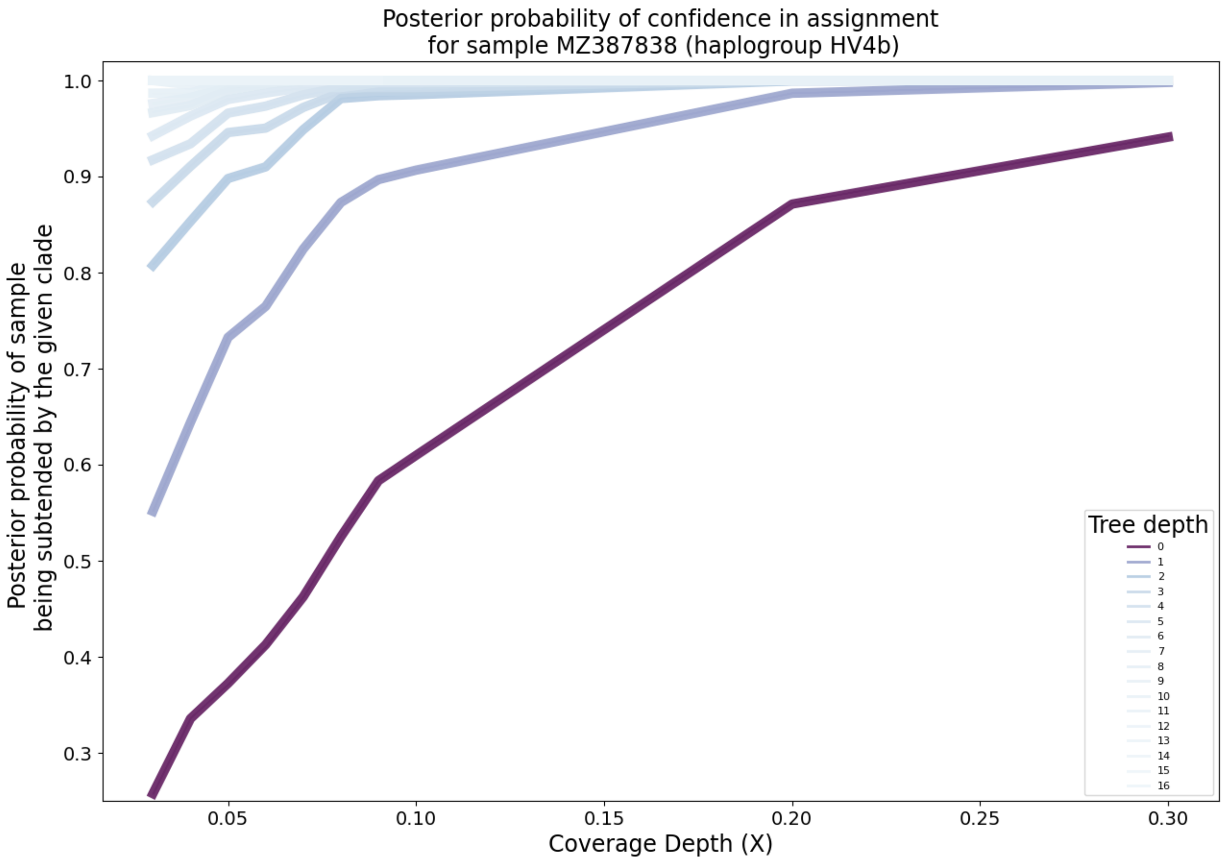

Supplementary Figure 13: **Clade-level posterior probabilities of haplogroup assignment on simulated paired-end FASTQ data.** Each lineplot represents the mean over replicates at a fixed depth on the mitochondrial tree. The darker the line, the more basal the haplogroups.

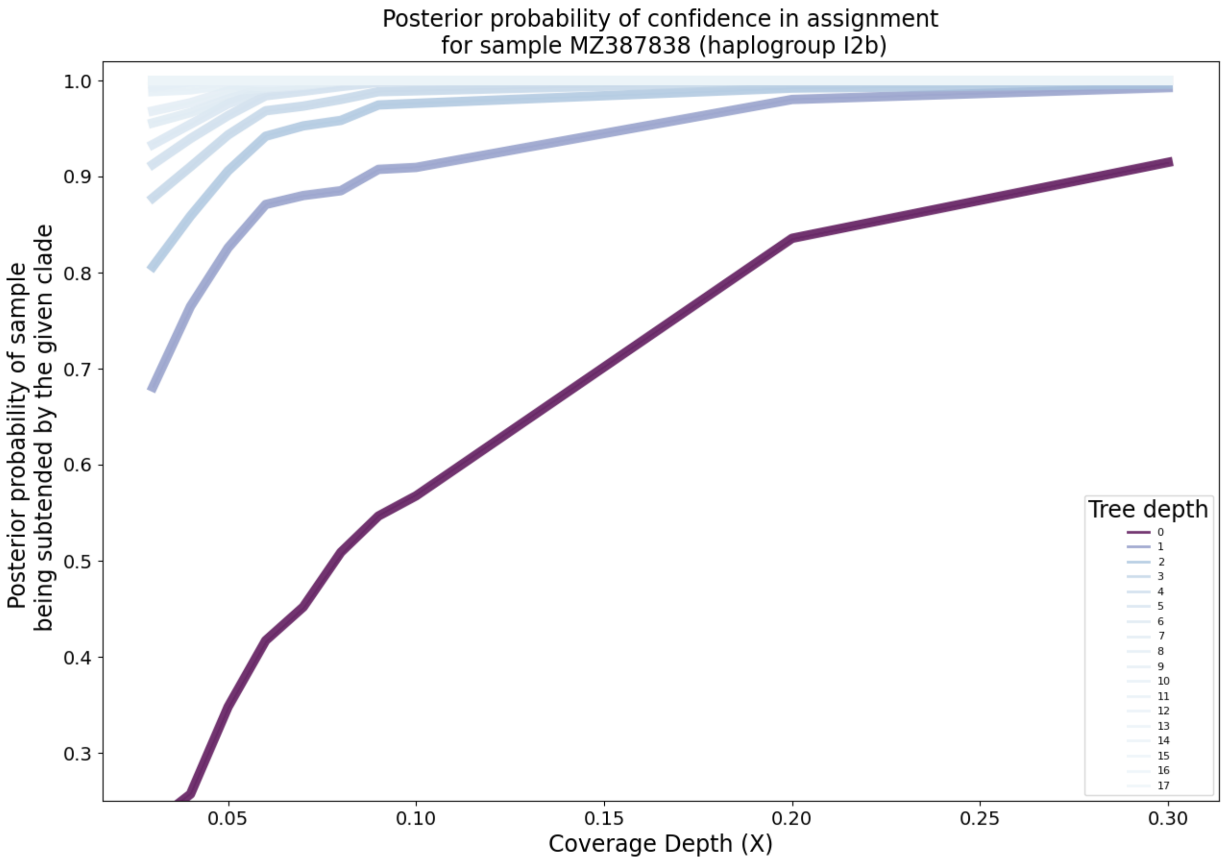

Supplementary Figure 14: **Clade-level posterior probabilities of haplogroup assignment on simulated paired-end FASTQ data.** Each lineplot represents the mean over replicates at a fixed depth on the mitochondrial tree. The darker the line, the more basal the haplogroups.

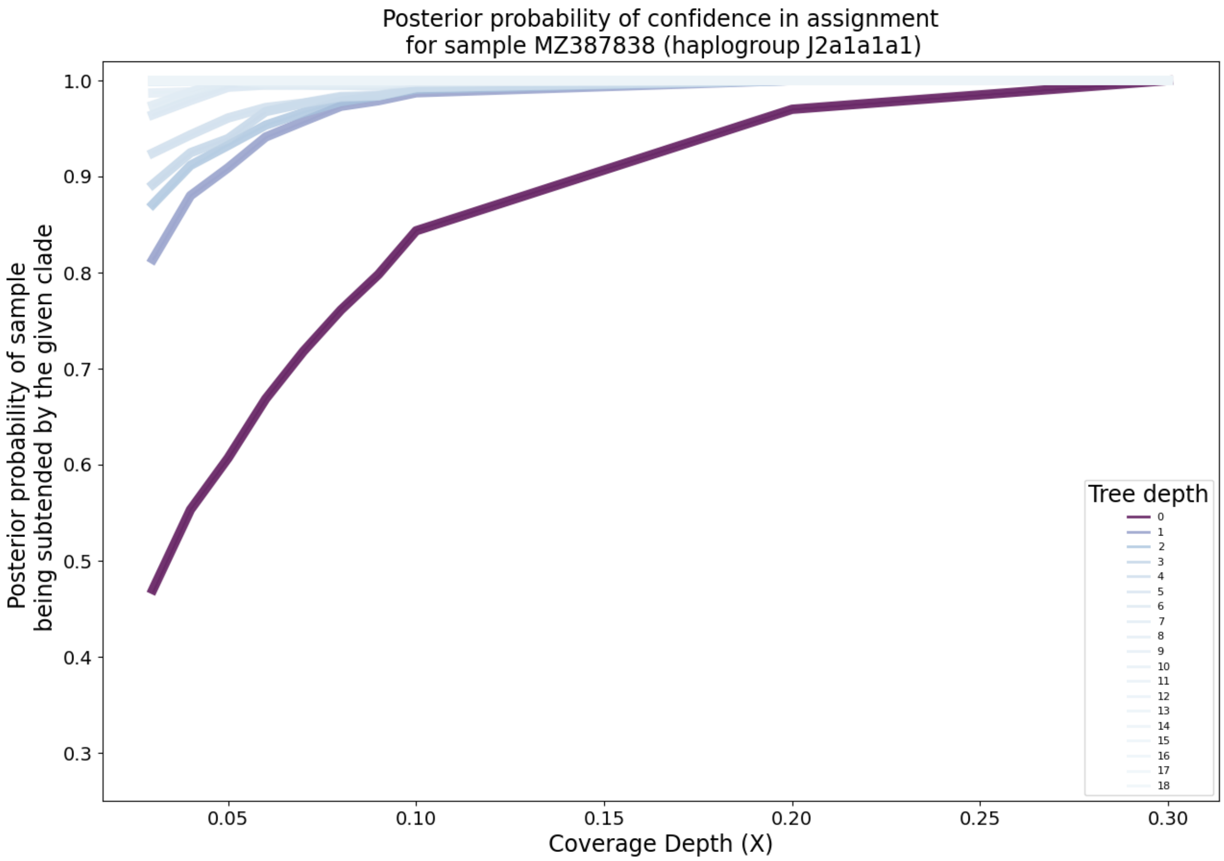

Supplementary Figure 15: **Clade-level posterior probabilities of haplogroup assignment on simulated paired-end FASTQ data.** Each lineplot represents the mean over replicates at a fixed depth on the mitochondrial tree. The darker the line, the more basal the haplogroups.

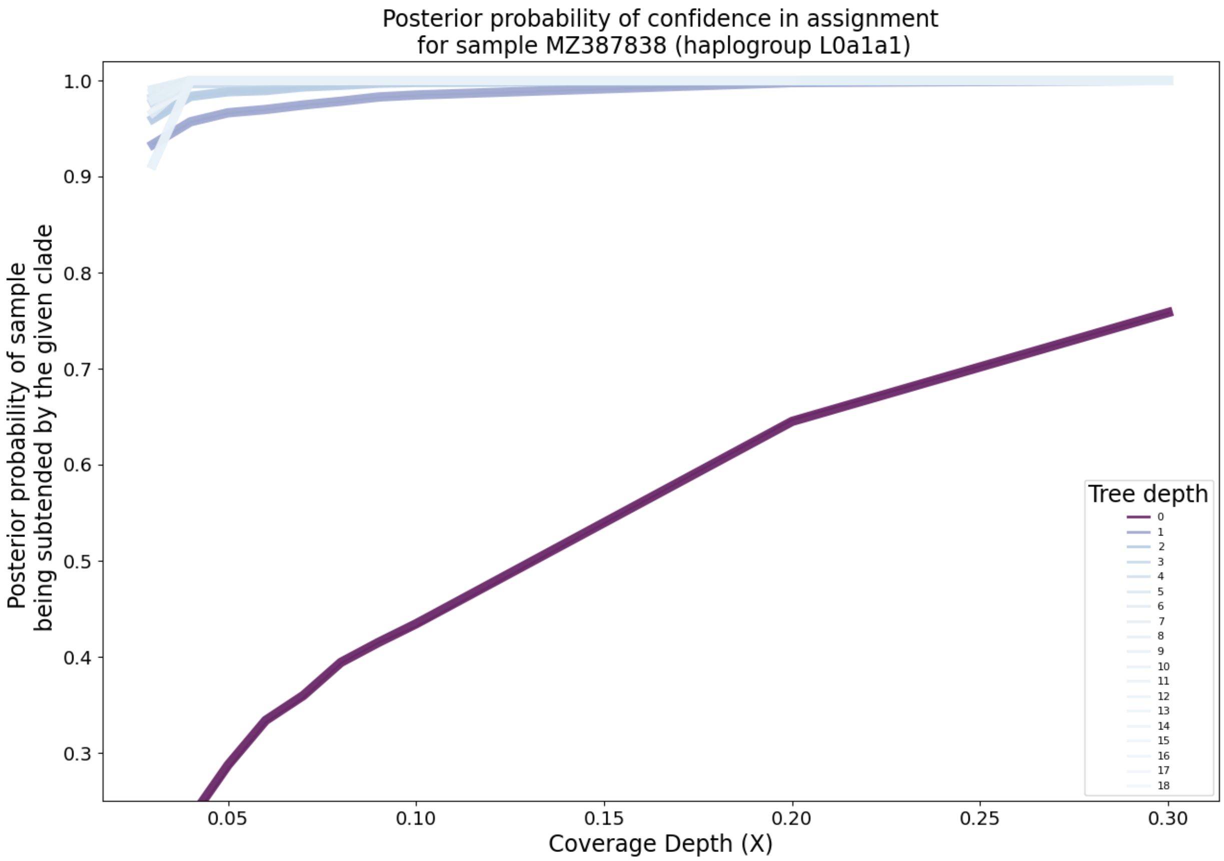

Supplementary Figure 16: **Clade-level posterior probabilities of haplogroup assignment on simulated paired-end FASTQ data.** Each lineplot represents the mean over replicates at a fixed depth on the mitochondrial tree. The darker the line, the more basal the haplogroups.

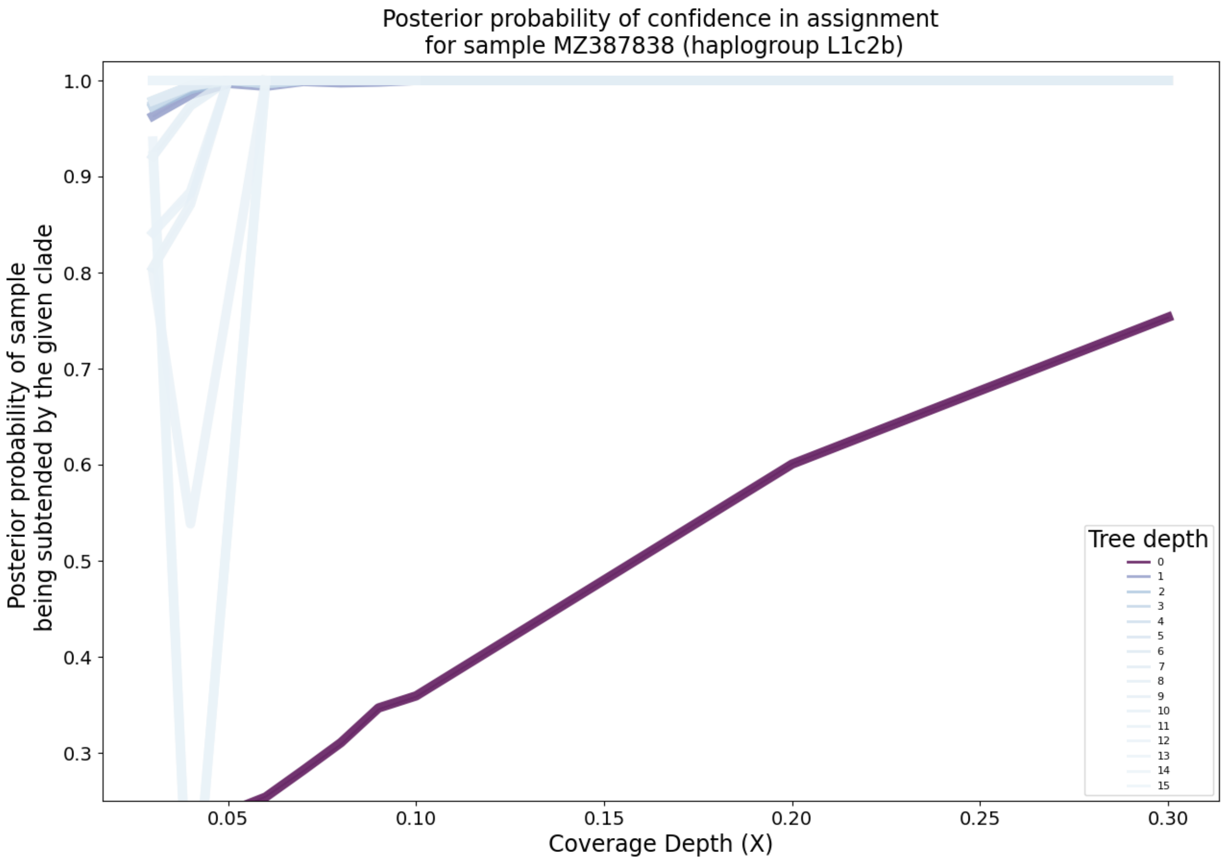

Supplementary Figure 17: **Clade-level posterior probabilities of haplogroup assignment on simulated paired-end FASTQ data.** Each lineplot represents the mean over replicates at a fixed depth on the mitochondrial tree. The darker the line, the more basal the haplogroups.

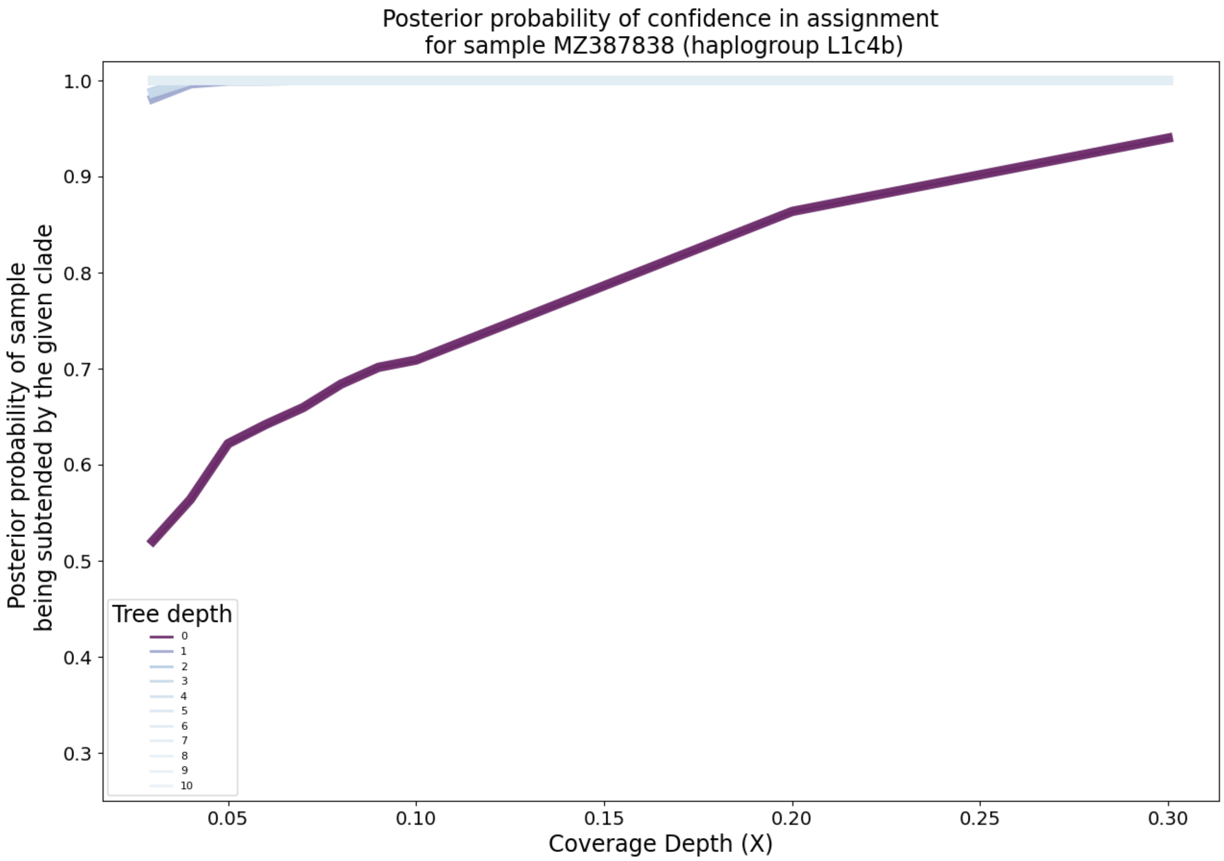

Supplementary Figure 18: **Clade-level posterior probabilities of haplogroup assignment on simulated paired-end FASTQ data.** Each lineplot represents the mean over replicates at a fixed depth on the mitochondrial tree. The darker the line, the more basal the haplogroups.

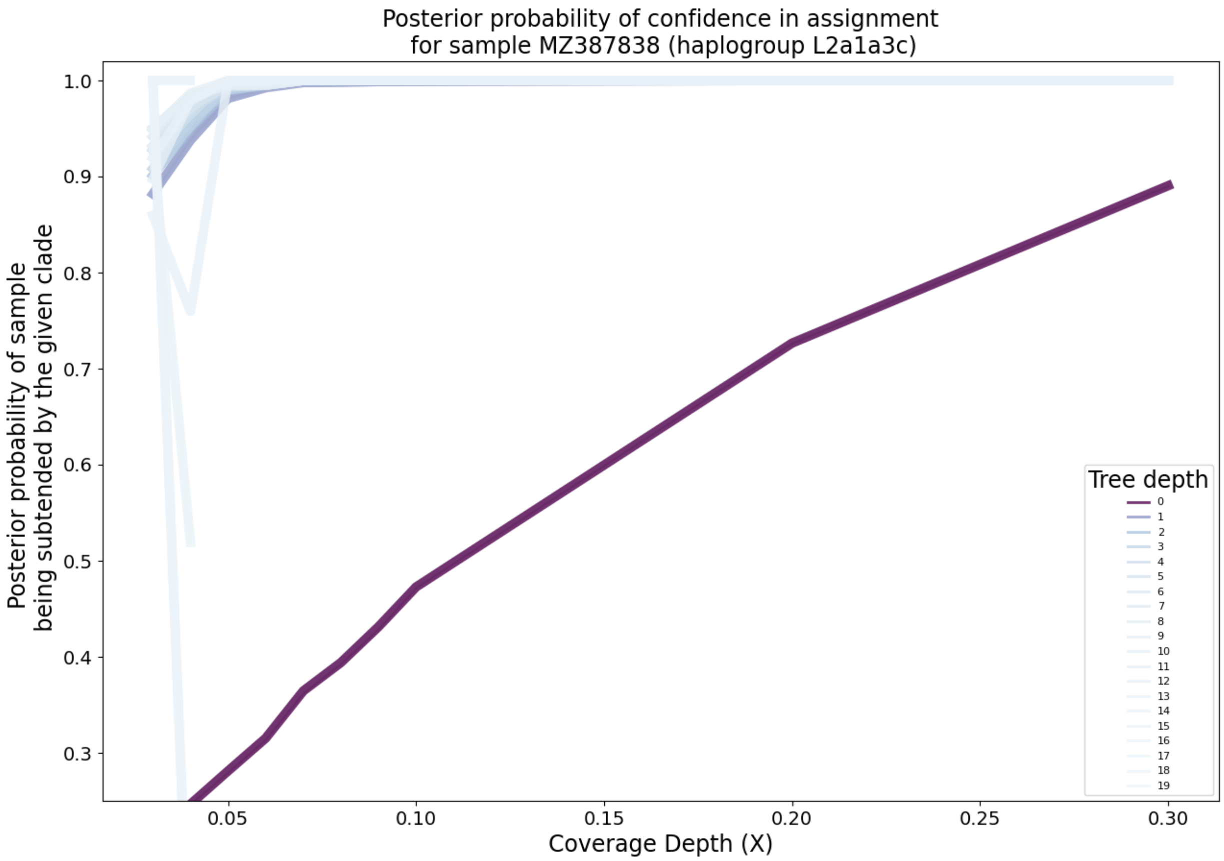

Supplementary Figure 19: **Clade-level posterior probabilities of haplogroup assignment on simulated paired-end FASTQ data.** Each lineplot represents the mean over replicates at a fixed depth on the mitochondrial tree. The darker the line, the more basal the haplogroups.

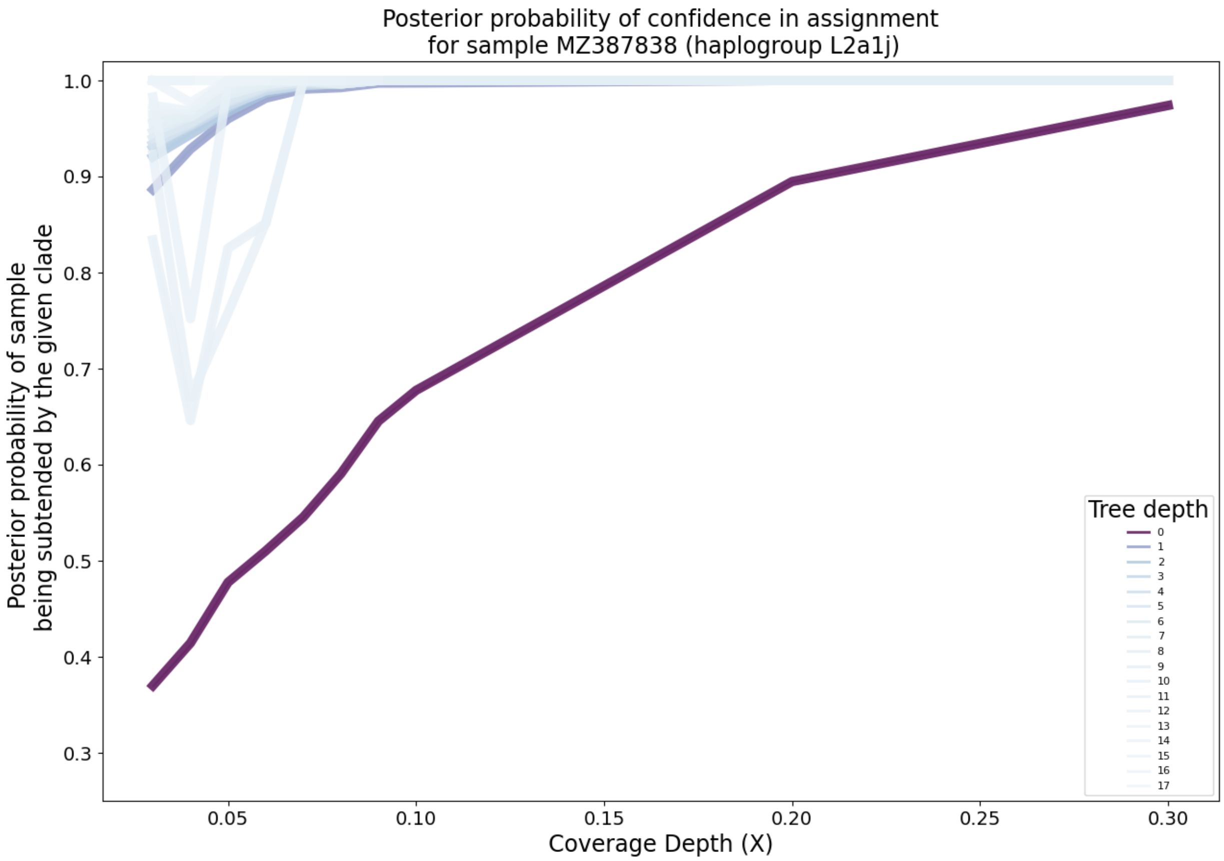

Supplementary Figure 20: **Clade-level posterior probabilities of haplogroup assignment on simulated paired-end FASTQ data.** Each lineplot represents the mean over replicates at a fixed depth on the mitochondrial tree. The darker the line, the more basal the haplogroups.

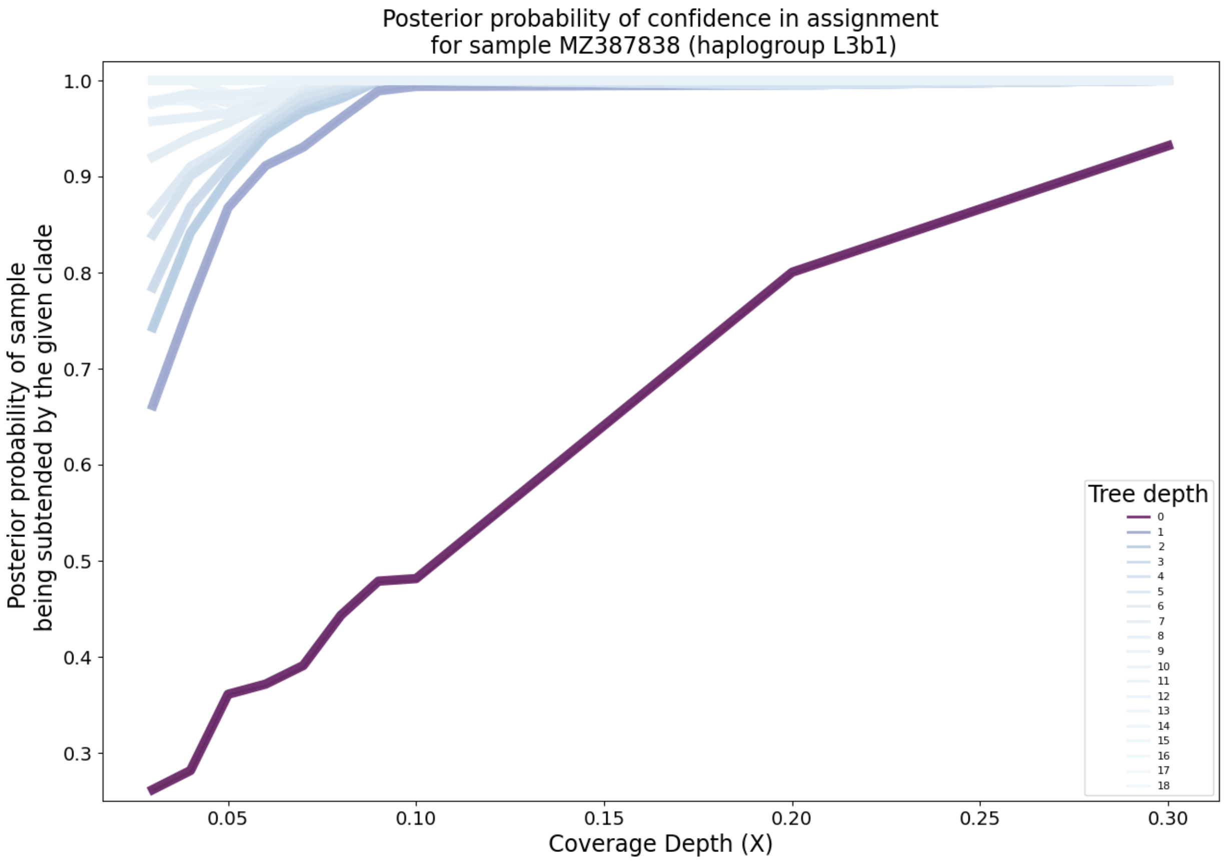

Supplementary Figure 21: **Clade-level posterior probabilities of haplogroup assignment on simulated paired-end FASTQ data.** Each lineplot represents the mean over replicates at a fixed depth on the mitochondrial tree. The darker the line, the more basal the haplogroups.

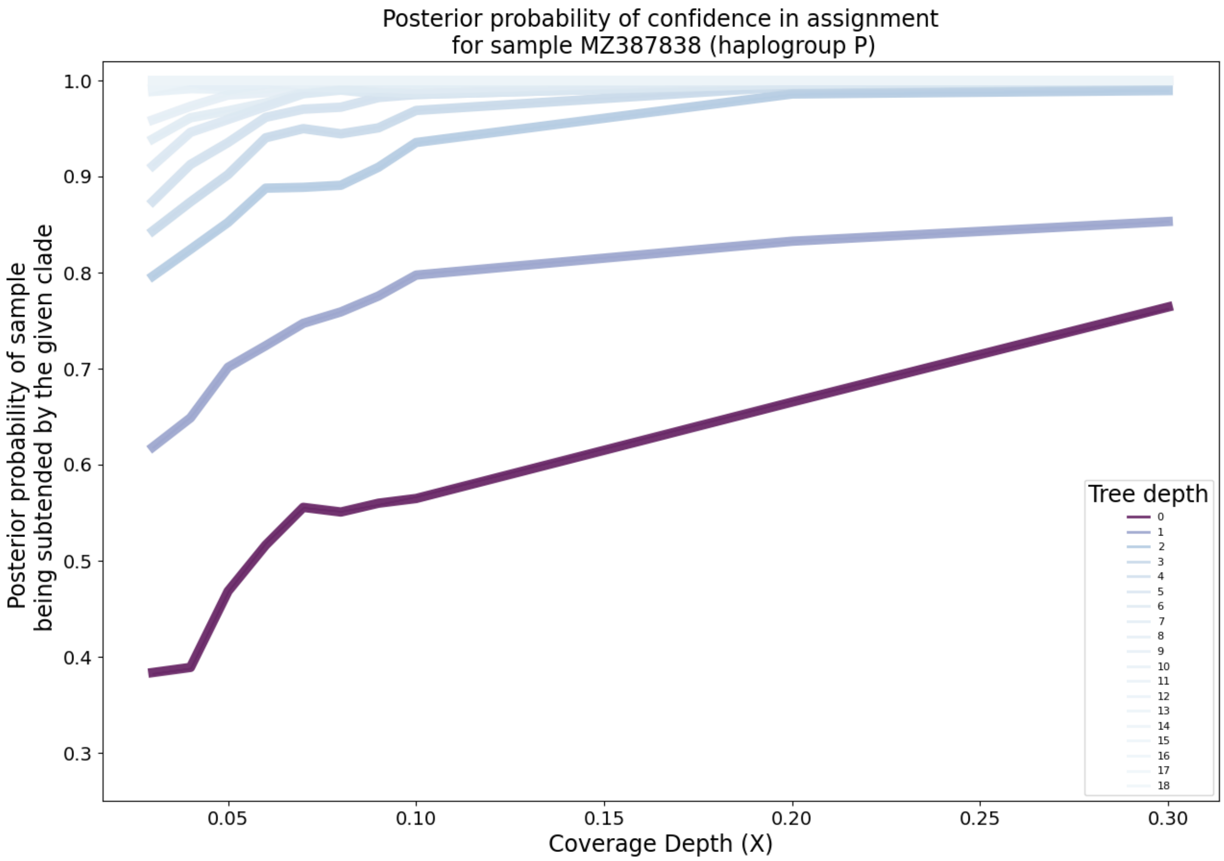

Supplementary Figure 22: **Clade-level posterior probabilities of haplogroup assignment on simulated paired-end FASTQ data.** Each lineplot represents the mean over replicates at a fixed depth on the mitochondrial tree. The darker the line, the more basal the haplogroups.

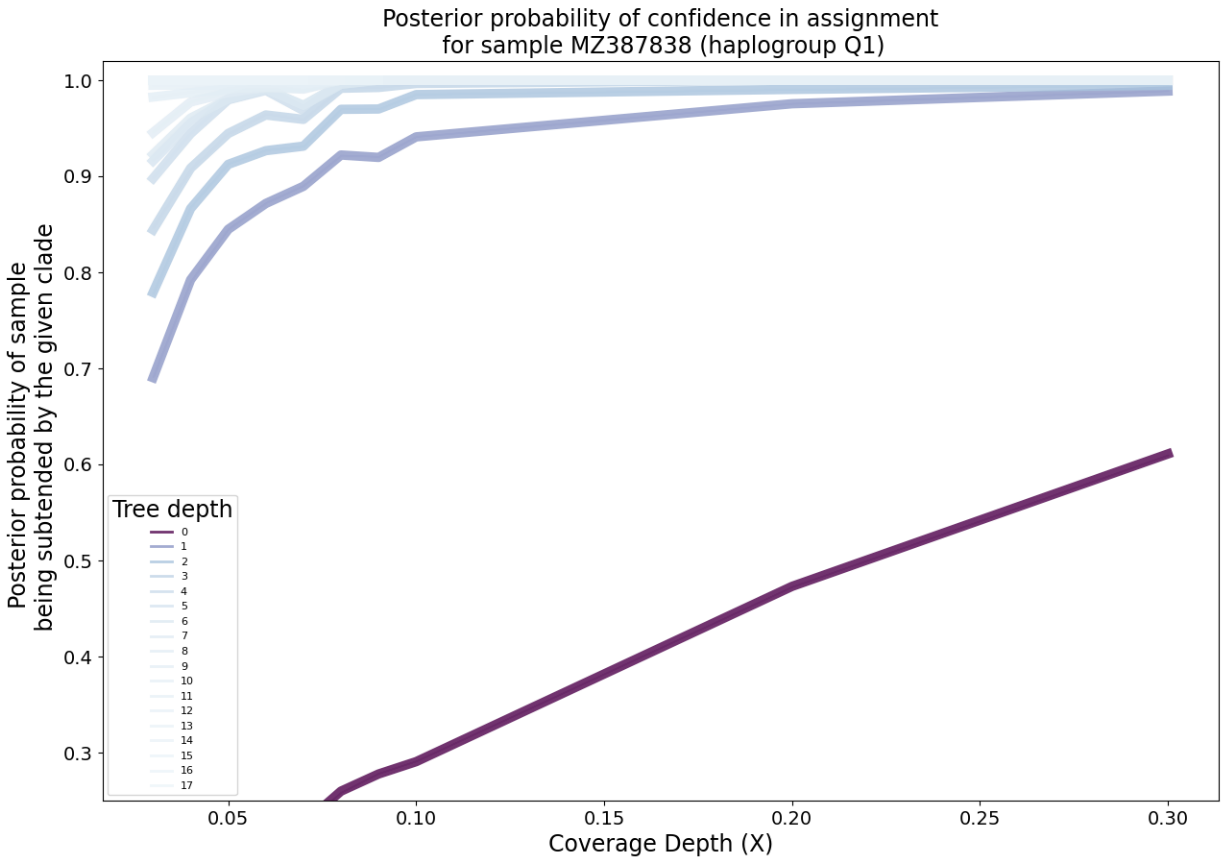

Supplementary Figure 23: **Clade-level posterior probabilities of haplogroup assignment on simulated paired-end FASTQ data.** Each lineplot represents the mean over replicates at a fixed depth on the mitochondrial tree. The darker the line, the more basal the haplogroups.

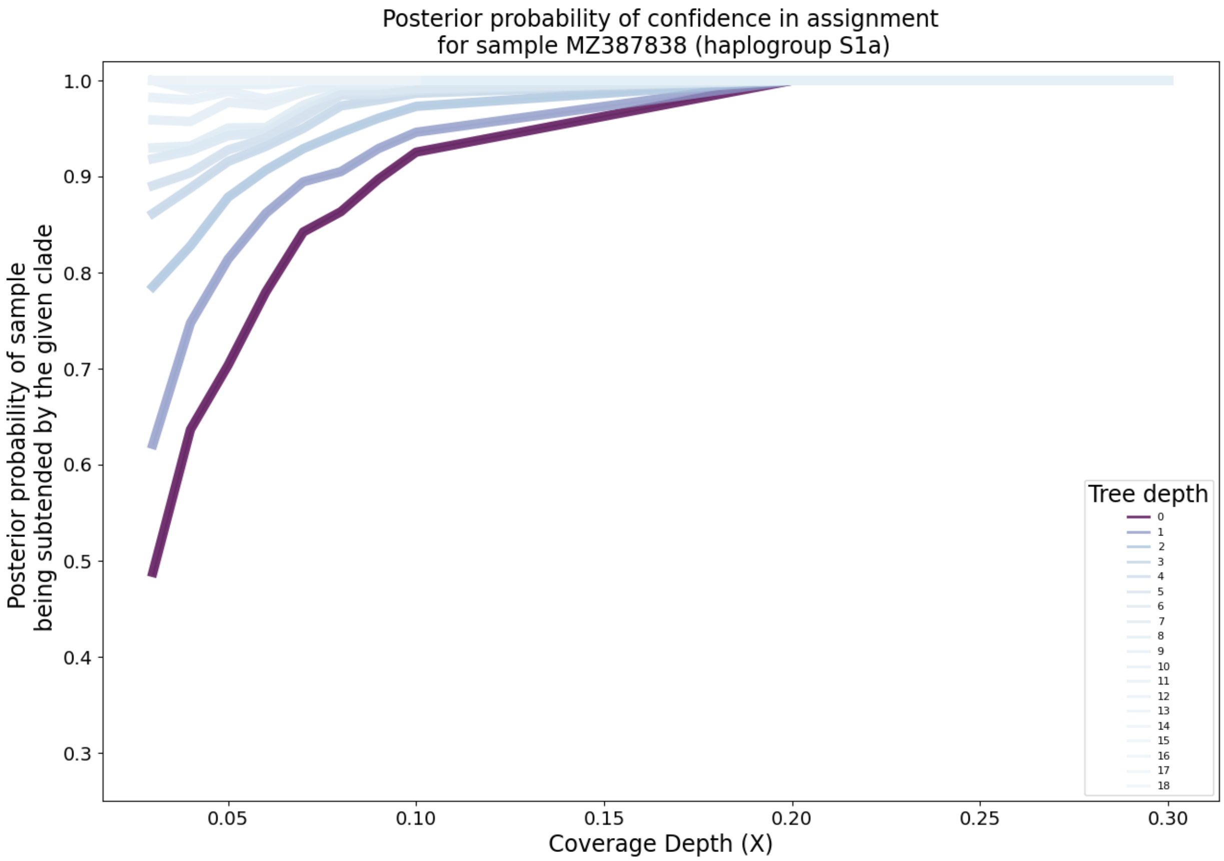

Supplementary Figure 24: **Clade-level posterior probabilities of haplogroup assignment on simulated paired-end FASTQ data.** Each lineplot represents the mean over replicates at a fixed depth on the mitochondrial tree. The darker the line, the more basal the haplogroups.

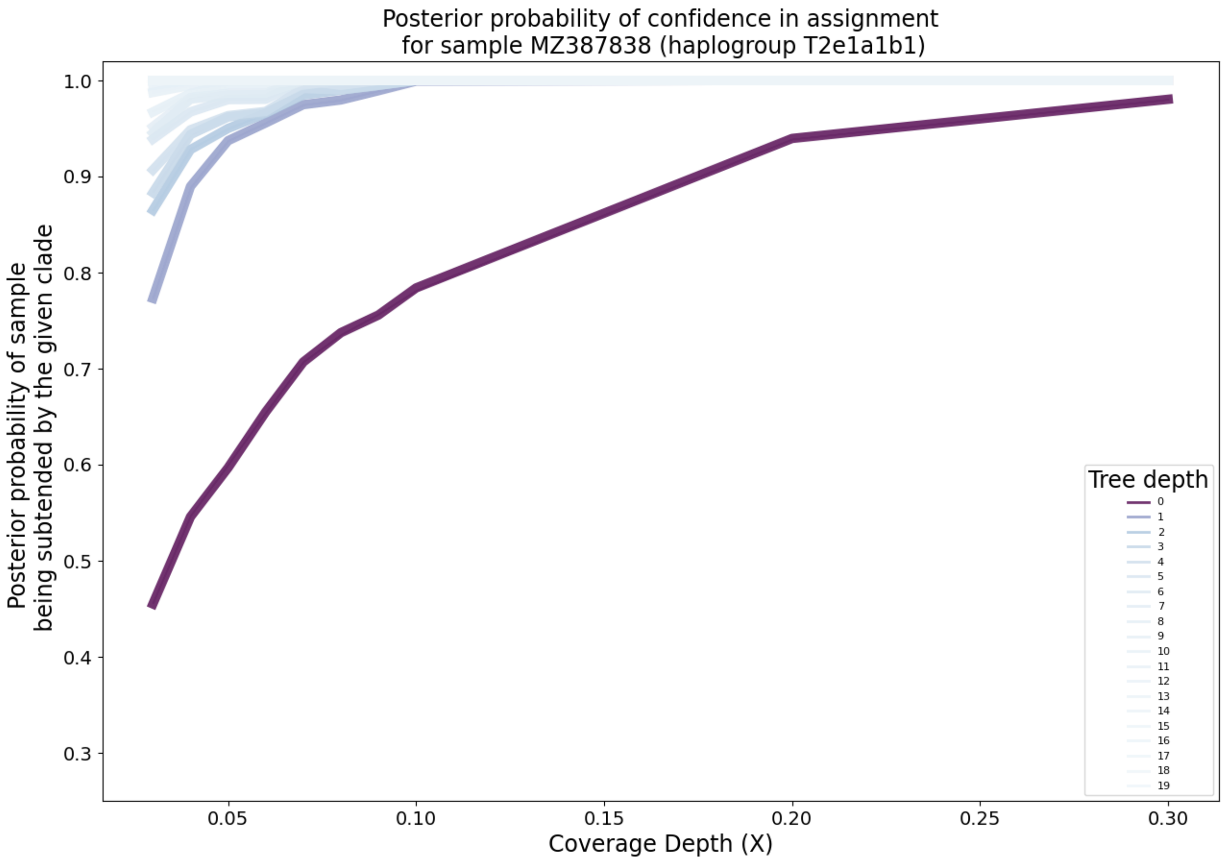

Supplementary Figure 25: **Clade-level posterior probabilities of haplogroup assignment on simulated paired-end FASTQ data.** Each lineplot represents the mean over replicates at a fixed depth on the mitochondrial tree. The darker the line, the more basal the haplogroups.

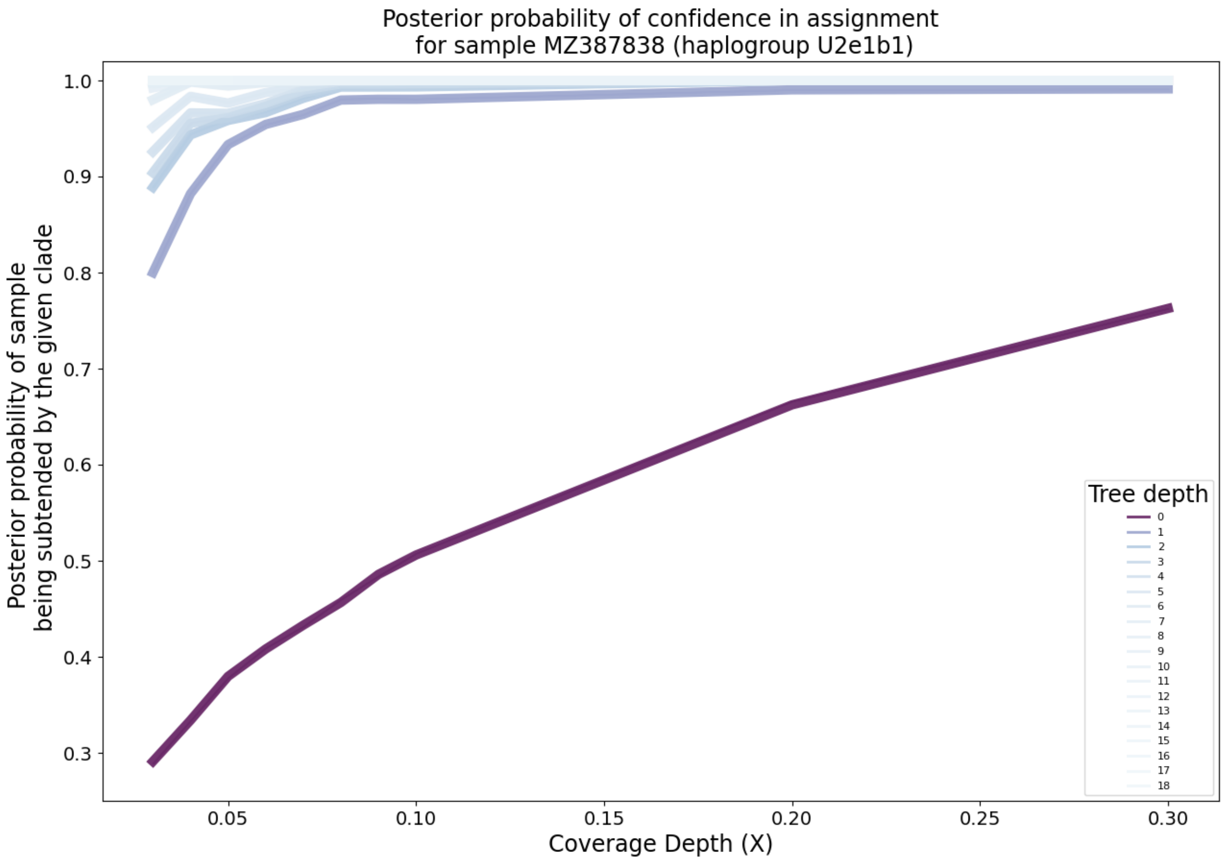

Supplementary Figure 26: **Clade-level posterior probabilities of haplogroup assignment on simulated paired-end FASTQ data.** Each lineplot represents the mean over replicates at a fixed depth on the mitochondrial tree. The darker the line, the more basal the haplogroups.

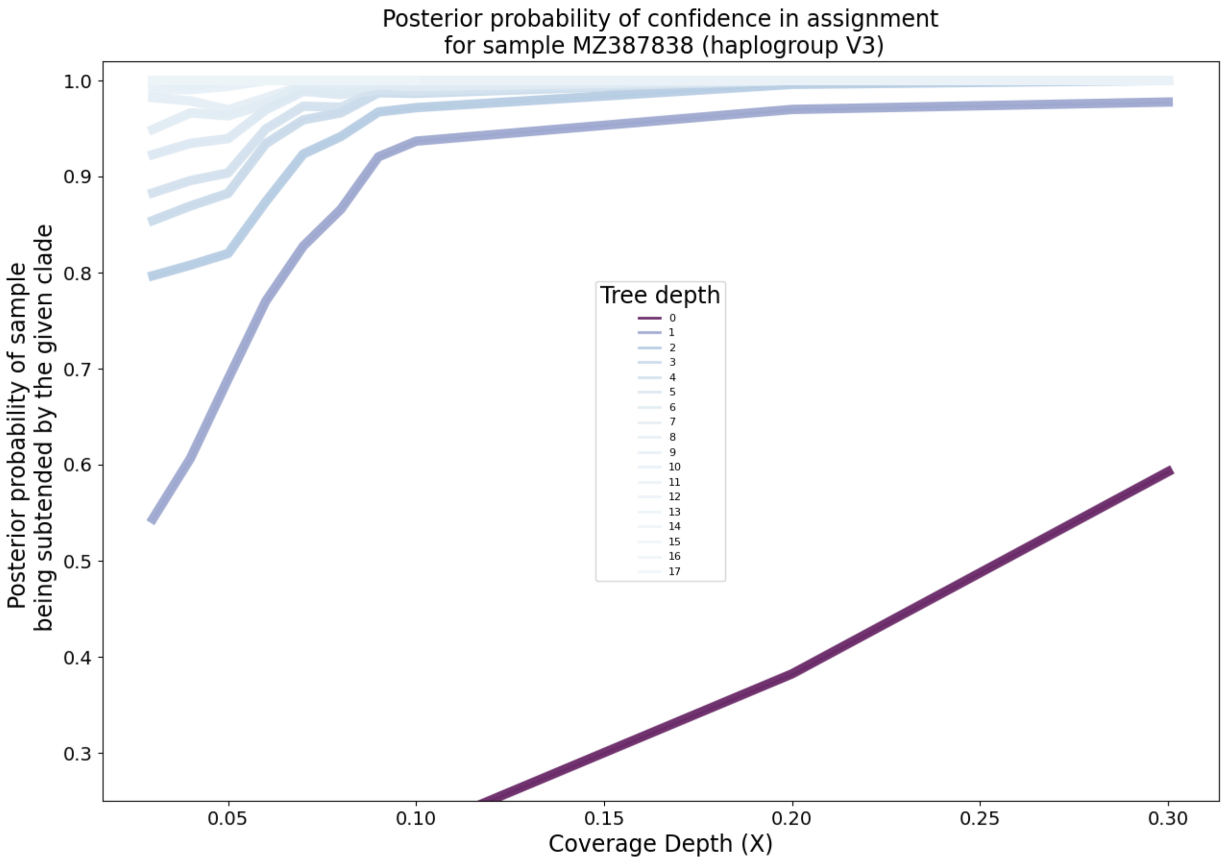

Supplementary Figure 27: **Clade-level posterior probabilities of haplogroup assignment on simulated paired-end FASTQ data.** Each lineplot represents the mean over replicates at a fixed depth on the mitochondrial tree. The darker the line, the more basal the haplogroups.

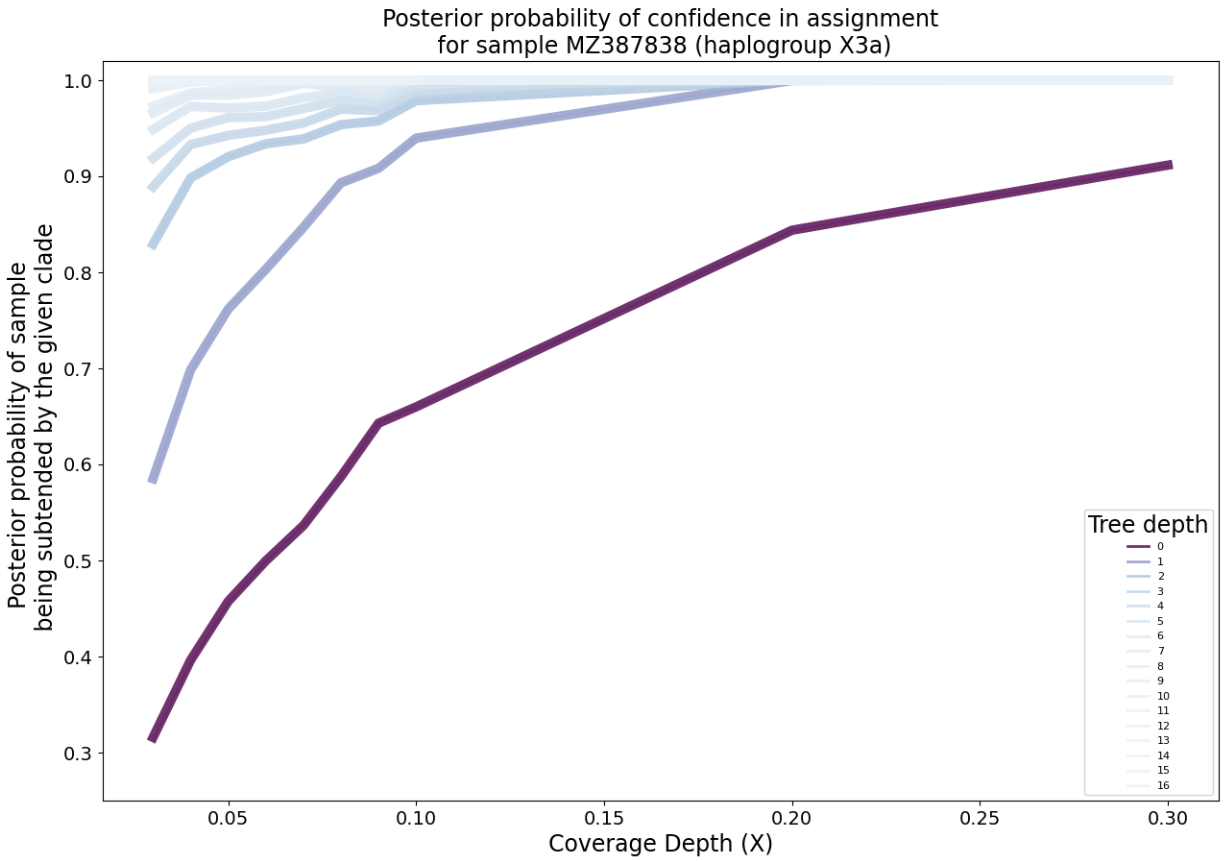

Supplementary Figure 28: **Clade-level posterior probabilities of haplogroup assignment on simulated paired-end FASTQ data.** Each lineplot represents the mean over replicates at a fixed depth on the mitochondrial tree. The darker the line, the more basal the haplogroups.

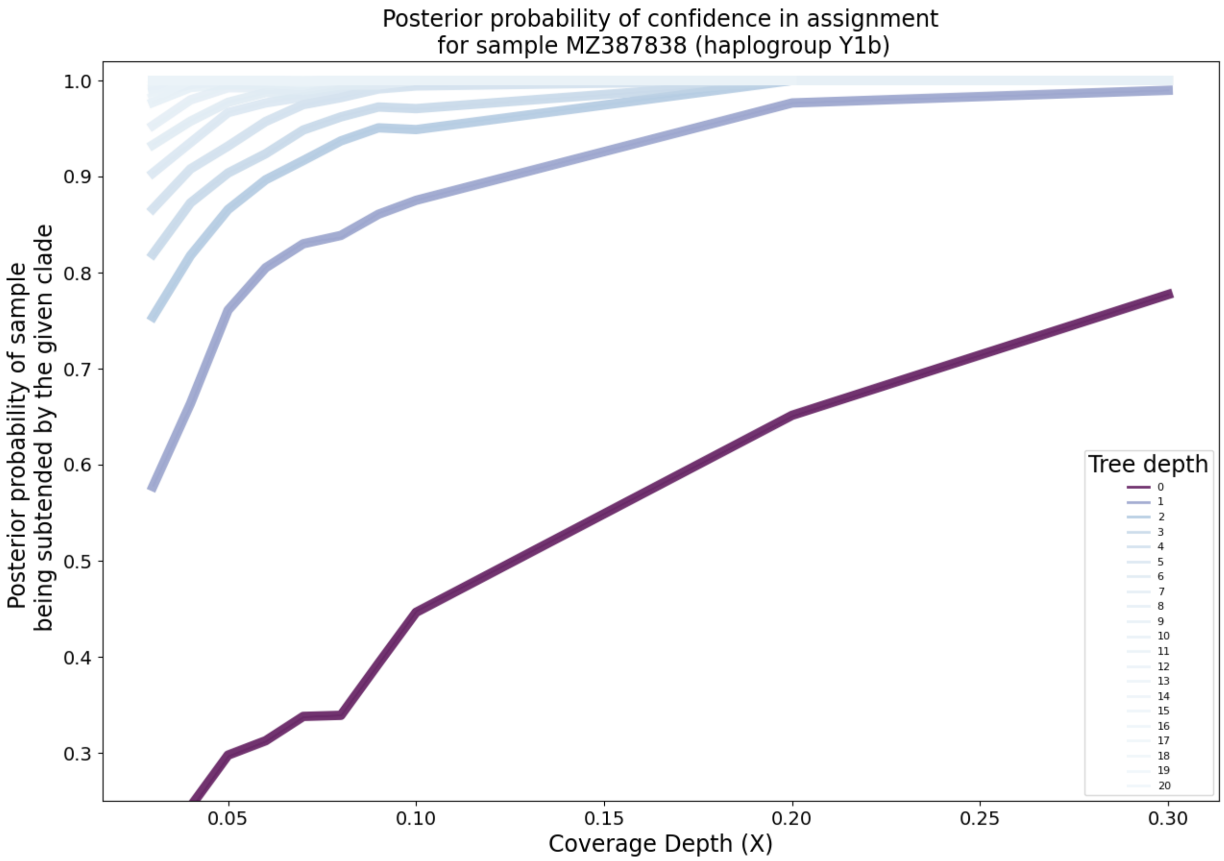

Supplementary Figure 29: **Clade-level posterior probabilities of haplogroup assignment on simulated paired-end FASTQ data.** Each lineplot represents the mean over replicates at a fixed depth on the mitochondrial tree. The darker the line, the more basal the haplogroups.

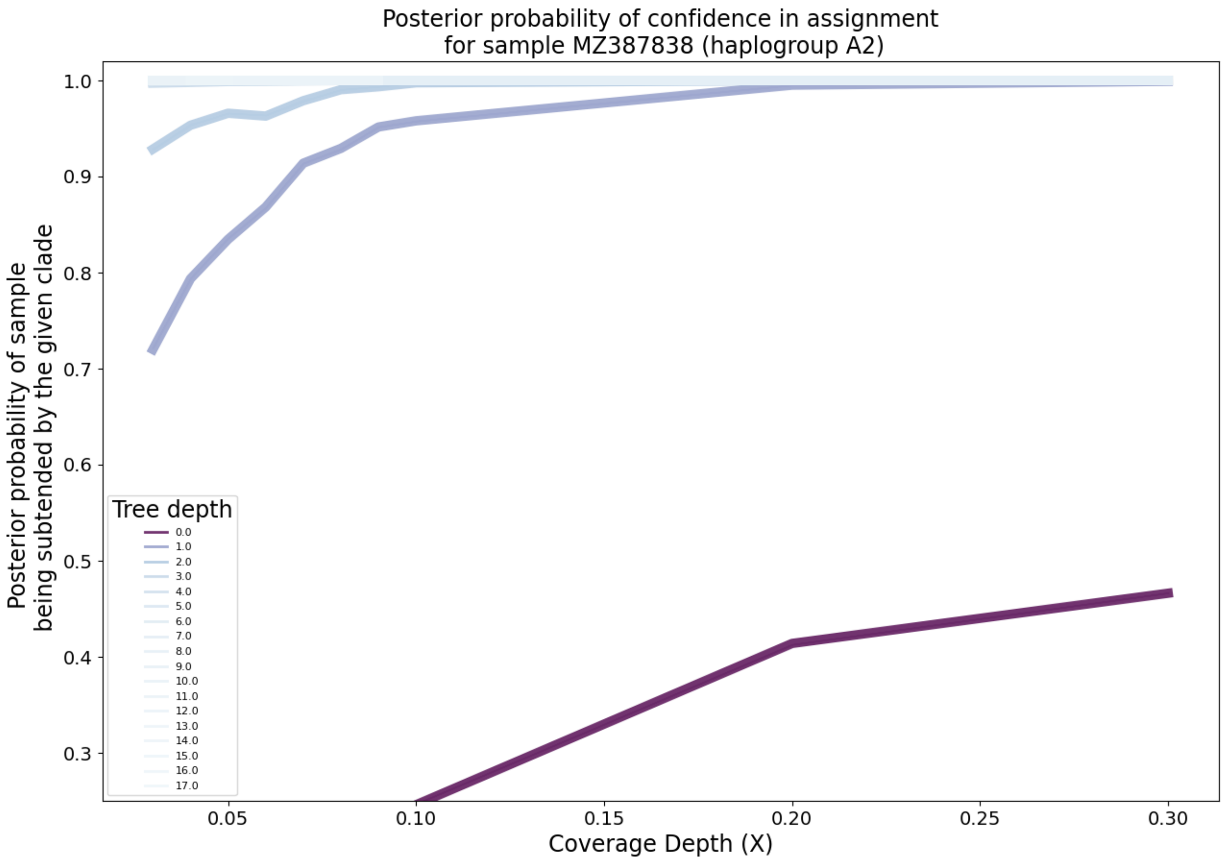

Supplementary Figure 30: **Clade-level posterior probabilities of haplogroup assignment on simulated paired-end FASTQ data with added NuMT reads.** NuMT reads were included at a rate of one in 200. Each lineplot represents the mean over replicates at a fixed depth on the mitochondrial tree. The darker the line, the more basal the haplogroups.

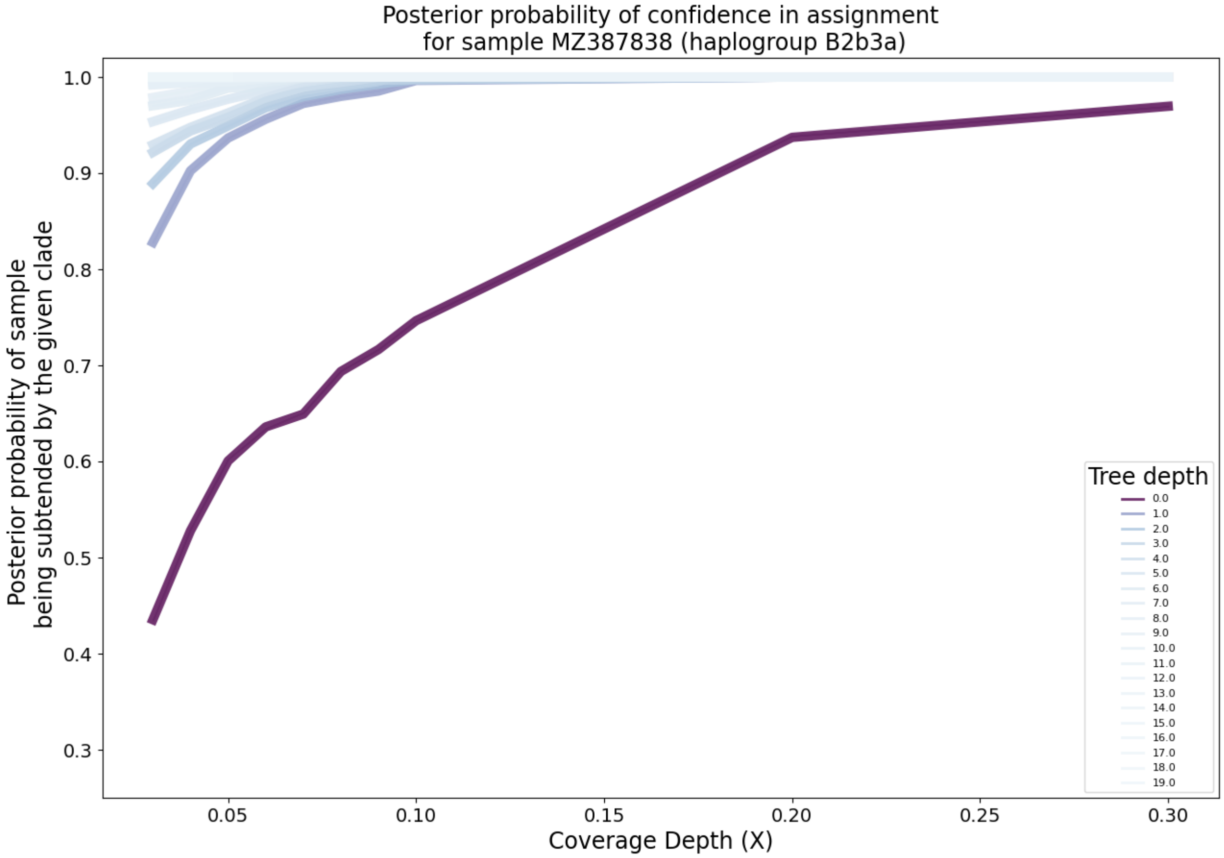

Supplementary Figure 31: **Clade-level posterior probabilities of haplogroup assignment on simulated paired-end FASTQ data with added NuMT reads (CONTINUED).** NuMT reads were included at a rate of one in 200. Each lineplot represents the mean over replicates at a fixed depth on the mitochondrial tree. The darker the line, the more basal the haplogroups.

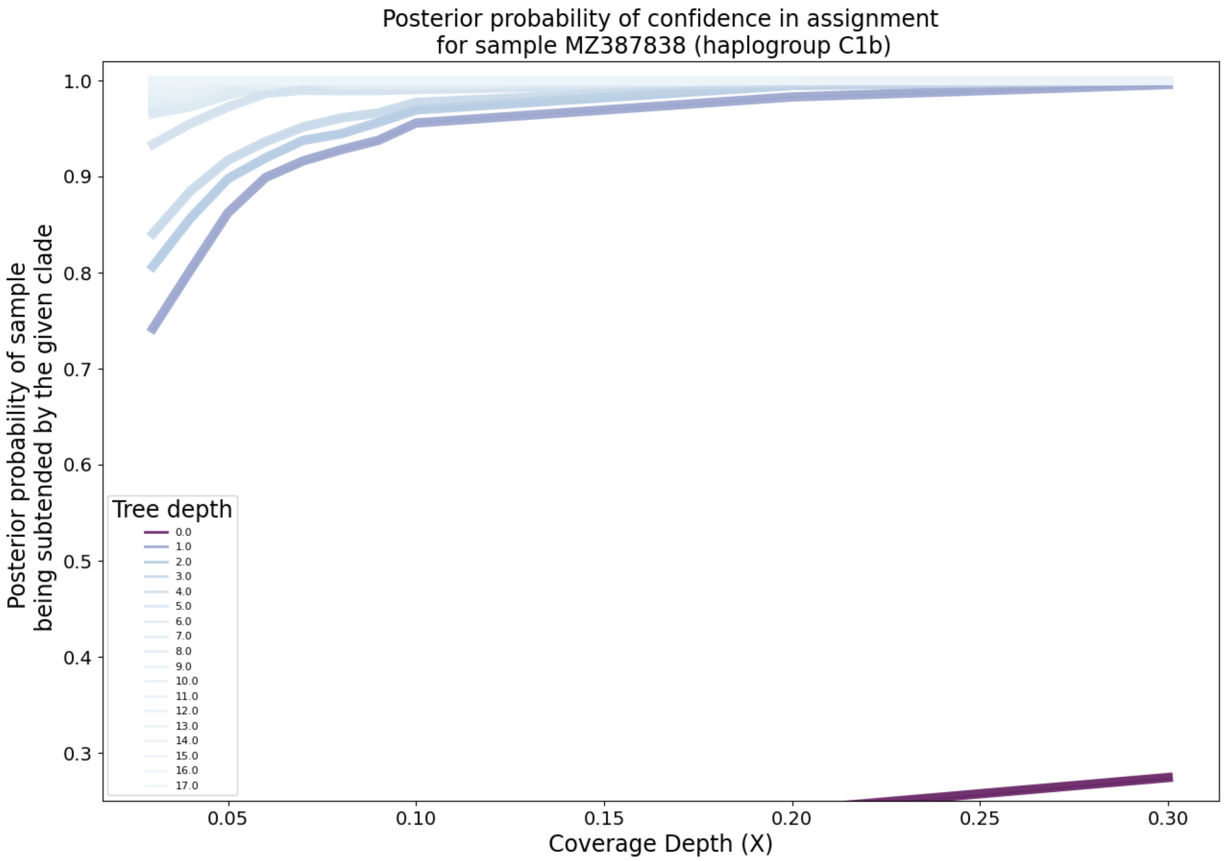

Supplementary Figure 32: **Clade-level posterior probabilities of haplogroup assignment on simulated paired-end FASTQ data with added NuMT reads (CONTINUED).** NuMT reads were included at a rate of one in 200. Each lineplot represents the mean over replicates at a fixed depth on the mitochondrial tree. The darker the line, the more basal the haplogroups.

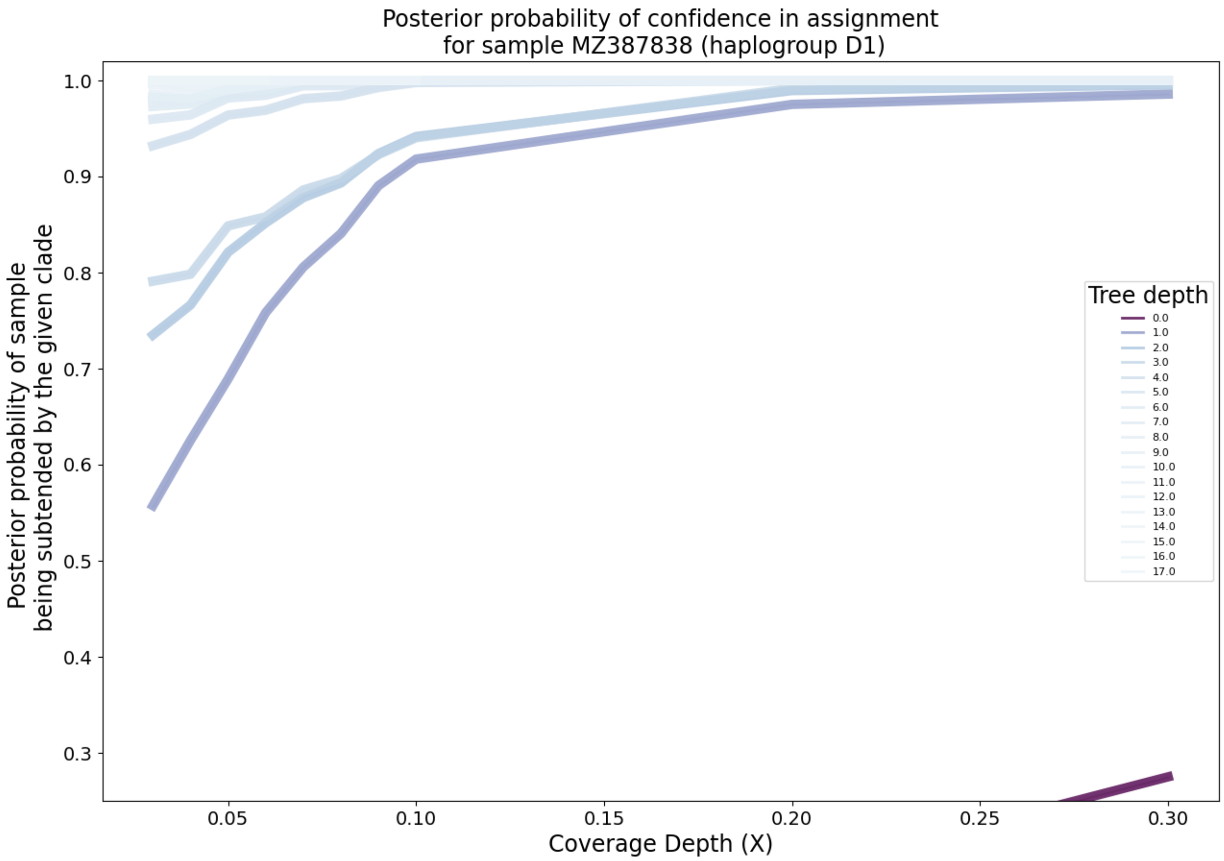

Supplementary Figure 33: **Clade-level posterior probabilities of haplogroup assignment on simulated paired-end FASTQ data with added NuMT reads (CONTINUED).** NuMT reads were included at a rate of one in 200. Each lineplot represents the mean over replicates at a fixed depth on the mitochondrial tree. The darker the line, the more basal the haplogroups.

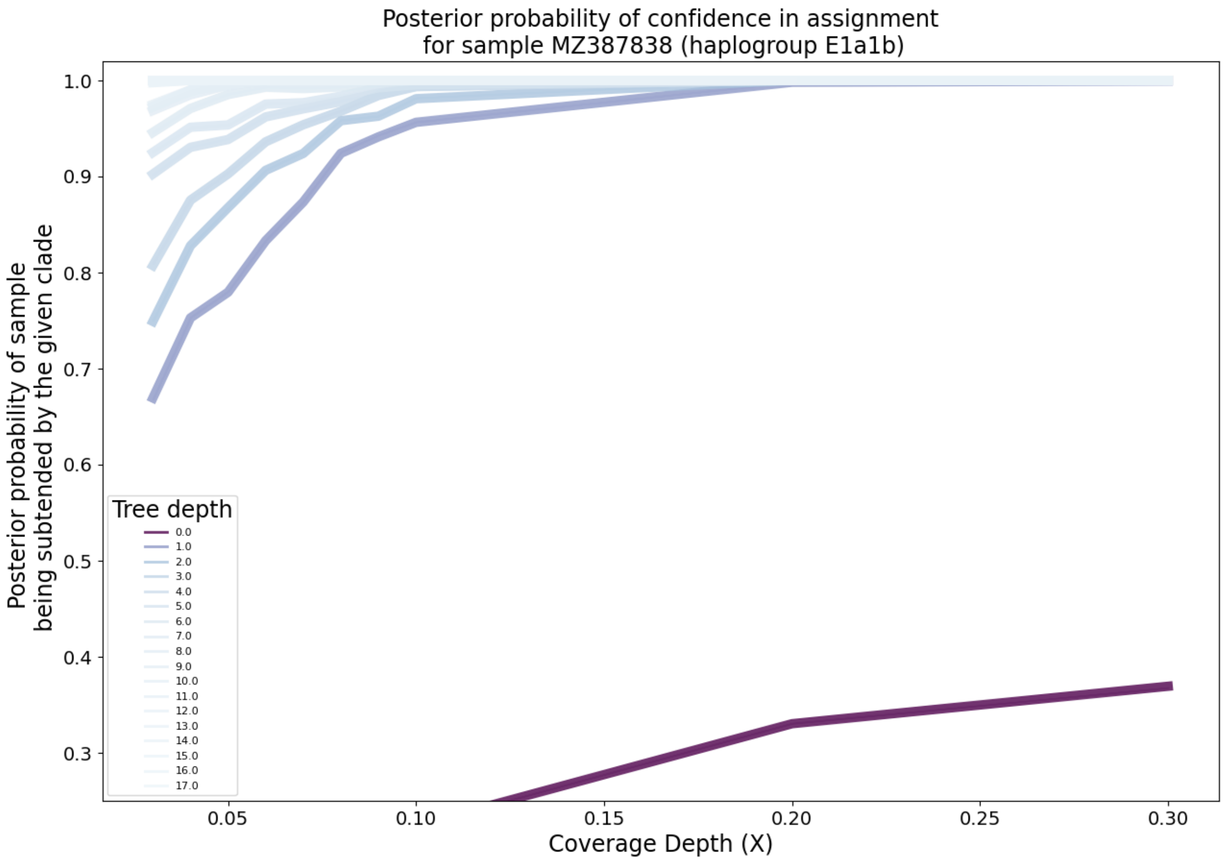

Supplementary Figure 34: **Clade-level posterior probabilities of haplogroup assignment on simulated paired-end FASTQ data with added NuMT reads (CONTINUED).** NuMT reads were included at a rate of one in 200. Each lineplot represents the mean over replicates at a fixed depth on the mitochondrial tree. The darker the line, the more basal the haplogroups.

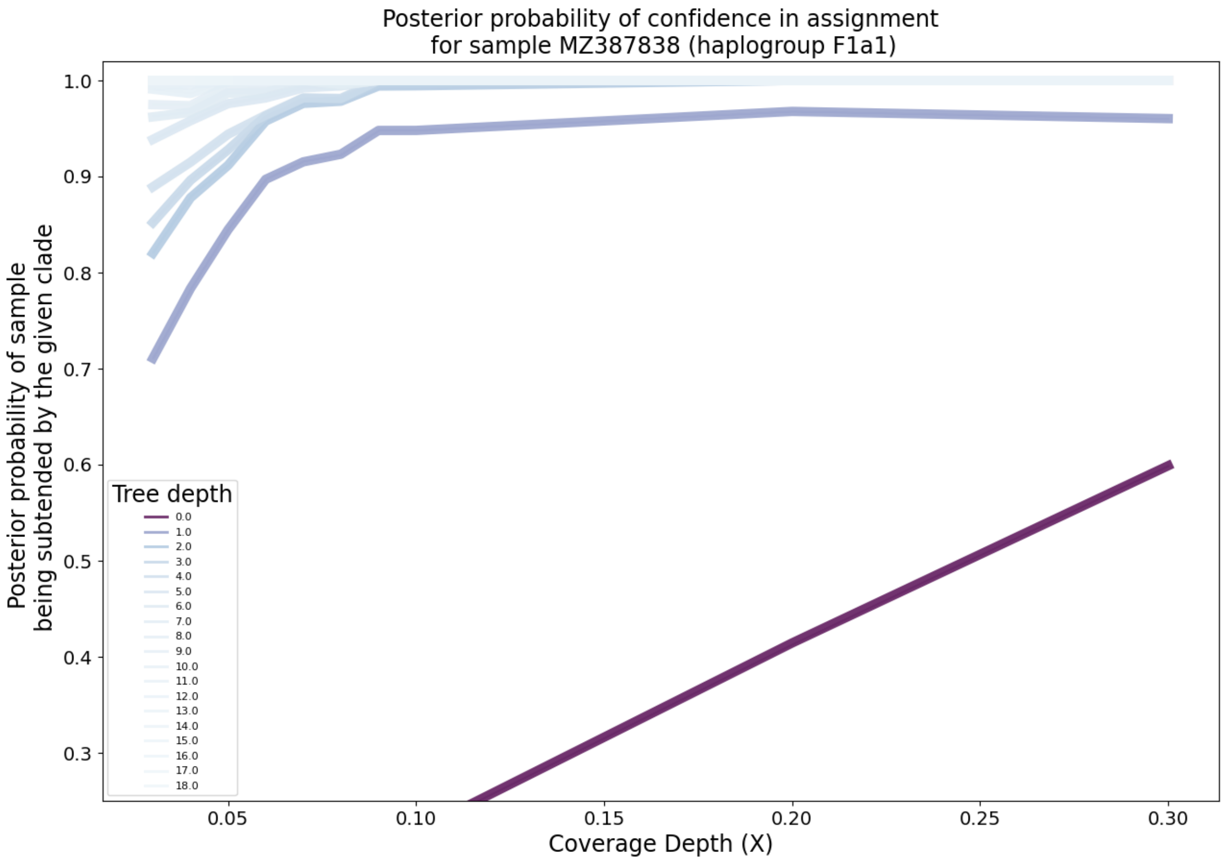

Supplementary Figure 35: **Clade-level posterior probabilities of haplogroup assignment on simulated paired-end FASTQ data with added NuMT reads (CONTINUED).** NuMT reads were included at a rate of one in 200. Each lineplot represents the mean over replicates at a fixed depth on the mitochondrial tree. The darker the line, the more basal the haplogroups.

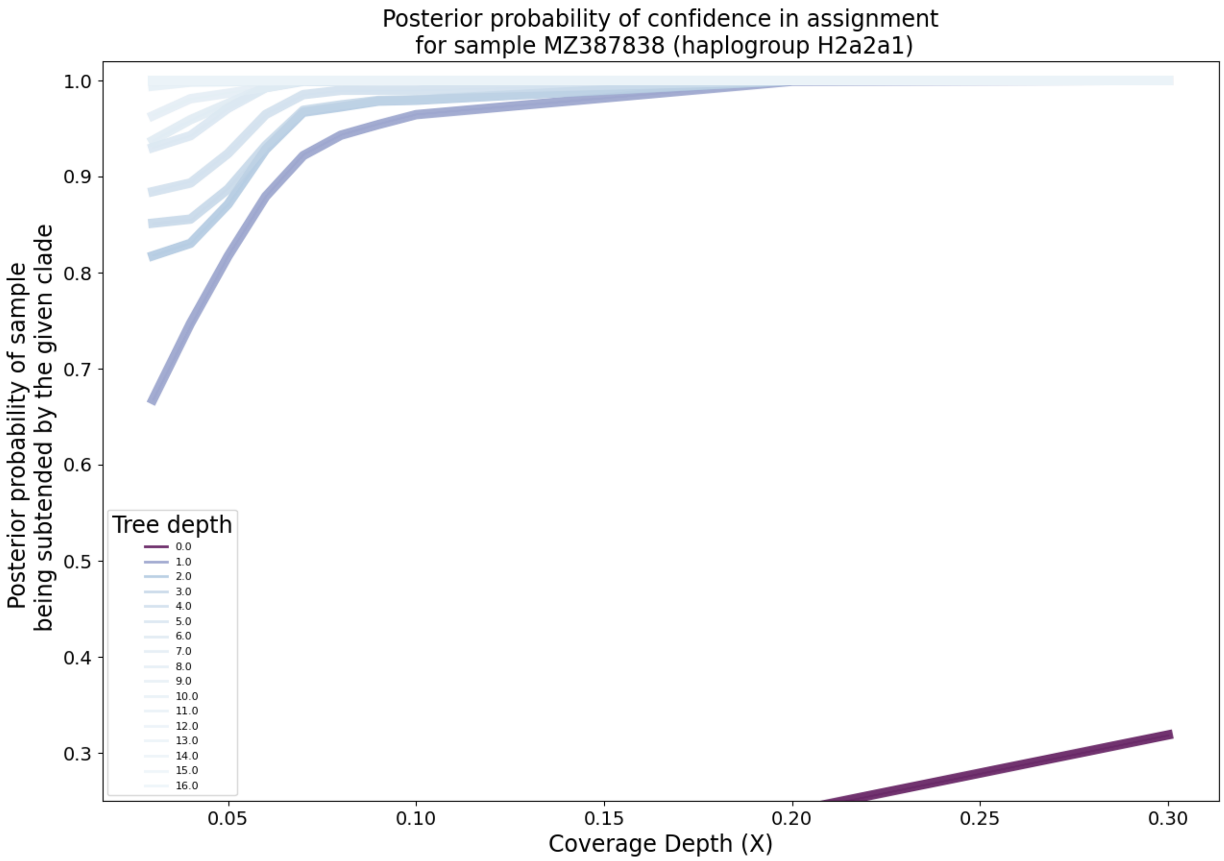

Supplementary Figure 36: **Clade-level posterior probabilities of haplogroup assignment on simulated paired-end FASTQ data with added NuMT reads (CONTINUED).** NuMT reads were included at a rate of one in 200. Each lineplot represents the mean over replicates at a fixed depth on the mitochondrial tree. The darker the line, the more basal the haplogroups.

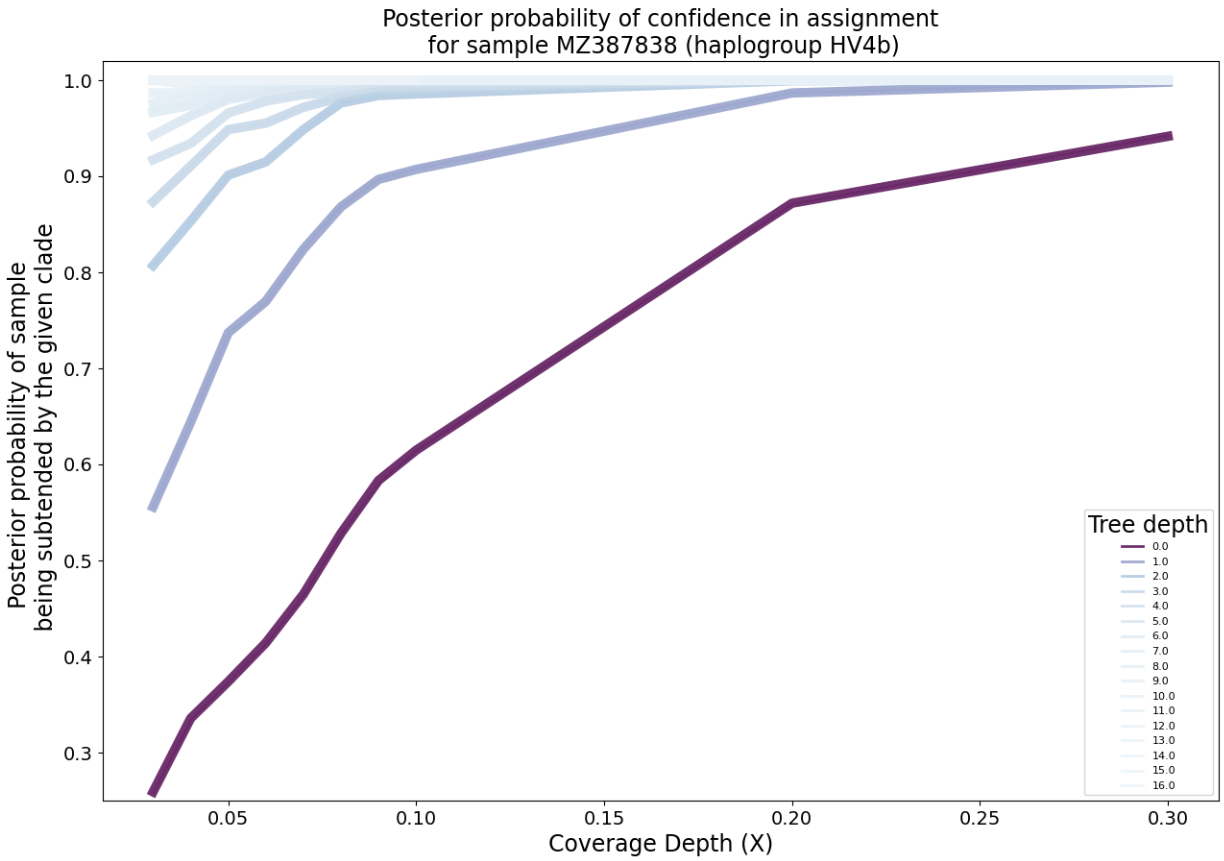

Supplementary Figure 37: **Clade-level posterior probabilities of haplogroup assignment on simulated paired-end FASTQ data with added NuMT reads (CONTINUED).** NuMT reads were included at a rate of one in 200. Each lineplot represents the mean over replicates at a fixed depth on the mitochondrial tree. The darker the line, the more basal the haplogroups.

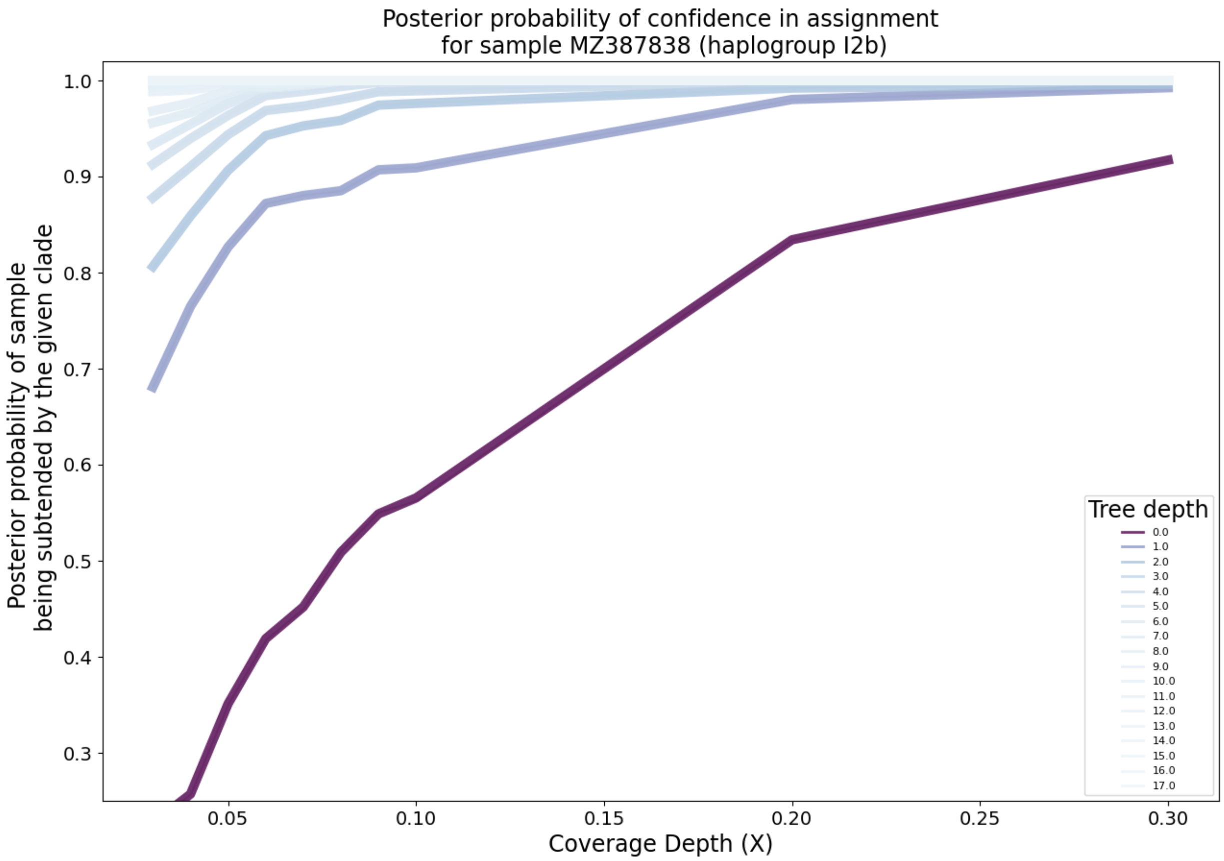

Supplementary Figure 38: **Clade-level posterior probabilities of haplogroup assignment on simulated paired-end FASTQ data with added NuMT reads (CONTINUED).** NuMT reads were included at a rate of one in 200. Each lineplot represents the mean over replicates at a fixed depth on the mitochondrial tree. The darker the line, the more basal the haplogroups.

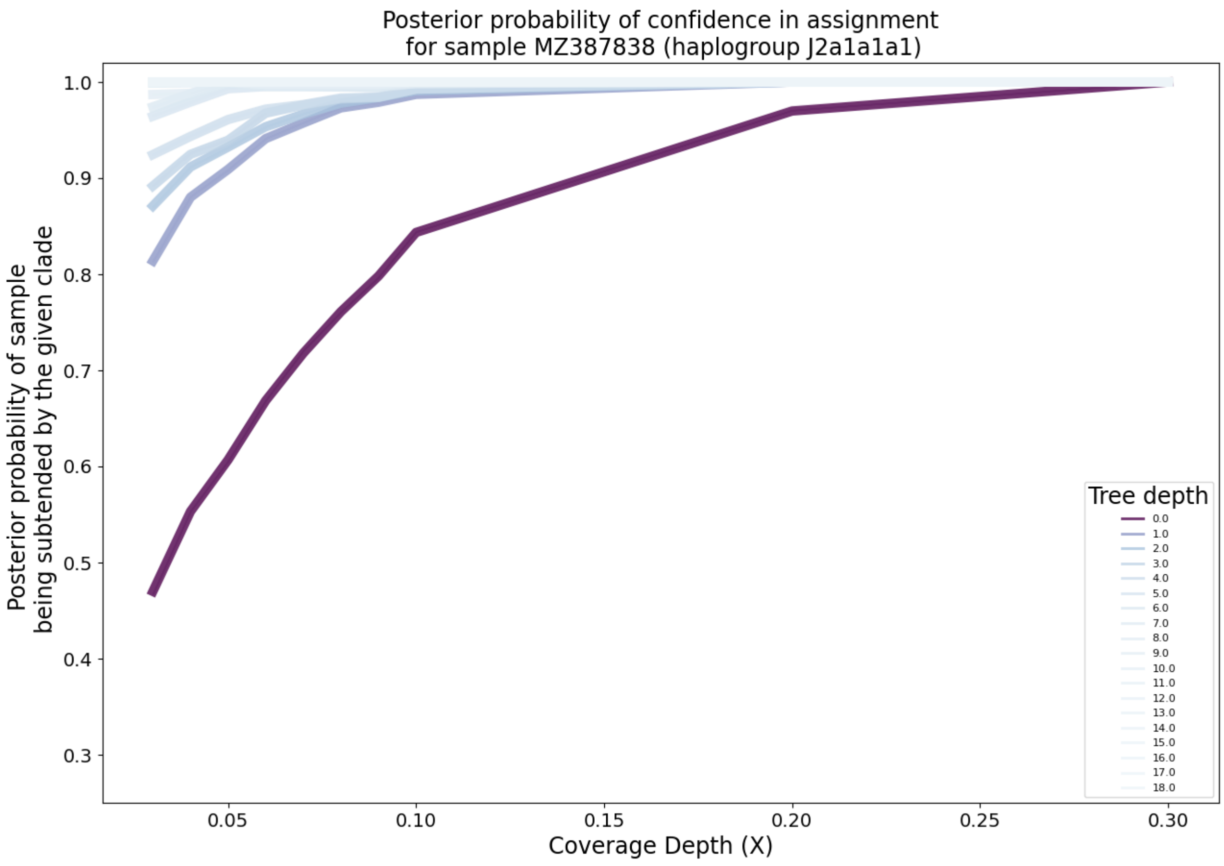

Supplementary Figure 39: **Clade-level posterior probabilities of haplogroup assignment on simulated paired-end FASTQ data with added NuMT reads (CONTINUED).** NuMT reads were included at a rate of one in 200. Each lineplot represents the mean over replicates at a fixed depth on the mitochondrial tree. The darker the line, the more basal the haplogroups.

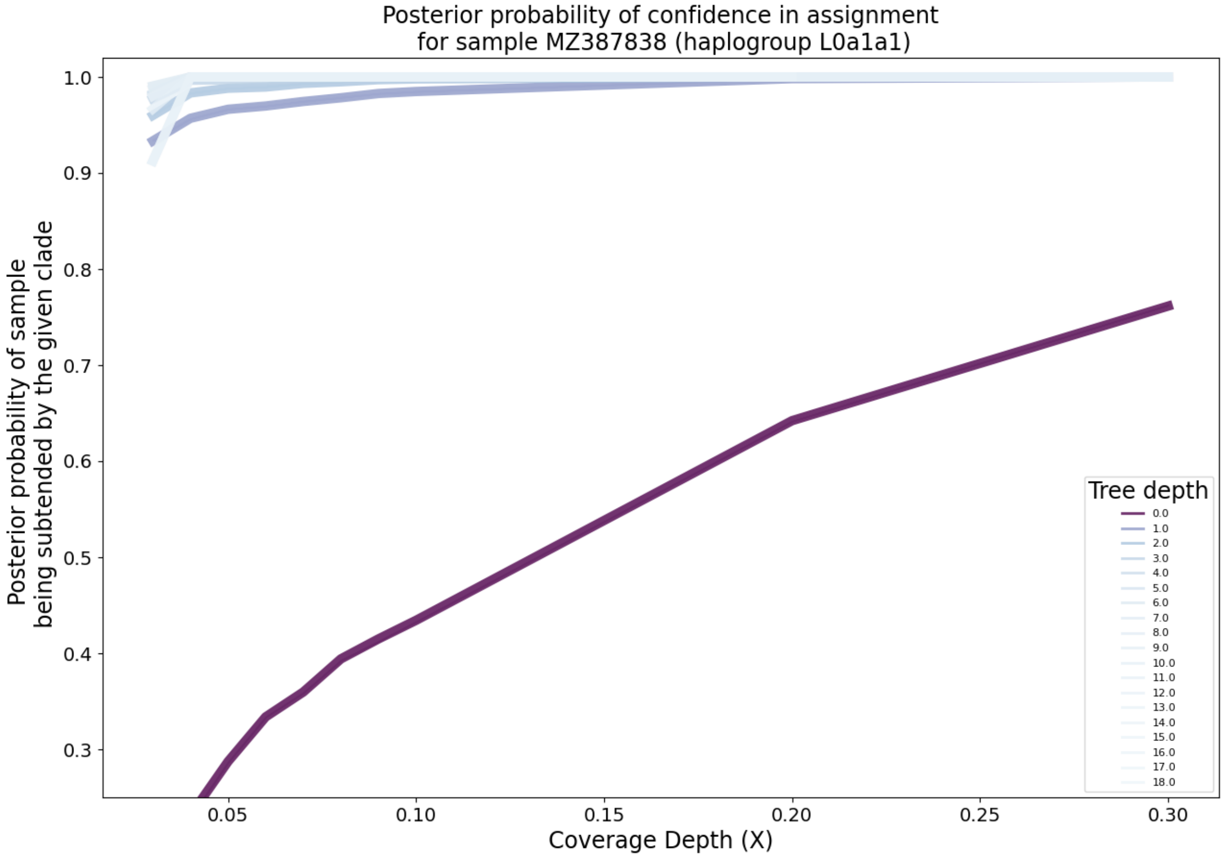

Supplementary Figure 40: **Clade-level posterior probabilities of haplogroup assignment on simulated paired-end FASTQ data with added NuMT reads (CONTINUED).** NuMT reads were included at a rate of one in 200. Each lineplot represents the mean over replicates at a fixed depth on the mitochondrial tree. The darker the line, the more basal the haplogroups.

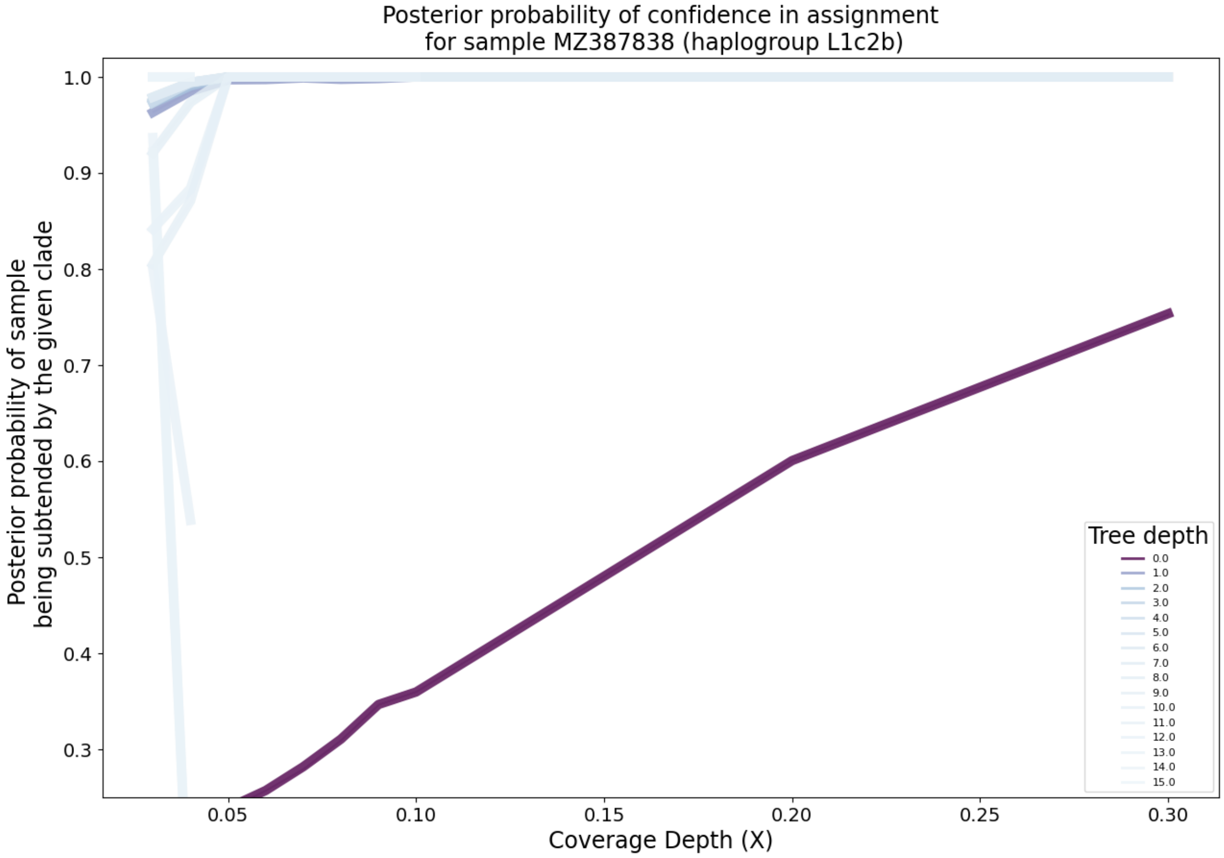

Supplementary Figure 41: **Clade-level posterior probabilities of haplogroup assignment on simulated paired-end FASTQ data with added NuMT reads (CONTINUED).** NuMT reads were included at a rate of one in 200. Each lineplot represents the mean over replicates at a fixed depth on the mitochondrial tree. The darker the line, the more basal the haplogroups.

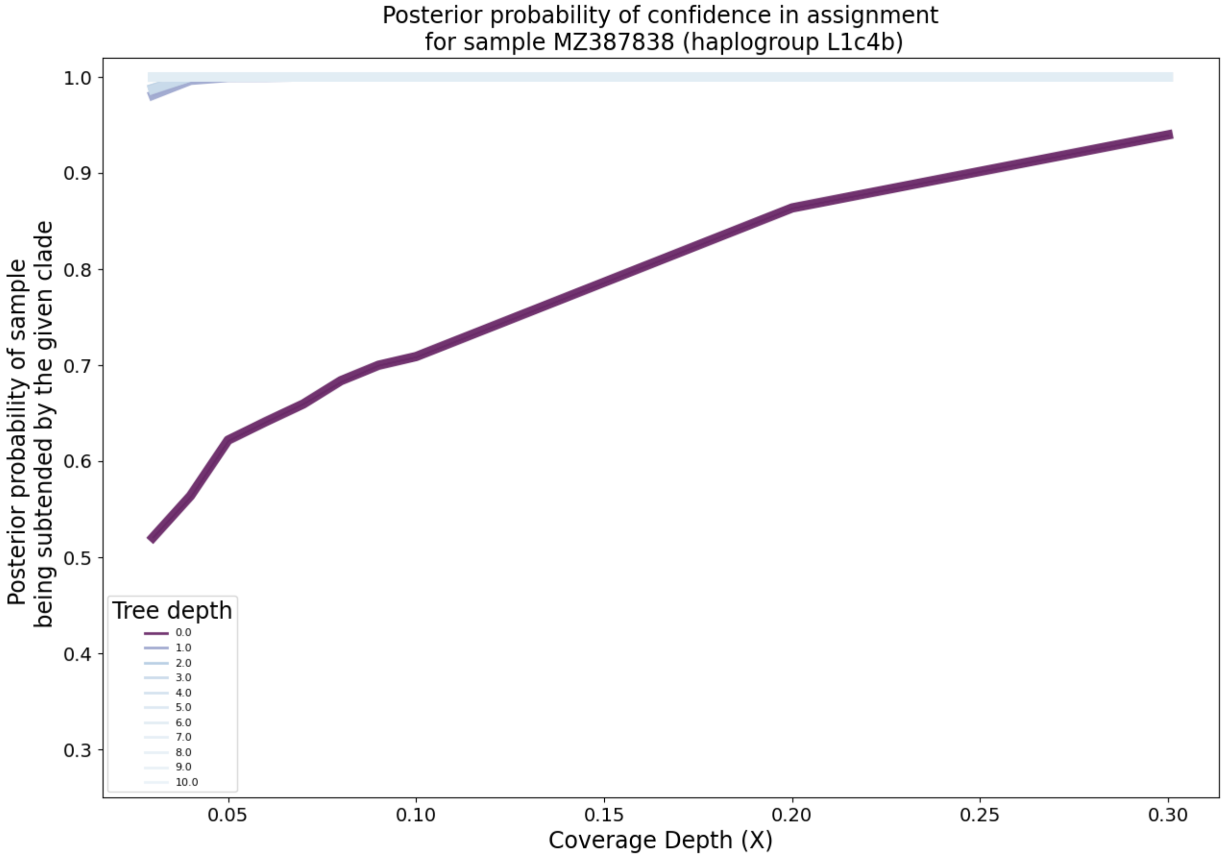

Supplementary Figure 42: **Clade-level posterior probabilities of haplogroup assignment on simulated paired-end FASTQ data with added NuMT reads (CONTINUED).** NuMT reads were included at a rate of one in 200. Each lineplot represents the mean over replicates at a fixed depth on the mitochondrial tree. The darker the line, the more basal the haplogroups.

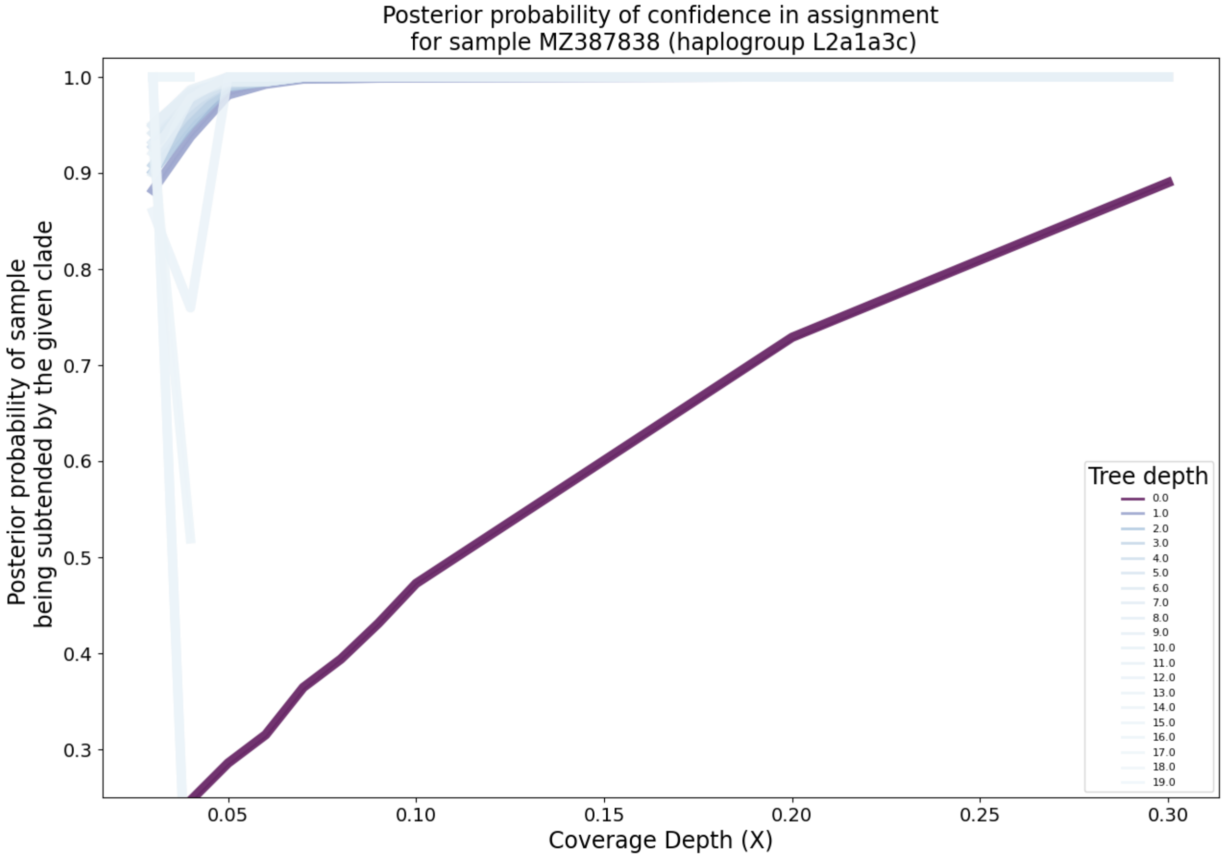

Supplementary Figure 43: **Clade-level posterior probabilities of haplogroup assignment on simulated paired-end FASTQ data with added NuMT reads (CONTINUED).** NuMT reads were included at a rate of one in 200. Each lineplot represents the mean over replicates at a fixed depth on the mitochondrial tree. The darker the line, the more basal the haplogroups.

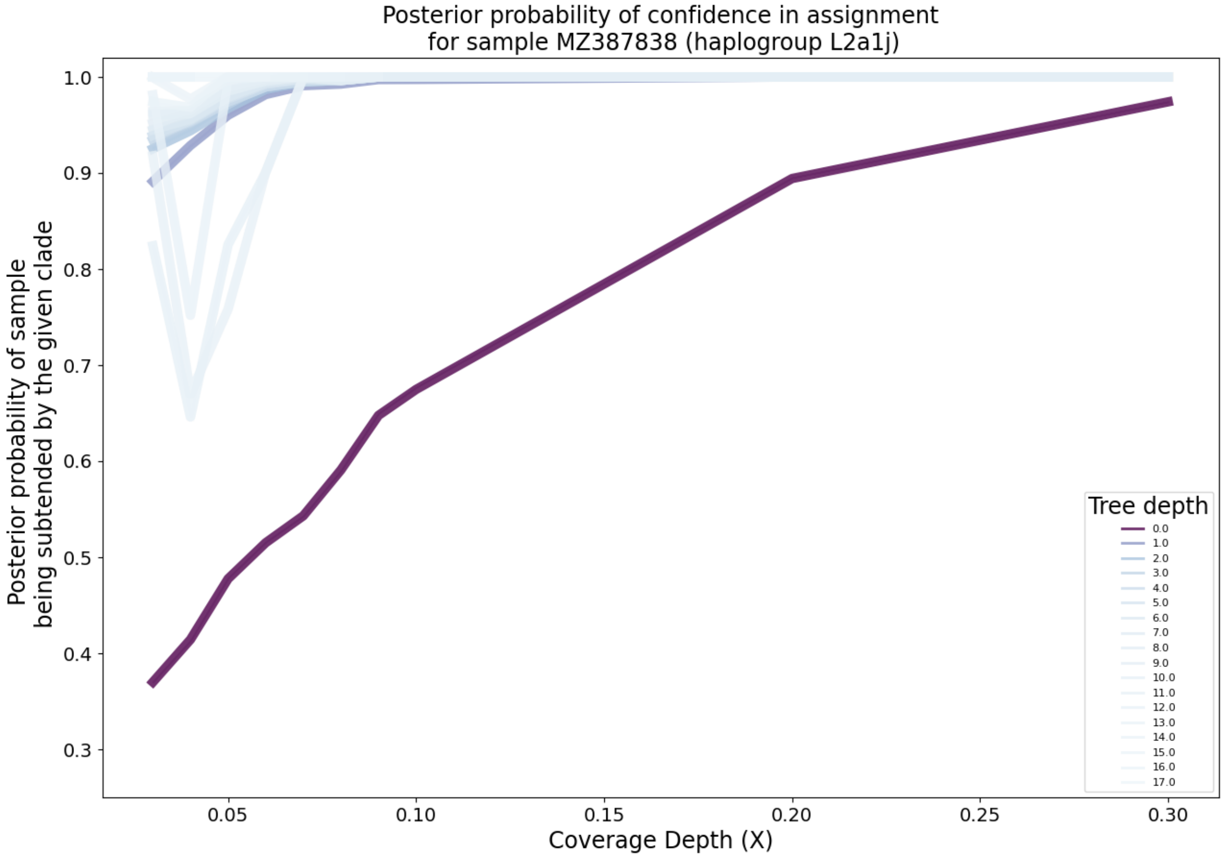

Supplementary Figure 44: **Clade-level posterior probabilities of haplogroup assignment on simulated paired-end FASTQ data with added NuMT reads (CONTINUED).** NuMT reads were included at a rate of one in 200. Each lineplot represents the mean over replicates at a fixed depth on the mitochondrial tree. The darker the line, the more basal the haplogroups.

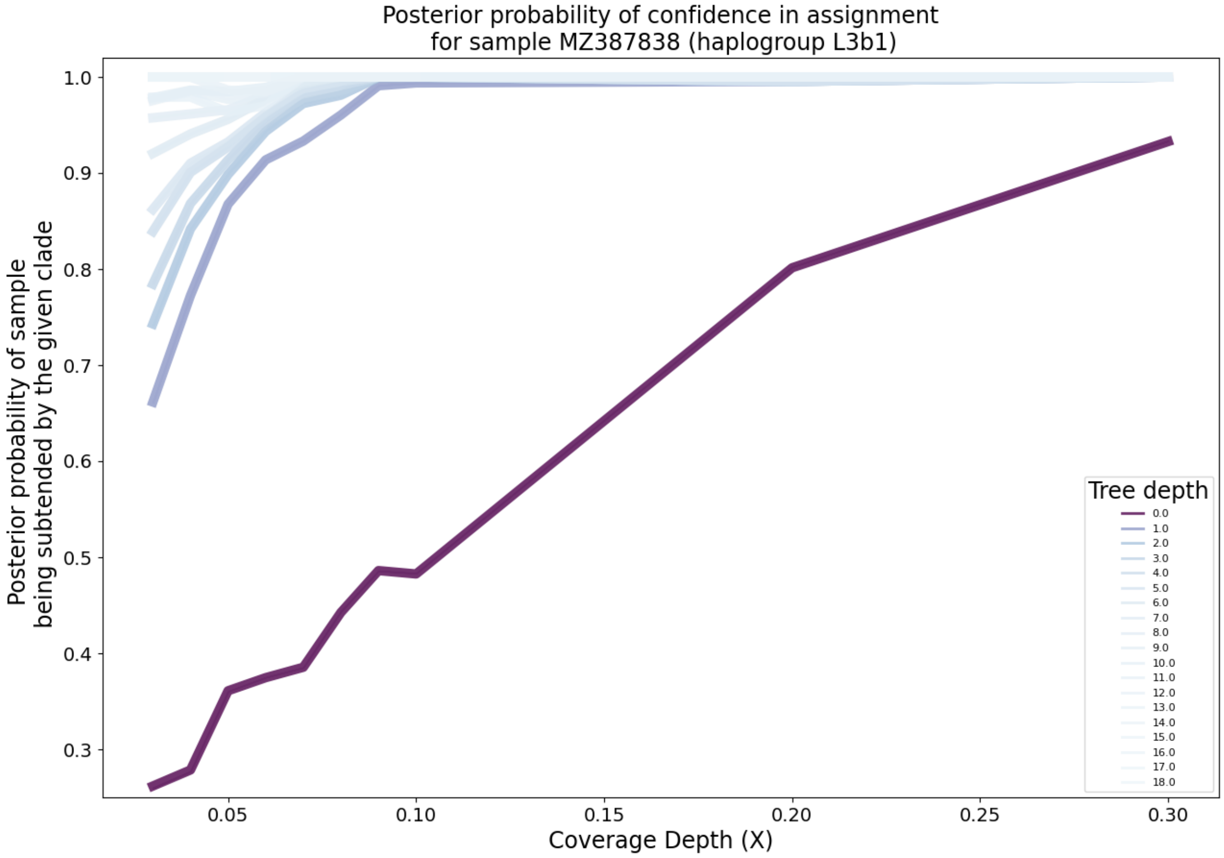

Supplementary Figure 45: **Clade-level posterior probabilities of haplogroup assignment on simulated paired-end FASTQ data with added NuMT reads (CONTINUED).** NuMT reads were included at a rate of one in 200. Each lineplot represents the mean over replicates at a fixed depth on the mitochondrial tree. The darker the line, the more basal the haplogroups.

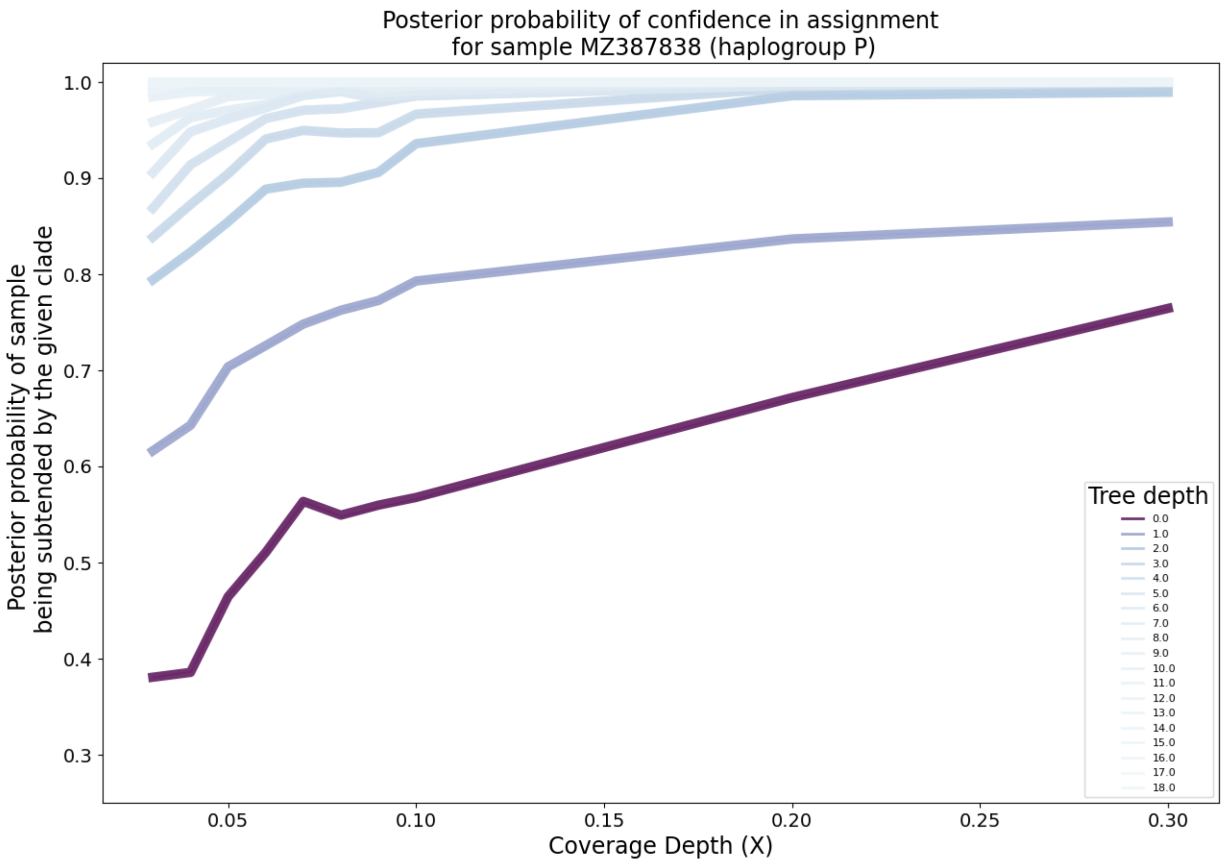

Supplementary Figure 46: **Clade-level posterior probabilities of haplogroup assignment on simulated paired-end FASTQ data with added NuMT reads (CONTINUED).** NuMT reads were included at a rate of one in 200. Each lineplot represents the mean over replicates at a fixed depth on the mitochondrial tree. The darker the line, the more basal the haplogroups.

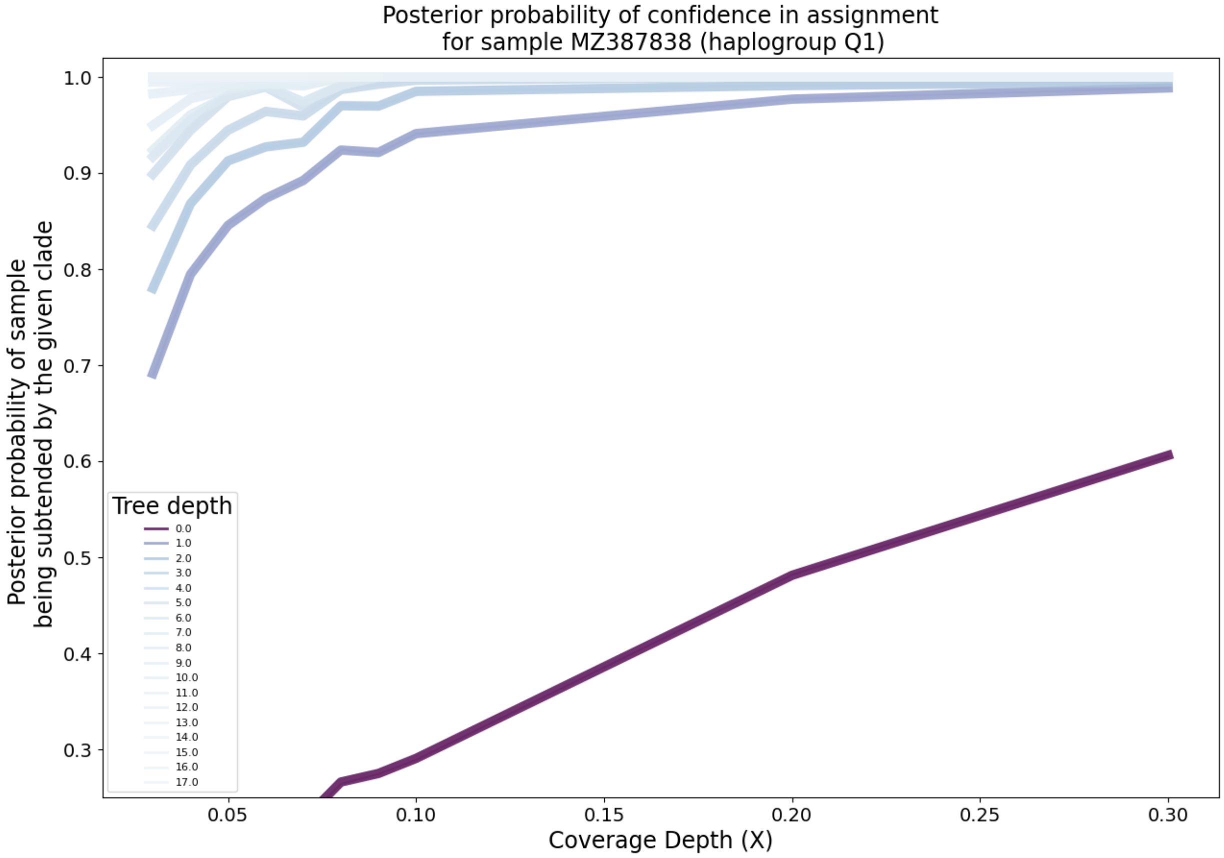

Supplementary Figure 47: **Clade-level posterior probabilities of haplogroup assignment on simulated paired-end FASTQ data with added NuMT reads (CONTINUED).** NuMT reads were included at a rate of one in 200. Each lineplot represents the mean over replicates at a fixed depth on the mitochondrial tree. The darker the line, the more basal the haplogroups.

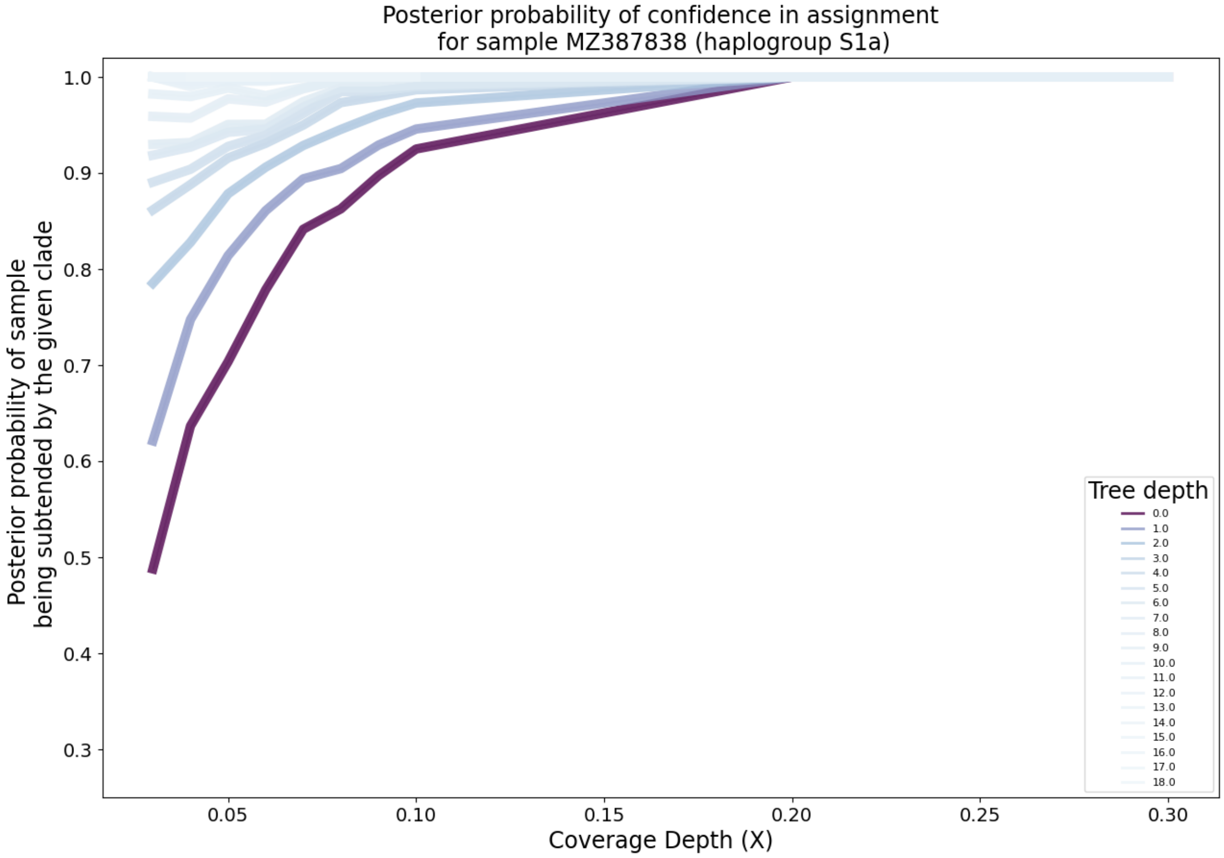

Supplementary Figure 48: **Clade-level posterior probabilities of haplogroup assignment on simulated paired-end FASTQ data with added NuMT reads (CONTINUED).** NuMT reads were included at a rate of one in 200. Each lineplot represents the mean over replicates at a fixed depth on the mitochondrial tree. The darker the line, the more basal the haplogroups.

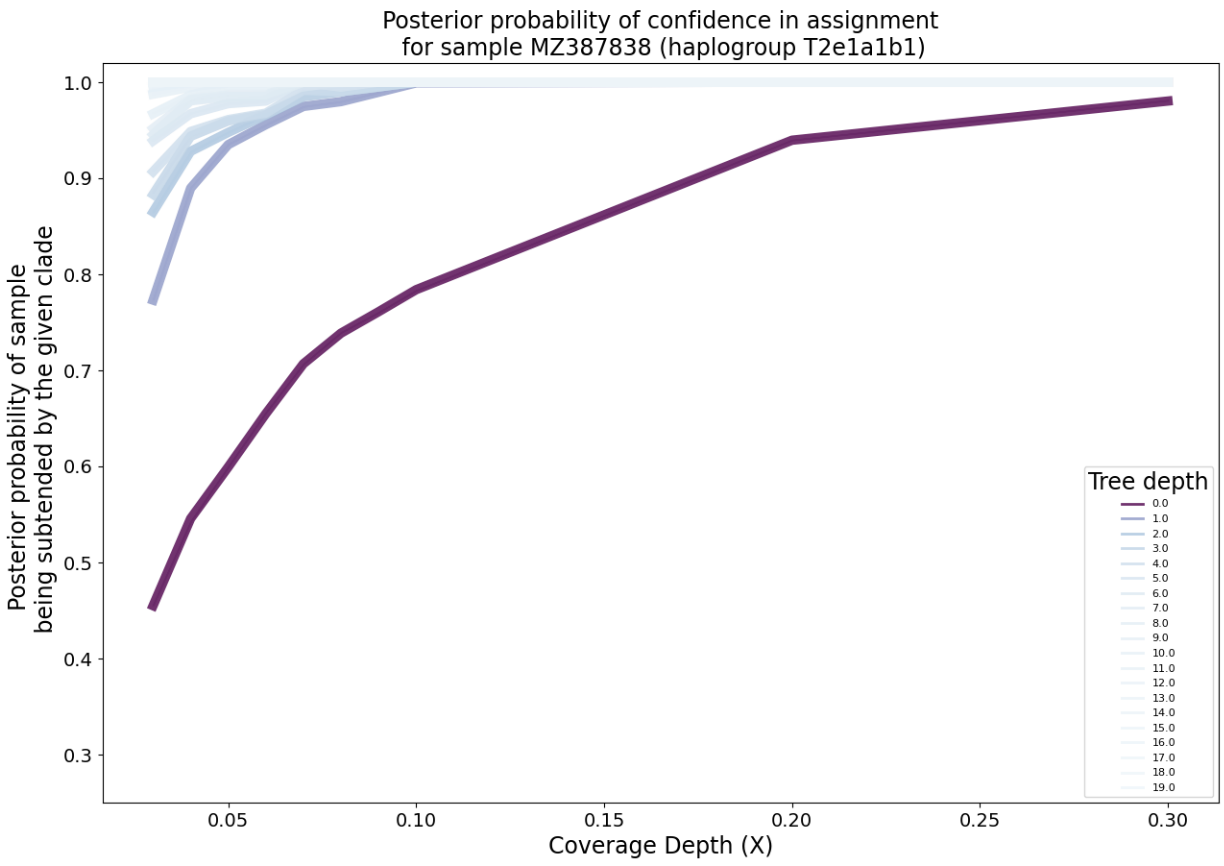

Supplementary Figure 49: **Clade-level posterior probabilities of haplogroup assignment on simulated paired-end FASTQ data with added NuMT reads (CONTINUED).** NuMT reads were included at a rate of one in 200. Each lineplot represents the mean over replicates at a fixed depth on the mitochondrial tree. The darker the line, the more basal the haplogroups.

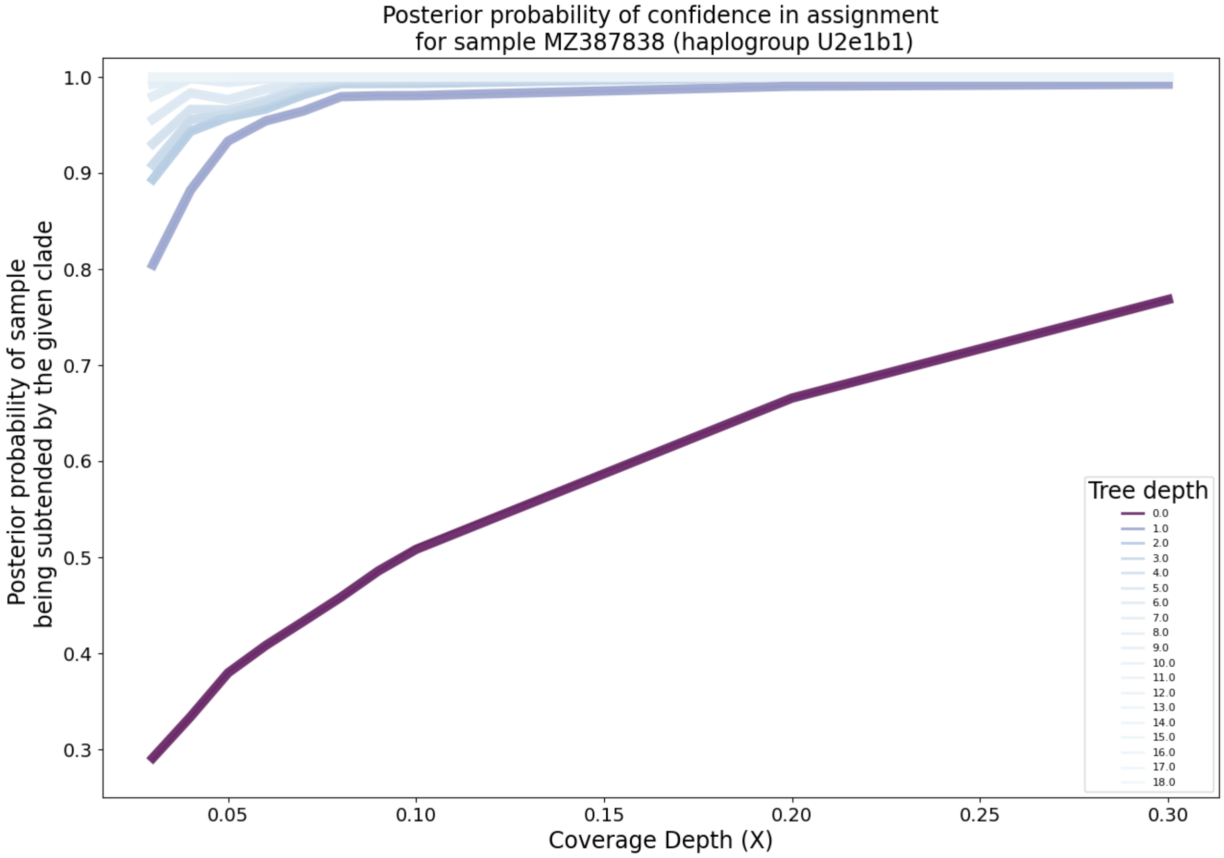

Supplementary Figure 50: **Clade-level posterior probabilities of haplogroup assignment on simulated paired-end FASTQ data with added NuMT reads (CONTINUED).** NuMT reads were included at a rate of one in 200. Each lineplot represents the mean over replicates at a fixed depth on the mitochondrial tree. The darker the line, the more basal the haplogroups.

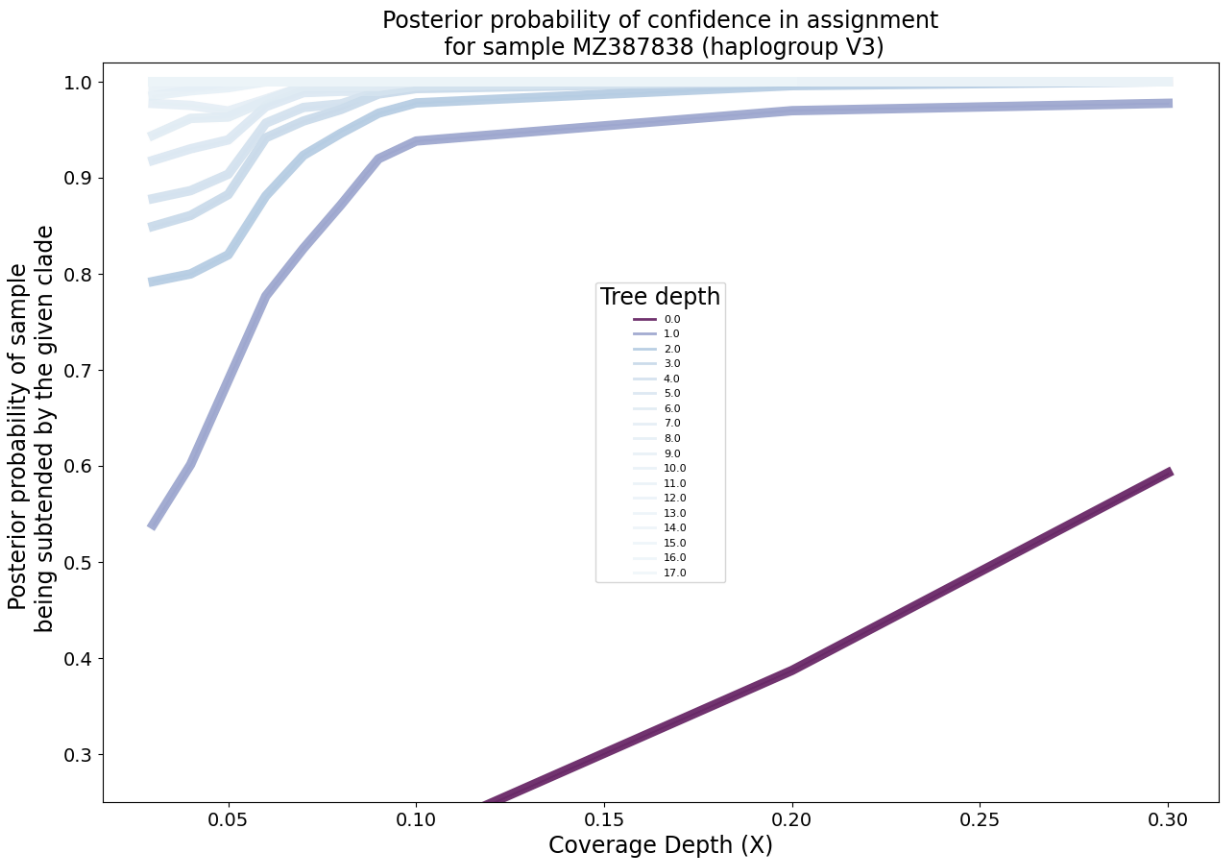

Supplementary Figure 51: **Clade-level posterior probabilities of haplogroup assignment on simulated paired-end FASTQ data with added NuMT reads (CONTINUED).** NuMT reads were included at a rate of one in 200. Each lineplot represents the mean over replicates at a fixed depth on the mitochondrial tree. The darker the line, the more basal the haplogroups.

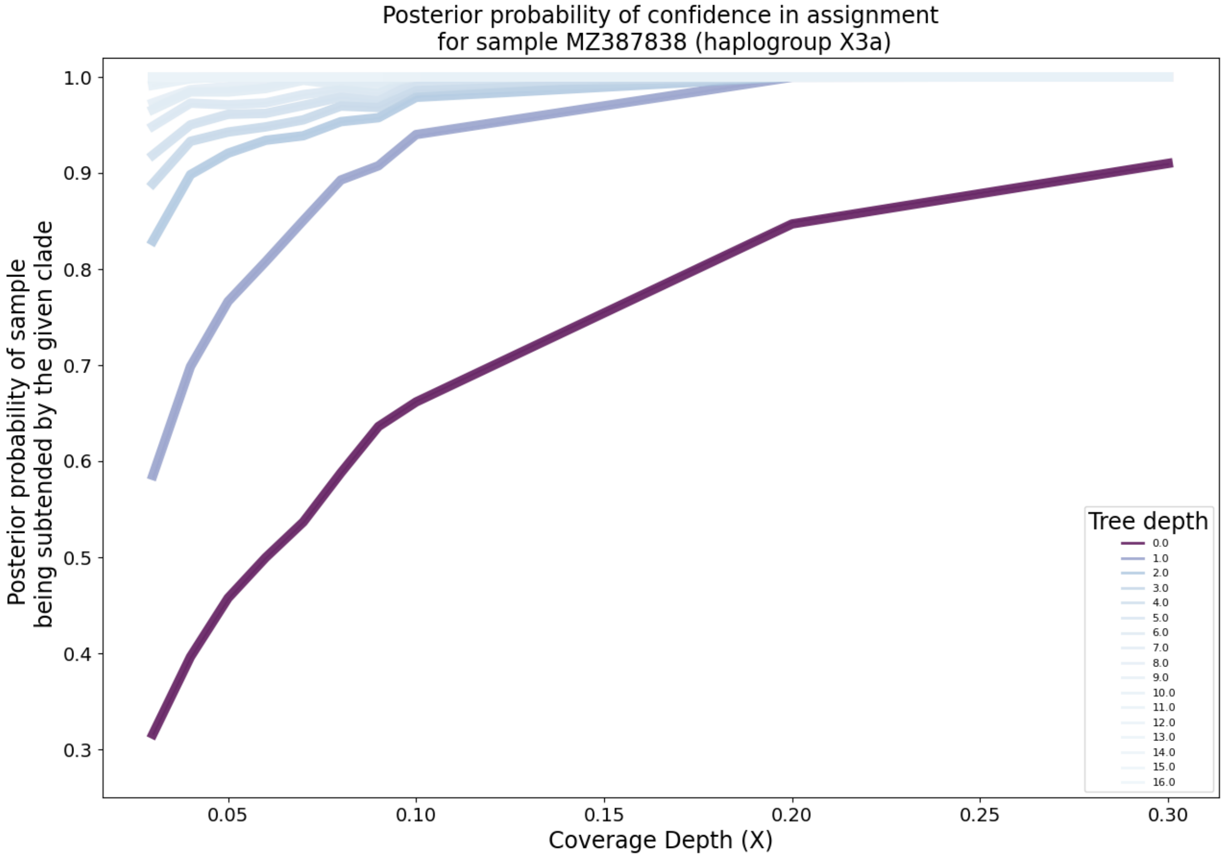

Supplementary Figure 52: **Clade-level posterior probabilities of haplogroup assignment on simulated paired-end FASTQ data with added NuMT reads (CONTINUED).** NuMT reads were included at a rate of one in 200. Each lineplot represents the mean over replicates at a fixed depth on the mitochondrial tree. The darker the line, the more basal the haplogroups.

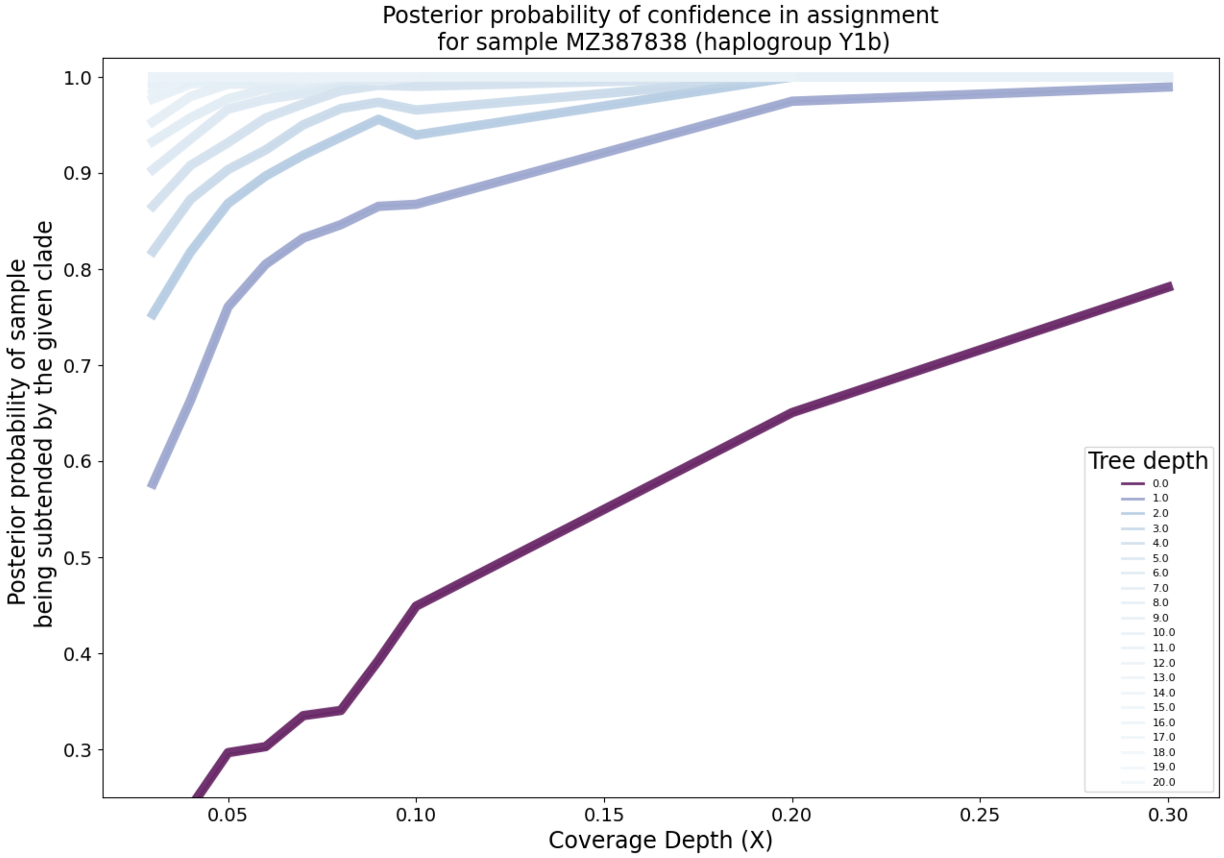

Supplementary Figure 53: **Clade-level posterior probabilities of haplogroup assignment on simulated paired-end FASTQ data with added NuMT reads (CONTINUED).** NuMT reads were included at a rate of one in 200. Each lineplot represents the mean over replicates at a fixed depth on the mitochondrial tree. The darker the line, the more basal the haplogroups.

#### 2.4 Empirical Paired-end FASTQ Posterior Plots

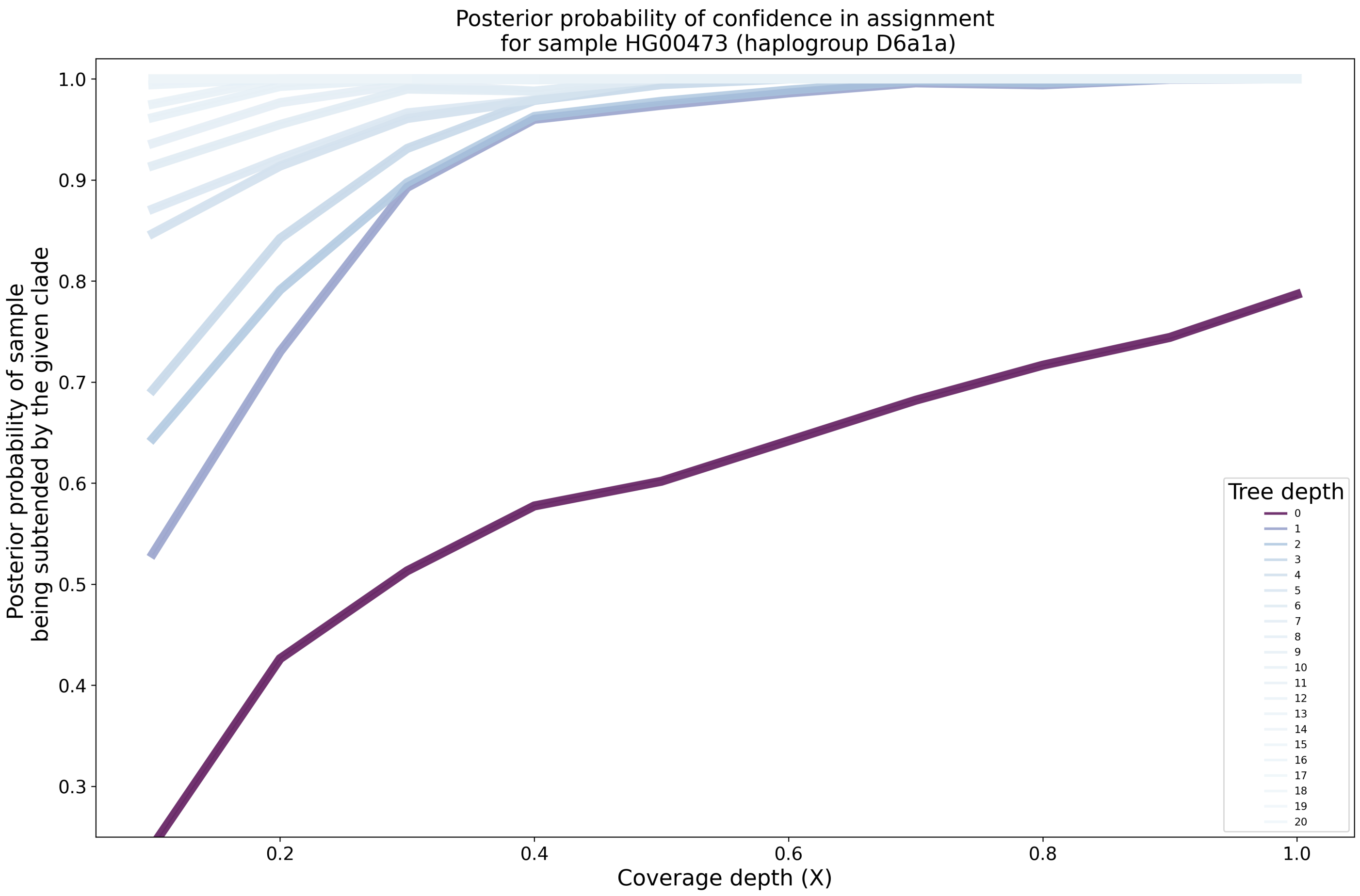

Supplementary Figure 54: **Clade-level posterior probabilities on empirical paired-end FASTQ data.** Each lineplot represents the mean over replicates at a fixed depth on the mitochondrial tree. The darker the line, the more basal the haplogroups.

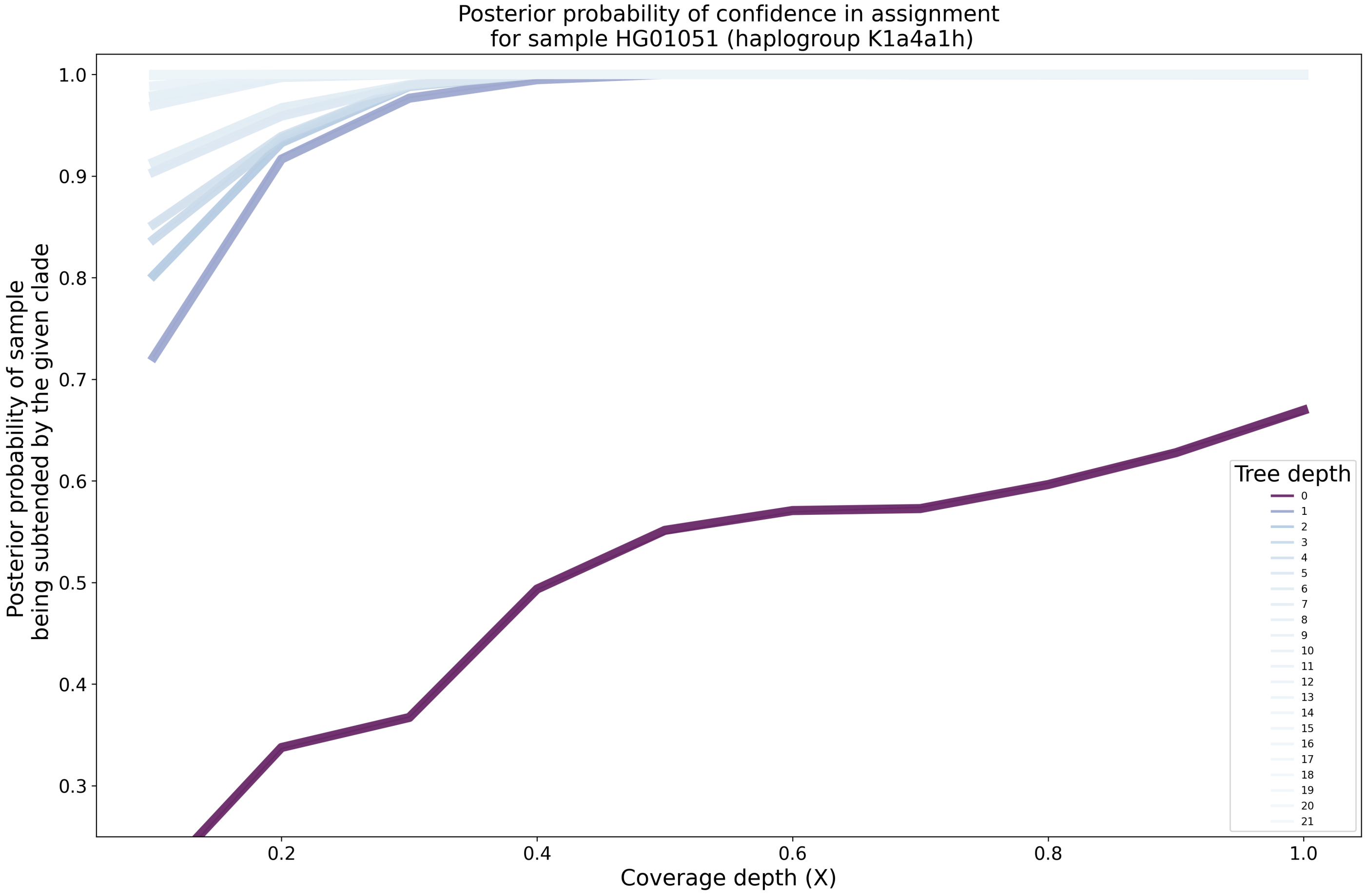

Supplementary Figure 55: **Clade-level posterior probabilities on empirical paired-end FASTQ data (CONTINUED).** Each lineplot represents the mean over replicates at a fixed depth on the mitochondrial tree. The darker the line, the more basal the haplogroups.

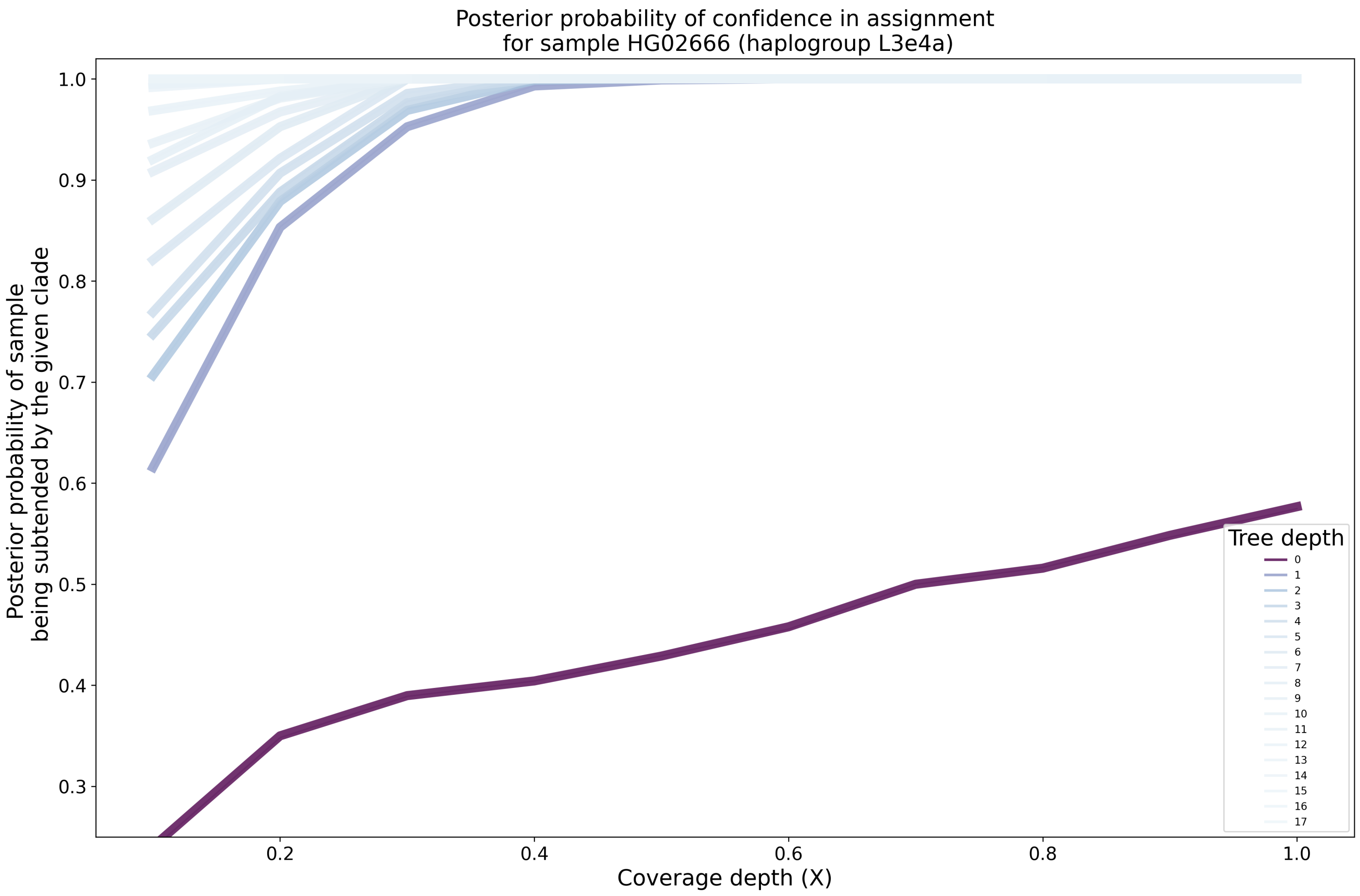

Supplementary Figure 56: **Clade-level posterior probabilities on empirical paired-end FASTQ data.** Each lineplot represents the mean over replicates at a fixed depth on the mitochondrial tree. The darker the line, the more basal the haplogroups.

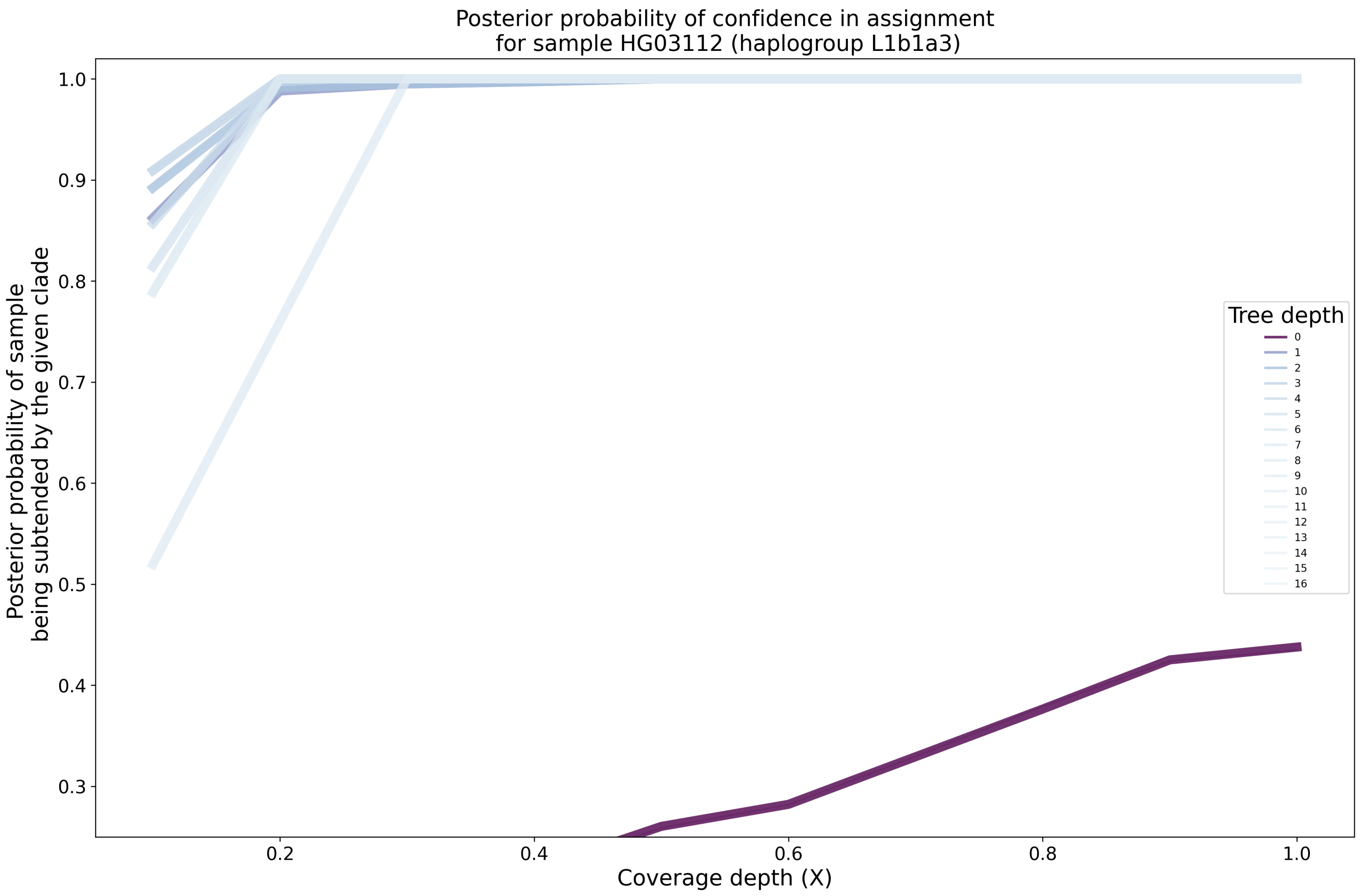

Supplementary Figure 57: **Clade-level posterior probabilities on empirical paired-end FASTQ data (CONTINUED).** Each lineplot represents the mean over replicates at a fixed depth on the mitochondrial tree. The darker the line, the more basal the haplogroups.

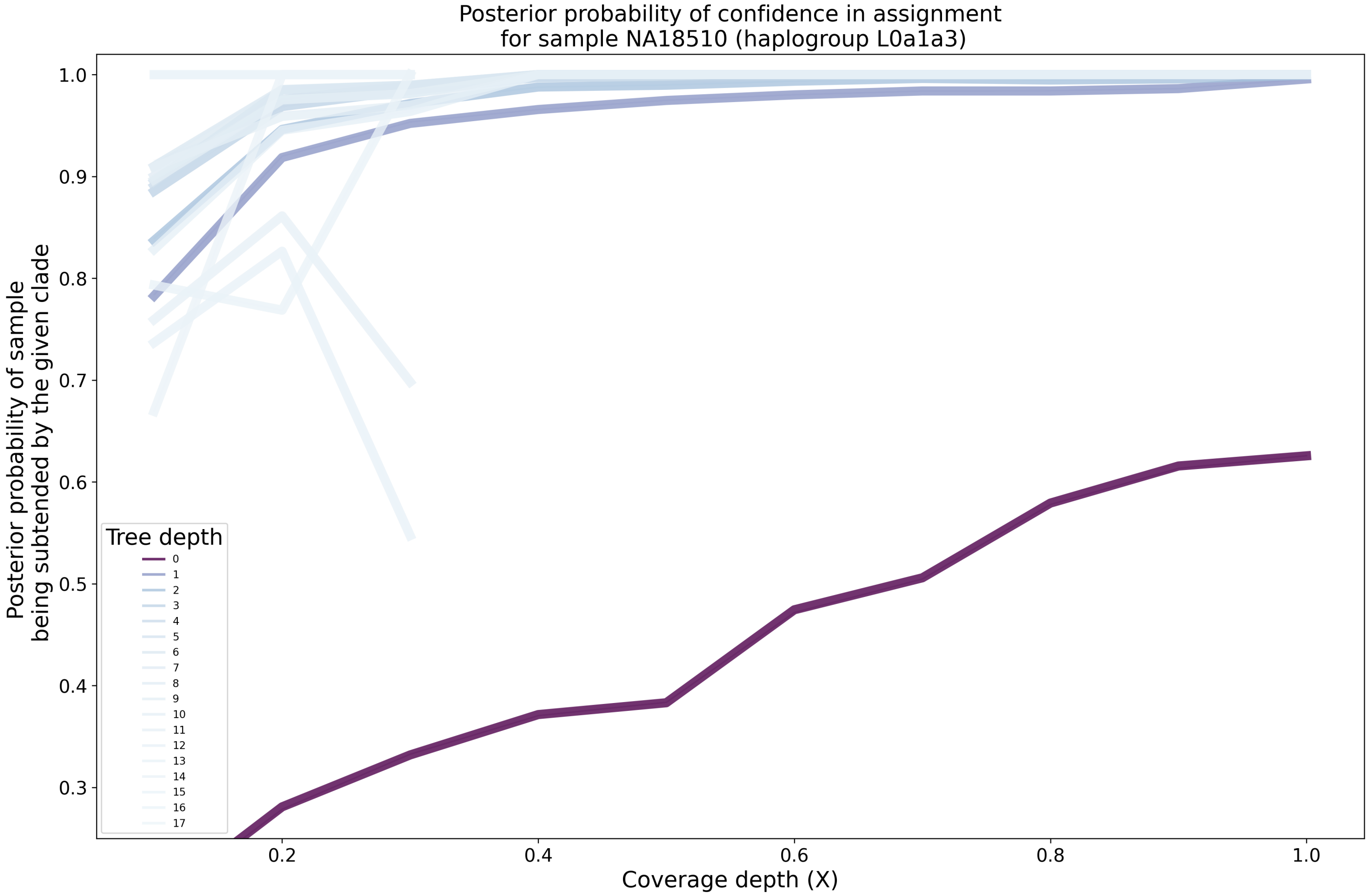

Supplementary Figure 58: **Clade-level posterior probabilities on empirical paired-end FASTQ data.** Each lineplot represents the mean over replicates at a fixed depth on the mitochondrial tree. The darker the line, the more basal the haplogroups.

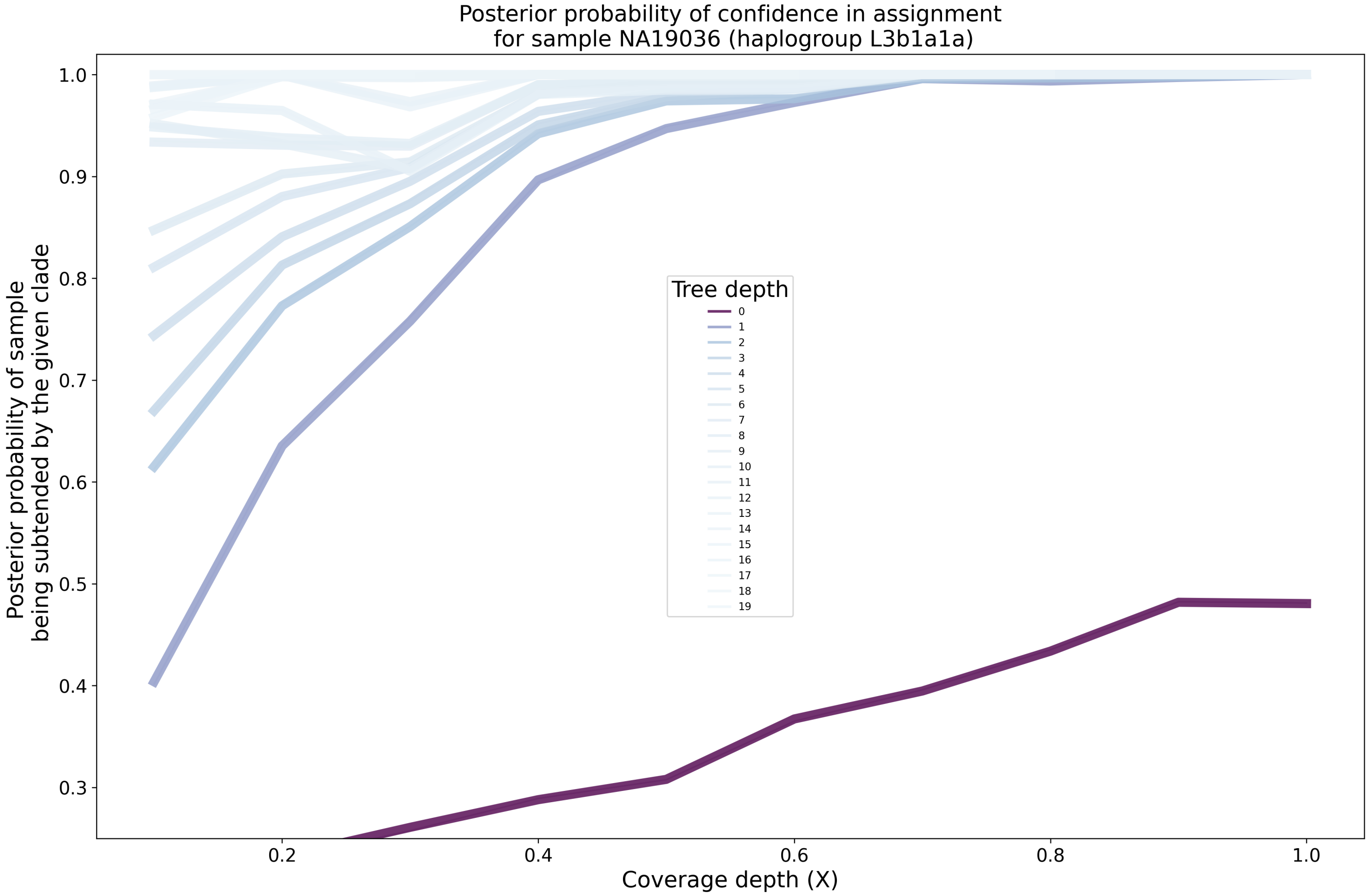

Supplementary Figure 59: **Clade-level posterior probabilities on empirical paired-end FASTQ data (CONTINUED).** Each lineplot represents the mean over replicates at a fixed depth on the mitochondrial tree. The darker the line, the more basal the haplogroups.

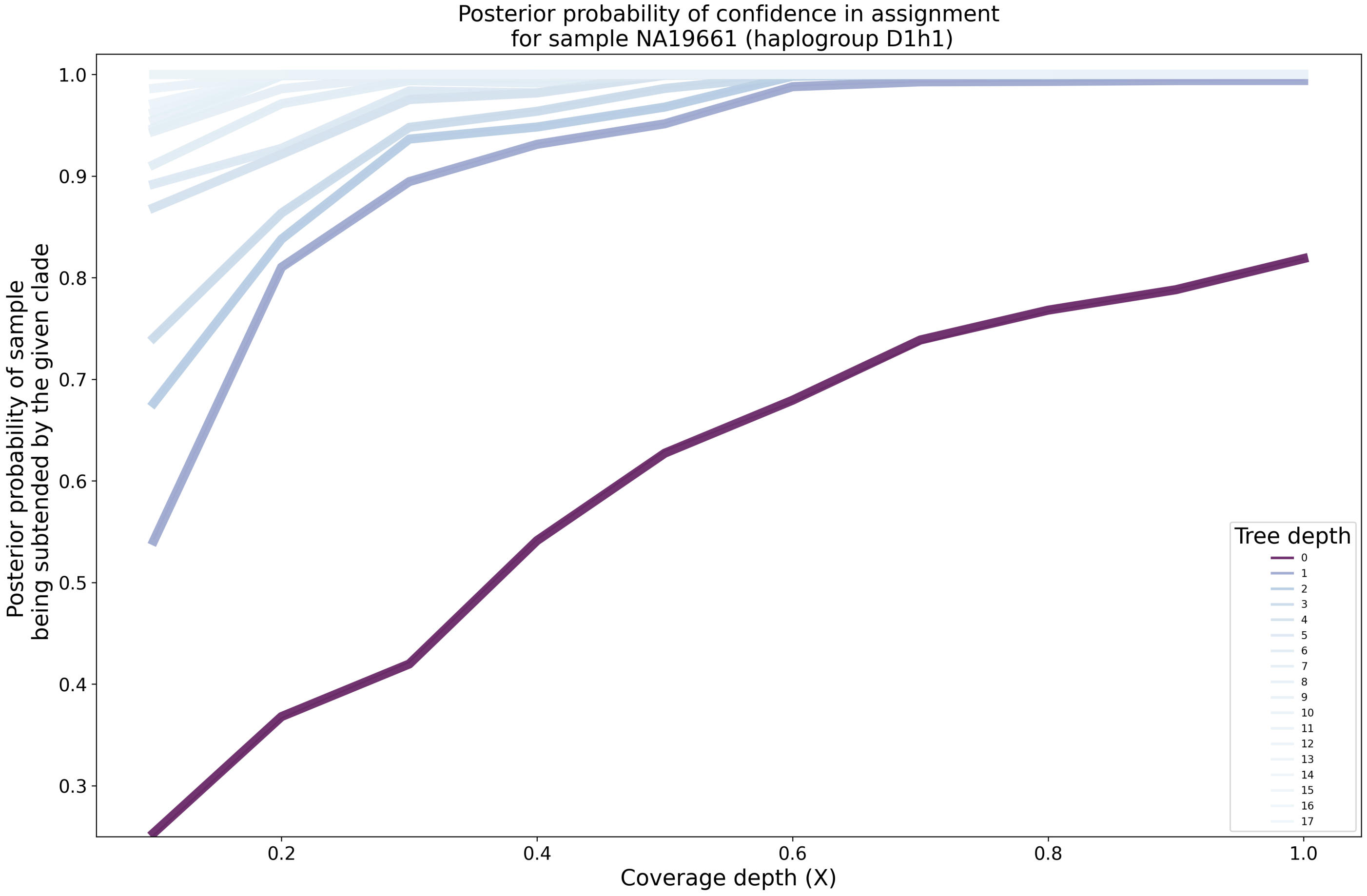

Supplementary Figure 60: **Clade-level posterior probabilities on empirical paired-end FASTQ data.** Each lineplot represents the mean over replicates at a fixed depth on the mitochondrial tree. The darker the line, the more basal the haplogroups.

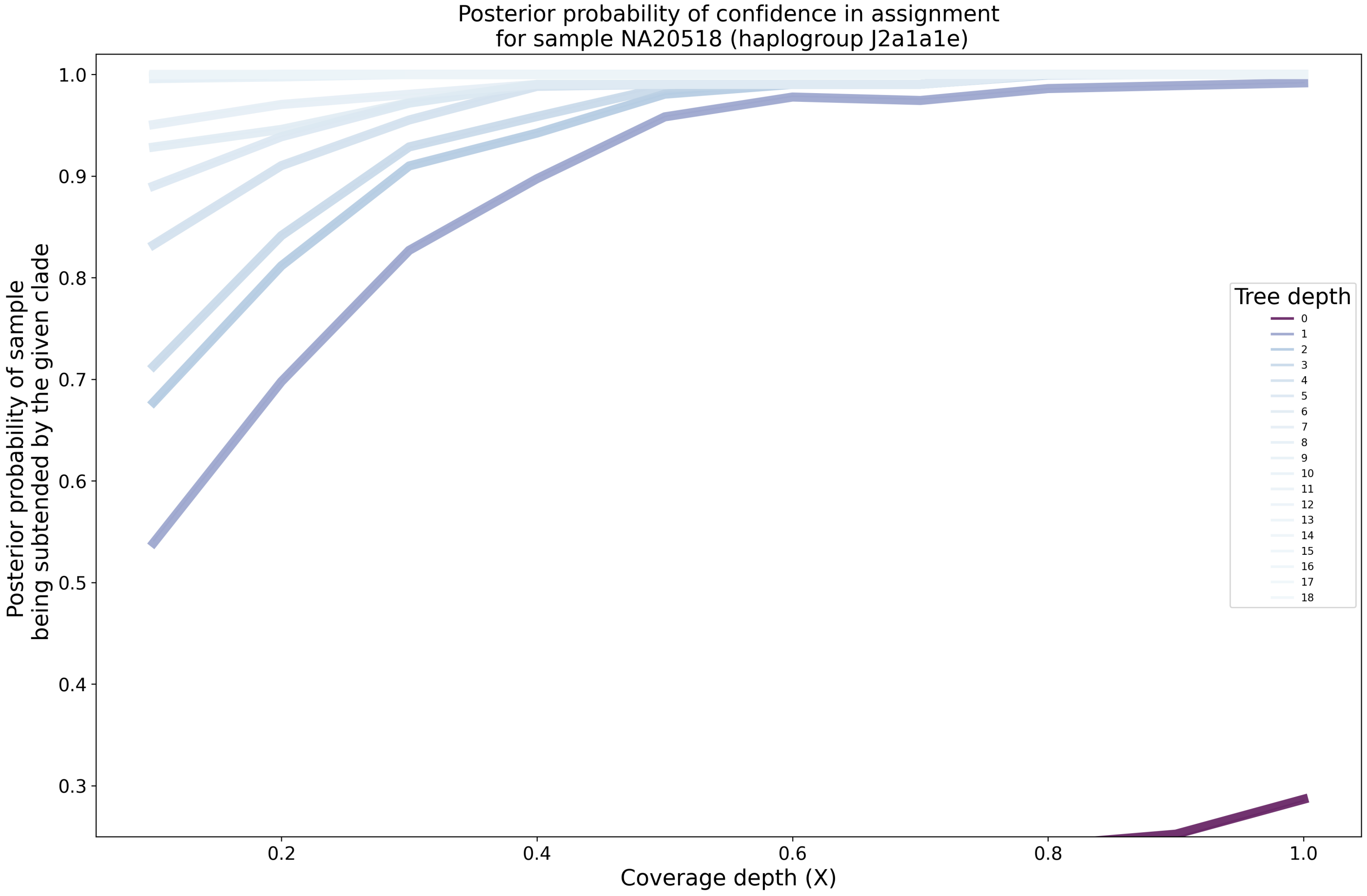

Supplementary Figure 61: **Clade-level posterior probabilities on empirical paired-end FASTQ data (CONTINUED).** Each lineplot represents the mean over replicates at a fixed depth on the mitochondrial tree. The darker the line, the more basal the haplogroups.

#### 2.5 Runtime and Peak Memory Usage

| Program | User time (s) | Wall clock time (s) | Peak memory usage (Mb) |
|---|---|---|---|
| HaploCart | 24.63 | 28.06 | 1334 |
| HaploGrep2 | 4.22 | 1.49 | 191 |
| Phy-Mer | 29.30 | 30.16 | 579 |

Supplementary Table 4: **Runtime and Peak Memory Usage on FASTA input.** Results are averaged over three input samples (NCBI accessions MZ387838, MW057682, MN894713). Statistics were measured with the command /usr/bin/time -v. User time and wall clock time are reported to two significant digits, while peak memory usage is rounded to the nearest integer. In all cases HaploCart was run in quiet mode (-q) without posterior calculations (-np).

| Program | User time (s) | Wall clock time (s) | Peak memory usage (Mb) |
|---|---|---|---|
| HaploCart | 25.40 | 11.14 | 1335 |
| HaploGrep2 | 4.22 | 1.49 | 191 |
| Phy-Mer | 29.30 | 30.16 | 579 |

Supplementary Table 5: **Runtime and Peak Memory Usage on FASTA input.** Results are averaged over three input samples (NCBI accessions MZ387838, MW057682, MN894713). Statistics were measured with the command /usr/bin/time -v. User time and wall clock time are reported to two significant digits, while peak memory usage is rounded to the nearest integer. In all cases HaploCart was run in quiet mode (-q) without posterior calculations (-np).

| Program | User time (s) | Wall clock time (s) | Peak memory usage (Mb) |
|---|---|---|---|
| HaploCart | 25.29 | 8.03 | 1335 |
| HaploGrep2 | 4.22 | 1.49 | 191 |
| Phy-Mer | 29.30 | 30.16 | 579 |

Supplementary Table 6: **Runtime and Peak Memory Usage on FASTA input.** Results are averaged over three input samples (NCBI accessions MZ387838, MW057682, MN894713). Statistics were measured with the command /usr/bin/time -v. User time and wall clock time are reported to two significant digits, while peak memory usage is rounded to the nearest integer. In all cases HaploCart was run in quiet mode (-q) without posterior calculations (-np).

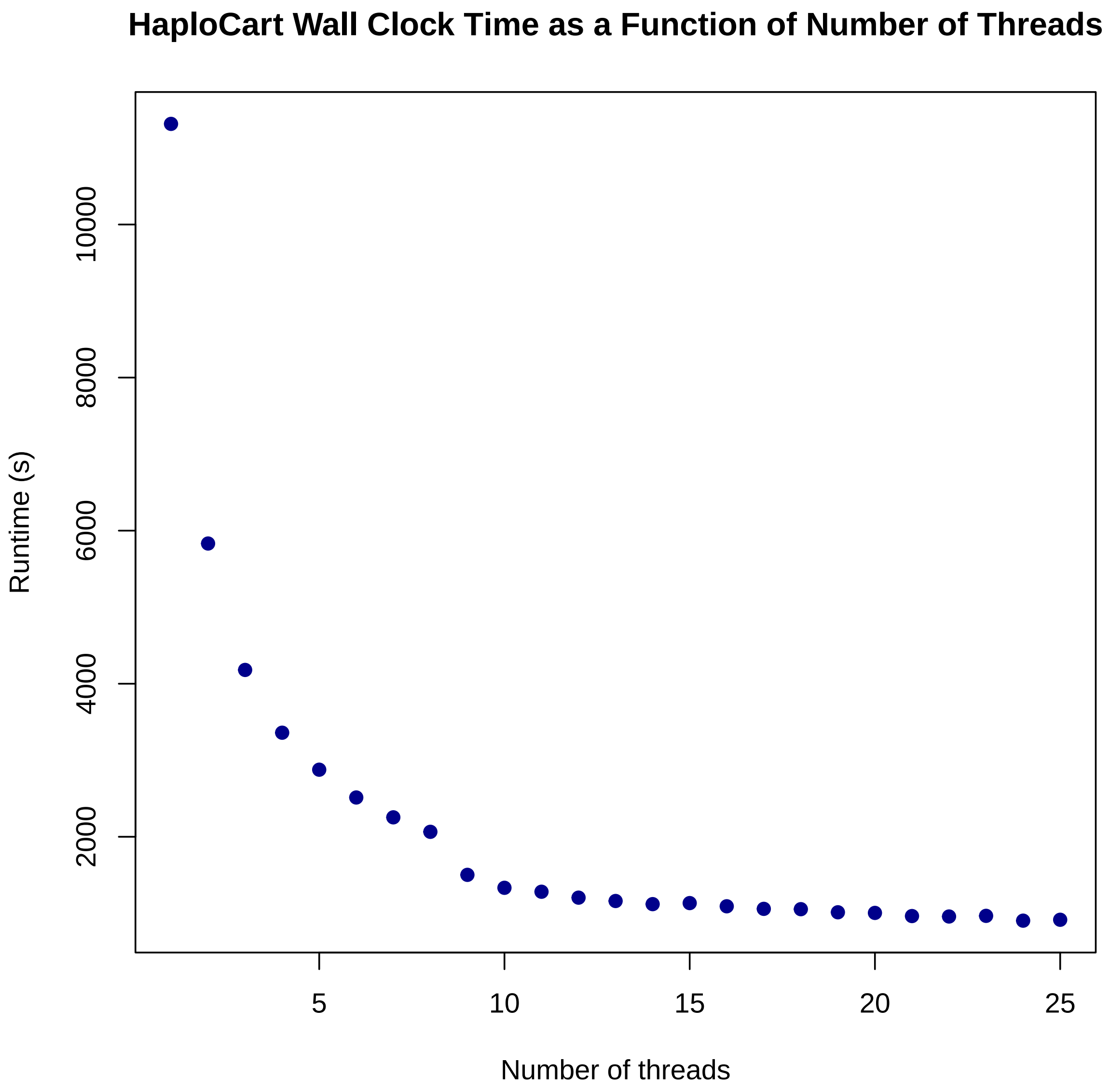

Supplementary Figure 62: `HaploCart` **Wall clock time (in seconds) as a function of number of threads from one to fifteen.** Each point represents the time taken to report haplogroup assignments on the 311 empirical consensus FASTA sequences detailed in Table 7. `HaploCart` was run in quiet mode (-q) without computing clade-level posterior probabilities of assignments (-np). Wall clock times were measured with the command `/usr/bin/time -v`.

#### 2.6 Empirical Data in FASTA Format

| Population | NCBI Accession | HaploCart prediction | HaploGrep2 prediction | |
|---|---|---|---|---|
| Australian | AF346963 | S1 | S1 | [7] |
| Australian | AF346964 | M42a | M42a | [7] |
| Australian | AF346965 | S2 | S2 | [7] |
| India | AF346966 | G3b1 | G3b1 | [7] |
| Bamileke | AF346967 | L3e3b2 | L3e3b2 | [7] |
| Biaka | AF346968 | L1c1a1a1a | L1c1a1a1a | [7] |
| Biaka | AF346969 | L1c1a2b | L1c1a2b | [7] |
| Buriat | AF346970 | C4b3b | C4b3b | [7] |
| Chukchi | AF346971 | A2b1 | A2b1 | [7] |
| Chinese | AF346972 | F4a1b | F4a1b | [7] |
| Chinese | AF346973 | M9a1a1a | M9a1a1a | [7] |
| Tatar | AF346974 | H5 | H5 | [7] |
| Dutch | AF346975 | H5a1a | H5a1a | [7] |
| Effik | AF346976 | L2a1i1 | L2a1i1 | [7] |
| Effik | AF346977 | L2a1a2 | L2a1a2 | [7] |
| English | AF346978 | HV0d | HV0+195 | [7] |
| Evenki | AF346979 | C4a2a1b | C4a2a1b | [7] |
| Ewondo | AF346980 | L3e1e1 | L3e1e1 | [7] |
| French | AF346981 | H1r | H1r | [7] |
| Georgian | AF346982 | T2e | T2e | [7] |
| German | AF346983 | J1c4 | J1c4 | [7] |
| Guarani | AF346984 | D1a2 | D1a2 | [7] |
| Hausa | AF346985 | L0a1a2 | L0a1a2 | [7] |
| Ibo | AF346986 | L1b1a3 | L1b1a3 | [7] |
| Ibo | AF346987 | L1c1d1 | L1c1d1 | [7] |
| Italian | AF346988 | U5b3a1a | U5b3a1a | [7] |
| Japan | AF346989 | D4b2b1 | D4b2b1 | [7] |
| Japan | AF346990 | D4a1 | D4a1 | [7] |
| Khirgiz | AF346991 | B4a1b1a | B4a1b1a | [7] |
| Kikuyu | AF346992 | L1c2a1a | L1c2a1a | [7] |
| Korea | AF346993 | B4a1b1a | B4a1b1a | [7] |
| Lisongo | AF346994 | L3e2b7 | L3e2b7 | [7] |
| Mandenka | AF346995 | L2c3a | L2c3a | [7] |
| Mbenzele | AF346996 | L1c1a2b | L1c1a2b | [7] |
| Mbenzele | AF346997 | L1c1a1a1a | L1c1a1a1a | [7] |
| Mbuti | AF346998 | L0a2b | L0a2b | [7] |
| Mbuti | AF346999 | L0a2b | L0a2b | [7] |
| Mkamba | AF347000 | L3h1a2a1 | L3h1a2a1 | [7] |
| Piman | AF347001 | B2a5 | B2a5 | [7] |
| Papua_New_Guinea_Coast | AF347002 | P1 | P1 | [7] |
| Papua_New_Guinea_Coast | AF347003 | Q1 | Q1 | [7] |
| Papua_New_Guinea_High | AF347004 | P1d1 | P1d1* | [7] |
| Papua_New_Guinea_High | AF347005 | P1 | P1+152 | [7] |
| Saami | AF347006 | V7a1 | V7a1 | [7] |
| Samoa | AF347007 | B4a1a1a16 | B4a1a1+152 | [7] |
| San | AF347008 | L0k1a1a | L0k1a1a | [7] |
| San | AF347009 | L0k1a1c | L0k1a1c | [7] |
| SibInuit | AF347010 | D2a1b | D2a1b | [7] |
| Uzbek | AF347011 | B4c2a | B4c2a | [7] |
| Warao | AF347012 | C1d1 | C1d1*5 | [7] |
| Warao | AF347013 | C1d1 | C1d1*5 | [7] |
| Yoruba | AF347014 | L3d1a1a | L3d1a1a | [7] |
| Yoruba | AF347015 | L3e2b1a1 | L3e2b1a1 | [7] |
| Mauritania | AF381981 | L2c1a | L2c1a*1 | [9] |
| Canary | AF381982 | U3a1c | U3a1c | [9] |
| Morocco | AF381983 | U3a | U3a | [9] |
| Morocco | AF381984 | M1a2a | M1a2a* | [9] |

| | | | | |
|---|---|---|---|---|
| Morocco | AF381985 | T2e1a | T | [9] |
| Morocco | AF381986 | X2b2 | X2b2 | [9] |
| Morocco | AF381987 | J1b2 | J1b2 | [9] |
| Morocco | AF381988 | L0a1b1 | L0a1b1 | [9] |
| Berber | AF381989 | U5b1b1 | U5b1b1+152 | [9] |
| Berber | AF381990 | V25 | V25 | [9] |
| Mauritania | AF381991 | L3b1a | L3b1a | [9] |
| Mauritania | AF381992 | L1c3a | L1c3a | [9] |
| Mauritania | AF381993 | H1 | H1 | [9] |
| Mauritania | AF381994 | L1b1a5 | L1b1a5 | [9] |
| Jordan | AF381995 | U2e | U2e | [9] |
| Jordan | AF381996 | M1b1 | M1b1 | [9] |
| Jordan | AF381997 | R | HV+73 | [9] |
| Jordan | AF381998 | L3d3b | L3d3b | [9] |
| Jordan | AF381999 | N1b1a | N1b1a | [9] |
| Jordan | AF382000 | U2d2 | U2d2 | [9] |
| Maragato | AF382001 | J1d1b | J1d1b | [9] |
| Maragato | AF382002 | H1a | H100 | [9] |
| Maragato | AF382003 | W1 | W1 | [9] |
| Leon | AF382004 | U2e1c1 | U2e1c1 | [9] |
| Leon | AF382005 | K1b2 | K1b2b | [9] |
| Leon | AF382006 | T1a1 | T1a1 | [9] |
| Leon | AF382007 | I5a1b | I5a1b | [9] |
| Morocco | AF382008 | U6a7a2 | U6a7a2 | [9] |
| Canary | AF382009 | C1b2 | C1b2 | [9] |
| Canary | AF382010 | A2 | A2 | [9] |
| Andalusia | AF382011 | U7b | U7b | [9] |
| Filipino | AF382012 | M7c1c3 | M7c1c3 | [9] |
| India | AF382013 | M30c | M30c | [9] |
| Taiwan_Aborigine | AJ842744 | B4a1a3a1a | B4a1a3a1a | [20] |
| Taiwan_Aborigine | AJ842745 | B4a1a2 | B4a1a2 | [20] |
| Taiwan_Aborigine | AJ842746 | B4a1a | B4a1a | [20] |
| Taiwan_Aborigine | AJ842747 | B4a1a4 | B4a1a4 | [20] |
| Taiwan_Aborigine | AJ842748 | B4a1a2 | B4a1a2 | [20] |
| Taiwan_Aborigine | AJ842749 | B4a1a | B4a1a | [20] |
| Taiwan_Aborigine | AJ842750 | B4a2a3 | B4a2a3 | [20] |
| Taiwan_Aborigine | AJ842751 | B4a2a1 | B4a2a1 | [20] |
| Caucasian | AY195745 | T2b21 | T2b21 | [13] |
| Caucasian | AY195746 | H3ak | H3ak | [13] |
| Caucasian | AY195747 | H5a1 | H5a1 | [13] |
| Native_American | AY195748 | D1 | D1 | [13] |
| Native_American | AY195749 | B2d | B2d | [13] |
| Caucasian | AY195750 | V | V | [13] |
| Caucasian | AY195751 | H11a7 | H11a7 | [13] |
| Caucasian | AY195752 | H3b1a | H3b1a | [13] |
| Evenki | AY195753 | C4a1a3a1 | C4a1a3a1 | [13] |
| Caucasian | AY195754 | J1c1d | J1c1d | [13] |
| Georgian | AY195755 | G2a1 | G2a1+16189 | [13] |
| Georgian | AY195756 | N1b1a3 | N1b1a3 | [13] |
| Iraqi_Israeli | AY195757 | H | H | [13] |
| Caucasian | AY195758 | H8c2 | H8c2 | [13] |
| Native_American | AY195759 | C1b1 | C1b1 | [13] |
| Korea | AY195760 | A5b1a | A5b1a* | [13] |
| Koryak | AY195761 | Z1a2a | Z1a2a | [13] |
| Koryak | AY195762 | G1b4 | G1b+16129 | [13] |
| Koryak | AY195763 | C4b2a | C4b2a | [13] |
| Caucasian | AY195764 | U2e1f1 | U2e1f1 | [13] |
| Caucasian | AY195765 | K1c1b | K1c1b | [13] |
| South_African | AY195766 | L2b1a3 | L2b1a3* | [13] |
| Caucasian | AY195767 | T2b1 | T2b1 | [13] |

| | | | | |
|---|---|---|---|---|
| Caucasian | AY195768 | W3a1 | W3a1 | [13] |
| Caucasian | AY195769 | I1b | I1b | [13] |
| Tofalar_Negidal | AY195770 | B4a1c2 | B4a1c2 | [13] |
| Taiwan | AY195771 | A5b1c1 | A5b1c1 | [13] |
| Udegei | AY195772 | C4b1 | C4b1 | [13] |
| Finland | AY195773 | X2c1 | X2c1 | [13] |
| Caucasian | AY195774 | J1c4b | J1c4b | [13] |
| Caucasian | AY195775 | H1b1c | H1b1+16362 | [13] |
| South_African | AY195776 | L2a1f | L2a1f | [13] |
| Khwe | AY195777 | L0d2a1a | L0d2a1a | [13] |
| Caucasian | AY195778 | J2b1a3 | J2b1a3 | [13] |
| Finland | AY195779 | W1a | W1a | [13] |
| South_African | AY195780 | L0a1a2 | L0a1a2 | [13] |
| Caucasian | AY195781 | V6 | V6 | [13] |
| South_African | AY195782 | L3d1d | L3d1d*2 | [13] |
| San | AY195783 | L1b1a4a | L1b1a4a | [13] |
| South_African | AY195784 | L3b1a | L3b1a*4 | [13] |
| South_African | AY195785 | L2c5 | L2c5 | [13] |
| Mixteca_Baja | AY195786 | A2v1 | A2v1 | [13] |
| Navajo | AY195787 | X2a2 | X2a2 | [13] |
| San | AY195788 | L2a2b1a | L2a2b1a | [13] |
| San | AY195789 | L1c1a1a1b1 | L1c1a1a1b1 | [13] |
| Nanaj_Negidal | AY195790 | D4o2a | D4o2a | [13] |
| Malay | AY195791 | F1a1a1 | F1a1a1 | [13] |
| Nanaj_Negidal | AY195792 | Y1a | Y1a+16189 | [13] |
| Berber | AY275527 | U6b | U6b | [10] |
| Canary | AY275528 | U6b1a1 | U6b1a1 | [10] |
| Senegal | AY275529 | U6b | U6b | [10] |
| Galicia | AY275530 | U6b | U6b | [10] |
| Mauritania | AY275531 | U6a7a1 | U6a7a1 | [10] |
| Maragato_Leon | AY275532 | U6a7a1 | U6a7a1 | [10] |
| Canary | AY275533 | U6a7b1 | U6a7b1 | [10] |
| Morocco | AY275534 | U6a1a1 | U6a1a1 | [10] |
| Mauritania | AY275535 | U6a | U6a+16189 | [10] |
| Berber | AY275536 | U6c2 | U6c2 | [10] |
| Canary | AY275537 | U6c1 | U6c1 | [10] |
| Australian | AY289051 | S2 | S2 | [6] |
| Australian | AY289052 | P3a | P3a | [6] |
| Australian | AY289053 | P6 | P6 | [6] |
| Australian | AY289054 | P7 | P7 | [6] |
| Australian | AY289055 | P6 | P6 | [6] |
| Australian | AY289056 | O1a | O1a | [6] |
| Australian | AY289057 | P4b1 | P4b1 | [6] |
| Australian | AY289058 | O1a | O1a | [6] |
| Australian | AY289059 | O | O | [6] |
| Australian | AY289060 | S2 | S2* | [6] |
| Australian | AY289061 | S2 | S2 | [6] |
| Australian | AY289062 | S4 | S4 | [6] |
| Australian | AY289063 | P5 | P5 | [6] |
| Australian | AY289064 | P4b | P4b | [6] |
| Australian | AY289065 | P3a | P3a | [6] |
| Australian | AY289066 | S3 | S3 | [6] |
| Australian | AY289067 | S3 | S3 | [6] |
| Cook | AY289068 | B4a1a1m1 | B4a1a1m1 | [6] |
| Cook | AY289069 | B4a1a1a16 | B4a1a1a*1 | [6] |
| Filipino | AY289070 | E2a | E2a | [6] |
| Kannada | AY289071 | M30b | M30b | [6] |
| Koraga | AY289072 | M30a1 | M30a1 | [6] |
| Koraga | AY289073 | U1a1a | U1a1a | [6] |
| Mullukurun | AY289074 | M35a1a | M35a1a | [6] |

| | | | | |
|---|---|---|---|---|
| Nasioi | AY289075 | Q1c1a | Q1c1a | [6] |
| Papua_New_Guinea_Coast | AY289076 | B4a1a | B4a1a | [6] |
| Papua_New_Guinea_Coast | AY289077 | B4a1a1 | B4a1a1 | [6] |
| Papua_New_Guinea_Coast | AY289078 | Q3a1 | Q3a1* | [6] |
| Papua_New_Guinea_Coast | AY289079 | Q3a | Q3a | [6] |
| Papua_New_Guinea_Coast | AY289080 | B4a1a1 | B4a1a1 | [6] |
| Papua_New_Guinea_Coast | AY289081 | Q1 | Q1 | [6] |
| Papua_New_Guinea_Coast | AY289082 | Q1a1a | Q1a1a | [6] |
| Papua_New_Guinea_Coast | AY289083 | B4a1a1a1 | B4a1a1a1 | [6] |
| Papua_New_Guinea_High | AY289084 | P2 | P2*1a | [6] |
| Papua_New_Guinea_High | AY289085 | Q1a | Q1a | [6] |
| Papua_New_Guinea_High | AY289086 | P1 | P1 | [6] |
| Papua_New_Guinea_High | AY289087 | P1d1 | P1d1* | [6] |
| Papua_New_Guinea_High | AY289088 | P2 | P2*1a | [6] |
| Papua_New_Guinea_High | AY289089 | Q3a1 | Q3a1* | [6] |
| Papua_New_Guinea_High | AY289090 | Q1 | Q1 | [6] |
| Papua_New_Guinea_High | AY289091 | P3b | P3b | [6] |
| Papua_New_Guinea_High | AY289092 | P1 | P1d | [6] |
| Samoa | AY289093 | B4a1a1a13 | B4a1a1a13 | [6] |
| Samoa | AY289094 | B4a1a1k1 | B4a1a1k1 | [6] |
| Taiwan_Aborigine | AY289095 | F4b1 | F4b1 | [6] |
| Taiwan_Aborigine | AY289096 | F4b1 | F4b1* | [6] |
| Taiwan_Aborigine | AY289097 | M7b1a2a1a | M7b1a2a1a* | [6] |
| Taiwan_Aborigine | AY289098 | M7b1a2a1 | M7b1a2a1 | [6] |
| Thai | AY289099 | F1a1a1 | F1a1a1 | [6] |
| Thai | AY289100 | B4c2a | B4c2a | [6] |
| Thai | AY289101 | B4c2 | B4c2 | [6] |
| Tonga | AY289102 | B4a1a1a16 | B4a1a1 | [6] |
| Buriat | AY519484 | B4d1’2’3 | B4d1’2’3 | [18] |
| Evenki | AY519485 | C4a2a1b | C4a2a1b | [18] |
| Ket | AY519486 | A8a | A8a | [18] |
| Koryak | AY519487 | C4b2a | C4b2a | [18] |
| Mansi | AY519488 | A12a | A12a | [18] |
| Negidal’tsy | AY519489 | B5b2a | B5b2a | [18] |
| Nganasan | AY519490 | C4b8a | C4b8a | [18] |
| Nganasan | AY519491 | D4o2a | D4o2a | [18] |
| Tofalar | AY519492 | B4a1c2 | B4a1c2 | [18] |
| Tofalar | AY519493 | Z1 | Z1 | [18] |
| Tubalar | AY519494 | B4b1a | B4b1a | [18] |
| Tuvan | AY519495 | B4a1c2 | B4a1c2 | [18] |
| Ul’chi | AY519496 | C1a | C1a | [18] |
| Ul’chi | AY519497 | M8a2b | M8a2b | [18] |
| Mansi | AY570524 | D5a3a1a | D5a3a1a | [18] |
| Tuvan | AY570525 | D5a2a1 | D5a2a1 | [18] |
| Tuvli | AY570526 | C4b1a | C4b1a | [18] |
| Nganasan | AY615359 | C5b1a1 | C5b1a1 | [18] |
| Tofalar | AY615360 | C4a1a3a1 | C4a1a3a1 | [18] |
| Ul’chi | AY615361 | C5a1 | C5a1 | [18] |
| Pakistan | AY882379 | U2a1a | U2a1a* | [1] |
| Pakistan | AY882380 | U2b2 | U2b2 | [1] |
| Pakistan | AY882381 | U2c1b | U2c1b*1a | [1] |
| Spain | AY882382 | U2e1a1 | U2e1a1 | [1] |
| Yemen | AY882383 | U3a2a1 | U3a2a1 | [1] |
| Adygei | AY882384 | U3b3 | U3b3 | [1] |
| Yemen | AY882385 | U3b1a1 | U3b1a1 | [1] |
| Spain | AY882386 | U4a1a | U4a1a | [1] |
| Italian | AY882387 | U4a2a | U4a2a | [1] |
| Adygei | AY882388 | U4b1a1a1 | U4b1a1a1 | [1] |
| Ethiopia | AY882389 | U9a | U9a | [1] |
| Pakistan | AY882390 | U9b1 | U9b1 | [1] |

| | | | | |
|---|---|---|---|---|
| Pakistan | AY882391 | U7b | U7b | [1] |
| Spain | AY882392 | U8a1a1 | U8a1a1 | [1] |
| Italian | AY882393 | U8b1a1 | U8b1a1* | [1] |
| Italian | AY882394 | K1c1a | K1c1a | [1] |
| Italian | AY882395 | K1a26 | K1a26 | [1] |
| Adygei | AY882396 | U1a1a | U1a1a+16129* | [1] |
| Italian | AY882397 | U1b3 | U1b3 | [1] |
| Adygei | AY882398 | U5a1f1a | U5a1f1a | [1] |
| Italian | AY882399 | U5a1 | U5a1* | [1] |
| Italian | AY882400 | U5b1b1d | U5b1b1+@16192 | [1] |
| Spain | AY882401 | U5b1b1d | U5b1b1d | [1] |
| Italian | AY882402 | U5b1b1d | U5b1b1d | [1] |
| Saami | AY882403 | U5b1b1a | U5b1b1a | [1] |
| Saami | AY882404 | U5b1b1a1 | U5b1b1a1 | [1] |
| Yakut | AY882405 | U5b1b1a | U5b1b1a | [1] |
| Saami | AY882406 | U5b1b1a3 | U5b1b1a3 | [1] |
| Fulbe | AY882407 | U5b1b1b | U5b1b1b | [1] |
| Berber | AY882408 | U5b1b1e | U5b1b1e | [1] |
| Italian | AY882409 | U5b1c | U5b1c | [1] |
| Italian | AY882410 | U5b1c1a1 | U5b1c1a1 | [1] |
| Italian | AY882411 | U5b1d1b | U5b1d1b | [1] |
| Berber | AY882412 | U5b1d1a | U5b1d1a | [1] |
| Italian | AY882413 | U5b2a1a2 | U5b2a1a2 | [1] |
| Spain | AY882414 | U5b2a1a2 | U5b2a1a2 | [1] |
| Italian | AY882415 | U5b2a2a1 | U5b2a2a1 | [1] |
| Ethiopia | AY882416 | U6a2a2a | U6a2a2a | [1] |
| Spain | AY882417 | U6b1 | U6b1 | [1] |
| Nicobarese | AY950286 | B5a1a1 | B5a1a1 | [19] |
| Nicobarese | AY950287 | B5a1a1 | B5a1a1 | [19] |
| Nicobarese | AY950288 | B5a1a1 | B5a1a1 | [19] |
| Nicobarese | AY950289 | F1a1a1 | F1a1a1 | [19] |
| Nicobarese | AY950290 | B5a1a1 | B5a1a1 | [19] |
| Onge | AY950291 | M32a | M32a | [19] |
| Onge | AY950292 | M32a | M32a | [19] |
| Onge | AY950293 | M31a1a | M31a1a | [19] |
| Onge | AY950294 | M31a1a | M31a1a | [19] |
| Onge | AY950295 | M32a | M32a | [19] |
| Great_Andamanese | AY950296 | M32a | M32a* | [19] |
| Great_Andamanese | AY950297 | M31a1a | M31a1a | [19] |
| Great_Andamanese | AY950298 | M31a1a | M31a1a | [19] |
| Great_Andamanese | AY950299 | M32a | M32a | [19] |
| Great_Andamanese | AY950300 | M31a1a | M31a1a | [19] |
| New_Britain | AY956412 | Q2a3a | Q2a3a | [3] |
| New_Ireland | AY956413 | Q2a | Q2a | [3] |
| New_Britain | AY956414 | Q2a | Q2a | [3] |
| Cambodian | AY963572 | F1a1a1 | F1a1a1 | [11] |
| Chinese | AY963573 | D4b2b | D4b2b | [11] |
| Bougainville | AY963574 | B4a1a1aa | B4a1a1aa | [11] |
| Chinese | AY963575 | A6a | A6a | [11] |
| Semang | AY963576 | M21a | M21a | [11] |
| Semang | AY963577 | M13b1 | M13b1 | [11] |
| Malay | AY963578 | N22a | N22a | [11] |
| Malay | AY963579 | R9b1a1a | R9b1a1a* | [11] |
| Malay | AY963580 | N21a | N21a | [11] |
| Malay | AY963581 | M21b1a | M21b1a* | [11] |
| Meyayu | AY963582 | M55 | M55 | [11] |
| Malayu | AY963583 | M22a | M22a | [11] |
| Semang | AY963584 | R21 | R21 | [11] |
| Ugandan | AY963585 | L0f2a1 | L0f2a1 | [11] |
| Italian | AY963586 | I3a1 | I3a1 | [2] |

| | | | | |
|---|---|---|---|---|
| New_Britain | DQ137398 | M28b | M28b | [12] |
| New_Britain | DQ137399 | M28b1 | M28b1 | [12] |
| New_Britain | DQ137400 | M28a7 | M28a7 | [12] |
| New_Britain | DQ137401 | M28a5b | M28a5b | [12] |
| New_Britain | DQ137402 | M27b1 | M27b1 | [12] |
| New_Britain | DQ137403 | M27b1 | M27b1 | [12] |
| New_Britain | DQ137404 | M27b1 | M27b1 | [12] |
| Bougainville | DQ137405 | M27c | M27c | [12] |
| New_Ireland | DQ137406 | M27c | M27c | [12] |
| New_Britain | DQ137407 | M29a | M29a | [12] |
| New_Britain | DQ137408 | M29a | M29a | [12] |
| New_Britain | DQ137409 | M29a | M29a | [12] |
| Bougainville | DQ137410 | M27a1a1 | M27a1a1 | [12] |
| Bougainville | DQ137411 | M27a1a1 | M27a1a1 | [12] |

Supplementary Table 7: Prediction of `HaploCart` and `HaploGrep2` on 311 human mitogenomes used in various publications including [4, 17, 15].
